# Mapping Human Survivability at Extreme Wet-Bulb Temperatures 32–35°C

**DOI:** 10.1101/2025.09.22.677706

**Authors:** Faming Wang, Haojian Wang, Yi Xu, Li Han, Xiong Shen, Yongchao Zhai, Xiang Zhang, Zhe Liang, Bo Hong, Ping Ma, Zhong Zhao

## Abstract

Graphical abstract

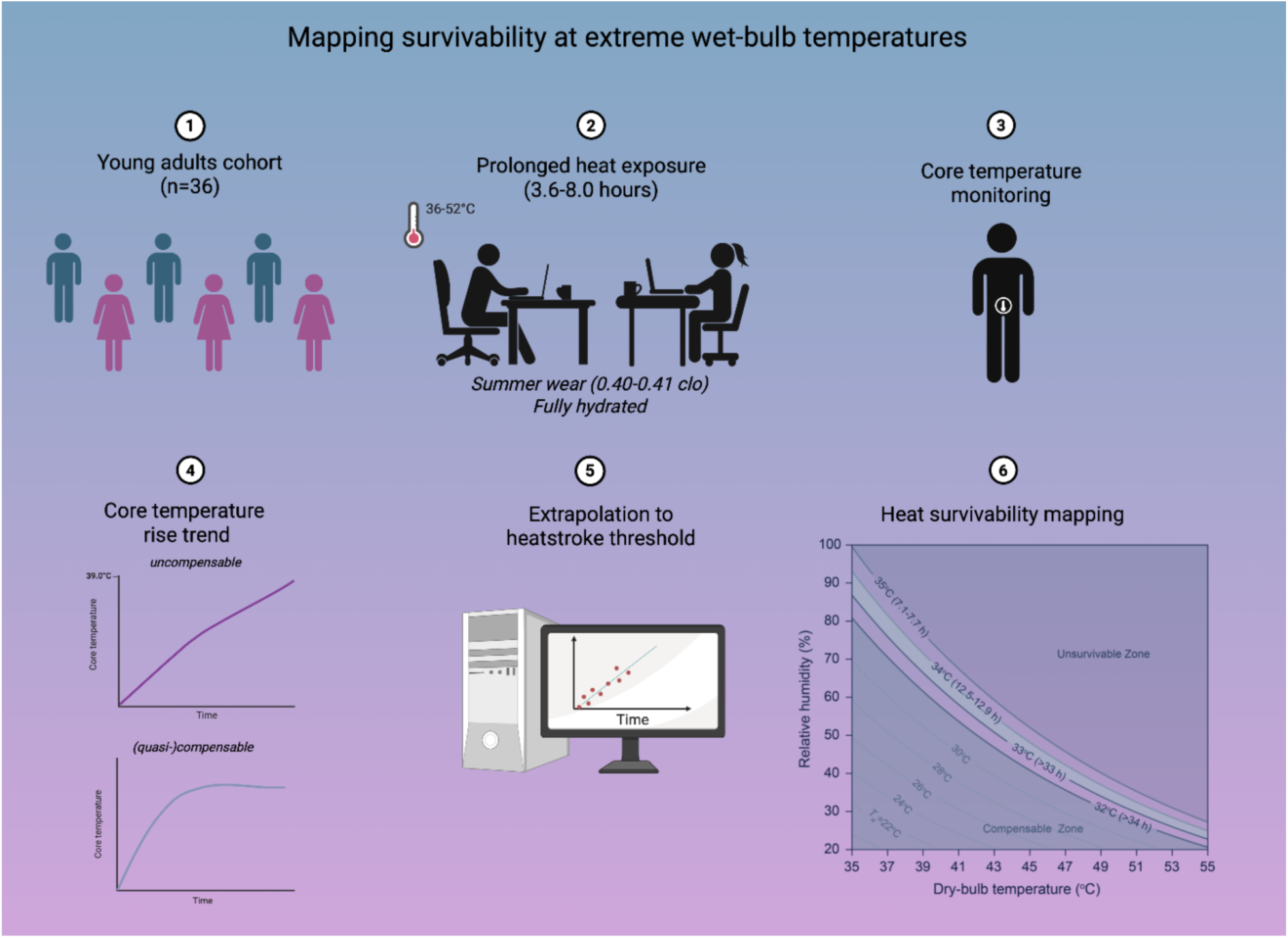

**Highlights:**

We report the first empirical human survival map across wet-bulb temperatures of 32–35°C.
HydraAon and behavior cooling extend survival beyond the 6-hour theoreAcal limit.
Females consistently tolerate extreme humid heat longer than males in controlled trials.
Our findings refine heat-health models and inform equitable climate adaptaAon strategies.

**In brief:** Extreme humid heat, measured by wet-bulb temperature (*T_w_*), is nearing the threshold of human survivability under climate change. In controlled trials with 36 adults, we provide the first empirical survival map across *T_w_*=32–35°C, showing that hydration and behavioral adaptations extend endurance beyond the theoretical 6-hour limit, with women tolerating heat longer than men. These findings refine human heat tolerance thresholds and inform equitable climate adaptation and heat-health strategies.

**SCIENCE FOR SOCIETY:** As climate change accelerates, extreme humid heat events—measured by wet-bulb temperature (*T_w_*)—pose a growing threat to human survival. This study provides the first empirical benchmarks of human survivability across *T_w_*=32–35°C, clarifying how long healthy adults can endure these conditions under full hydration and minimal activity. We find that while the theoretical 6-hour survival limit at *T_w_*=35°C is partly supported, both men and women can withstand these conditions for 7–8 hours when hydrated, with women demonstrating greater tolerance. At *T_w_*=34°C, projected heatstroke onset extends beyond 12–16 hours, while conditions at *T_w_*=32–33°C remain largely compensable for more than 30 hours. These results challenge prevailing heat tolerance models, underscore the role of behavioral and physiological adaptations, and provide critical data for designing equitable heat-health policies and early warning systems that account for sex differences and vulnerable populations.

Climate change-driven extreme heat events increasingly threaten human health. Here, we provide the first empirical map of human survivability across four extreme wet-bulb temperatures (*T_w_*=32– 35°C) in controlled trials. Thirty-six healthy young adults (20 males, 16 females) were exposed to twelve dry-bulb temperature and relative humidity combinations in shaded indoor settings under full hydration. Participants remained seated and performed light office tasks until core temperature (*T_core_*) reached 39°C or exposures lasted 8 hours. Clinical heatstroke threshold (*T_core_*=40.5°C) were projected by extrapolating from our direct *T_core_* measurements during controlled exposure trials. At *T_w_*=35°C, males and females reached heatstroke in approximately 7.1–7.7 and 8.3–8.6 hours, respectively—partly supporting the 6-hour theoretical limit while revealing enhanced short-duration resilience when hydrated. At *T_w_*=34°C, projected times are 12.5–12.9 hours (males) and 15.5–16.2 hours (females). Conditions at *T_w_*=32°C and 33°C remained (quasi-)compensable, with projected heatstroke onset after 33–35 hours. Females consistently tolerated heat stress longer than males. These physiological benchmarks are critical for improving heat-health models and embedding sex-specific vulnerability in equitable adaptation strategies amid accelerating global warming.

## INTRODUCTION

Anthropogenic climate change is intensifying the frequency, severity, and duration of heatwaves, posing mounting threats to human health, livelihoods, and ecosystems^1–4^. Since 2000, extreme heat has caused over 260,000 deaths globally, making it one of the deadliest weather-related hazards^5^. Projections are stark: under 1.5°C warming, 52% of those born in 2020 are expected to face unprecedented lifetime exposure to extreme heat, rising to 92% under 3.5°C by 2100^6^. These impacts disproportionately burden socioeconomically vulnerable regions—including the Middle East^7–9^, South Asia^10,11^, the North China Plain^12^, sub-Saharan Africa^13^, and Mexico^14^—where access to cooling, healthcare, and early warning systems remains limited. Recent city-level analyses of climate-health risks show that without adaptation, excess hospitalizations from extreme heat in China alone could reach over 5 million annually by 2100 under high-emission scenarios, underscoring the urgent need for locally tailored mitigation strategies^15^.

As global temperatures climb, wet-bulb temperatures (*T_w_*)—a combined measure of heat and humidity—are approaching the theorized survivability threshold of 35°C, beyond which thermoregulation is expected to fail, risking fatal hyperthermia within six hours^9,13,16^. This benchmark underpins many climate-health risk models but stems largely from thermodynamic theory, not empirical human data^17^. In practice, physiological tolerance varies: prior studies report critical *T_w_* thresholds ranging from 24.1°C to 34.6°C, depending on environmental conditions and study design^13,18–21^. A recent review notes that thresholds for uncompensable or unsurvivable thresholds (*T_w_*=19–34 °C) have already been breached in some regions^5^. However, these estimates are primarily based on modeling or short-duration exposures, leaving true physiological limits under prolonged, well-hydrated conditions uncertain.

Current estimates of human heat tolerance rely heavily on biophysical simulations and short-duration laboratory studies^19–23^, each with key limitations. Most simulations assume idealized heat balance conditions and are rarely validated above wet-bulb temperatures of 32°C^24,25^. They also neglect behavioral thermoregulation strategies that improve heat dissipation, especially in humid environments where evaporative cooling is constrained^26^. Notably, behaviors such as periodical sweat wiping^27^ and posture adjustments^28,29^ enhance evaporative, convective, or radiative heat loss, yet are often excluded from models or short-duration trials. These methodological gaps obscure the true limits of human heat tolerance and hinder the development of robust, evidence-based early warning systems and adaptation strategies.

To address this, we tested whether identical *T_w_* values induce consistent core temperature (*T_core_*) responses across varying combinations of dry-bulb temperatures (*T_db_*) and humidity. We further evaluated whether the *T_w_*=35°C threshold marks a definitive upper limit for human heat survivability. A total of 36 healthy young adults (20 males, 16 females) were exposed to 12 controlled humid heat conditions (*T_w_*=32–35°C) for up to 3.6–8.0 hours, with continuous *T_core_* monitoring. By extrapolating post-exposure core temperatures to the clinical heatstroke threshold (*T_core_*=40.5°C; Figs. 1a–c)^30,31^, we derived high-resolution survival estimates under extreme heat. These findings refine human heat tolerance limits and introduce actionable metrics for climate-health forecasting, with implications for early warning systems, hydration strategies, and equitable adaptation—especially in regions nearing human heat survivability thresholds.

**Fig. 1.**
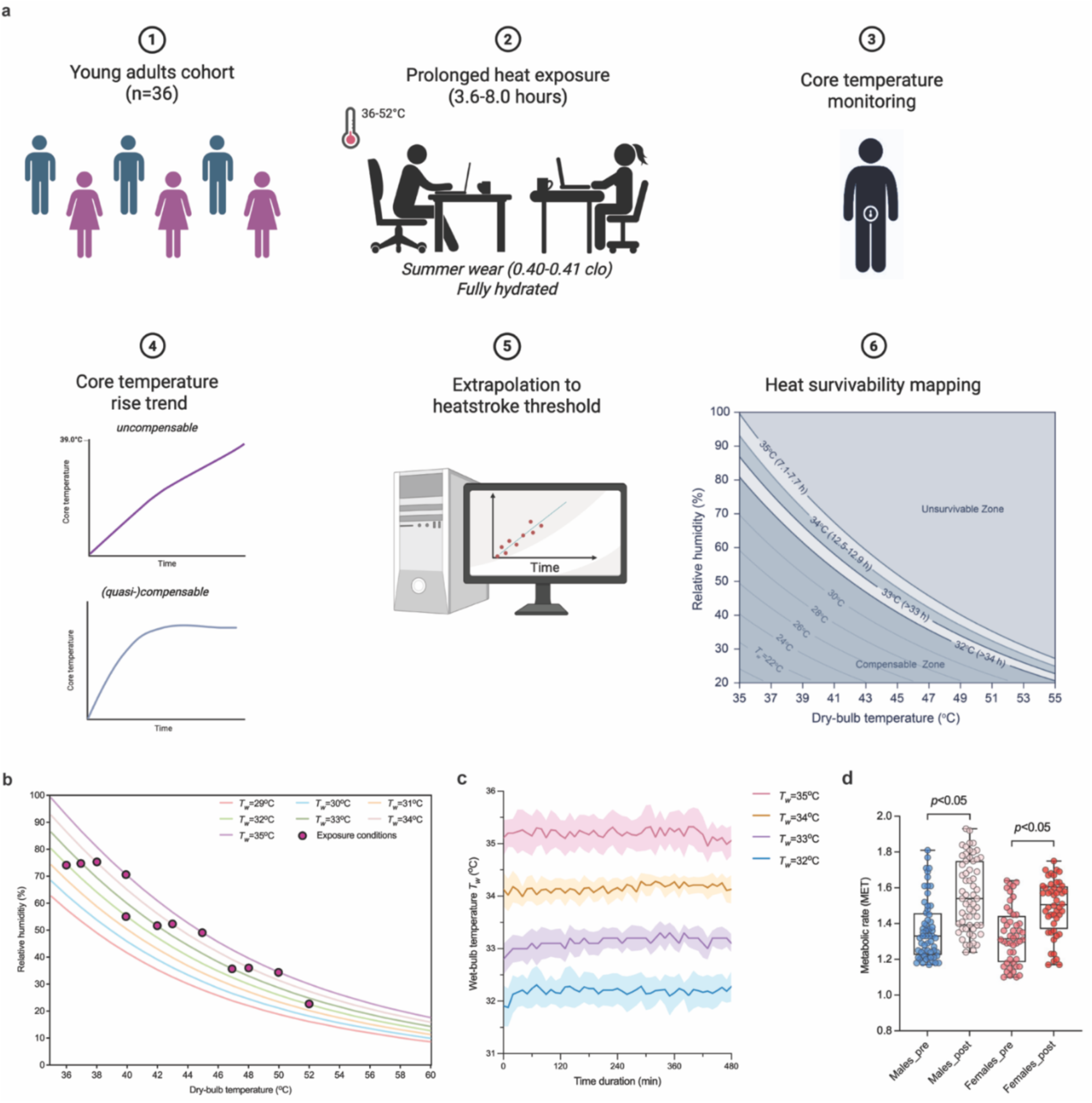
Overview of the experimental design, selected environmental conditions for prolonged heat exposure, and metabolic rate measurements. **a**, schematic of the experimental workflow used in this study. **b**, the twelve selected heat exposure conditions spanning *T_w_*=32–35°C. **c**, temporal variations of actual wet-bulb temperatures observed in the controlled indoor test chamber. **d**, metabolic rates of the young males (n=60) and females (n=48) measured at the beginning (11th– 20th minute) and end (471th–480th min) of the prolonged heat exposure under *T_w_*=33°C conditions. The final 5 minutes of each measurement period were used for analysis.

## RESULTS

### Metabolic rate and hydration

Baseline metabolic rates were modestly but significantly higher in males (males: 1.40±0.24 METs) compared to females (1.33±0.15 METs; *p*=0.010; Fig. 1d). Following 8-hour heat exposure, metabolic rates increased significantly in both sexes, rising to 1.63±0.31 METs in males (+16.7%; *p*<0.001) and 1.49±0.16 METs in females (+10.8%; *p*<0.001). Post-exposure metabolic rates remained significantly greater in males than females (*p*=0.013). Urine specific gravity (USG) measurements prior to exposure ranged narrowly between 1.008–1.016 in males and 1.008–1.015 in females, with post-exposure values maintained within a tight range (1.004–1.020; Supplementary Table S3). All USG values below the clinical threshold for euhydration (<1.020), confirming adequate hydration was preserved throughout the trials.

### Core temperature under matched *T_w_* conditions

Core temperature (*T_core_*) increased progressively at all tested wet-bulb temperature (*T_w_*) levels (i.e., 32°C, 33°C, 34°C, and 35°C), independent of the specific dry-bulb temperature (*T_db_*) and relative humidity (RH) combinations used to achieved those *T_w_* values (Fig. 2, Supplementary Fig. S1). Baseline *T_core_* values ranged from 36.7–37.0°C and increased to approximately 37.8°C, 37.8–38.0°C, 38.7–39.1°C, and 38.8–38.9°C at *T_w_*=32°C, 33°C, 34°C, and 35°C, respectively. The rate of *T_core_* rise, assessed via slope analyses, was consistent within each *T_w_* level, irrespective of the *T_db_*-RH combination. Although some visual variations were observed, statistically significant differences were rare and limited to comparisons between *T_db_*=40°C and 52°C, occurring in males between 70–200 minutes and in females between 365–480 minutes (all *p*<0.05), with relative *T_core_* differences of 0.16–0.27°C. Interindividual variability remained modest, with standard deviations spanning 0.13–0.28°C and rising to 0.3–0.4°C at higher *T_w_* levels.

**Fig. 2.**
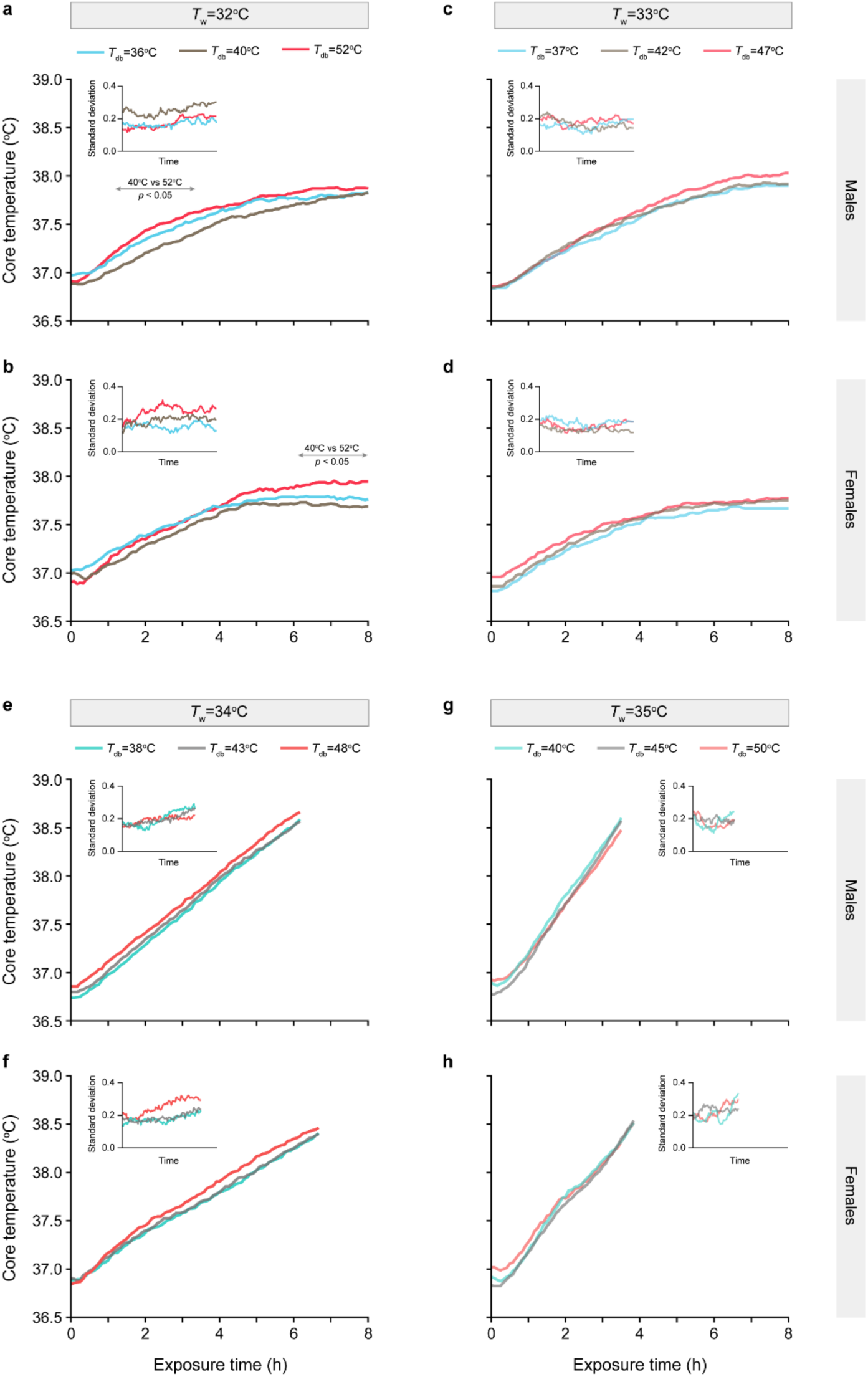
Core temperature (*T_core_*) responses under identical wet-bulb temperatures (*T_w_*). Core temperature (*T_core_*) responses are shown across different dry-bulb temperatures (*T_db_*) for *T_w_* values of 32°C (**a, b**), 33°C (**c, d**), 34°C (**e, f**), and 35°C (**g, h**), separated by sex (males: top panels; females: bottom panels). Insets display standard deviations. All *T_w_* levels induced consistent *T_core_* elevations across corresponding *T_db_* conditions, confirming the physiological equivalence of these matched *T_w_* environments.

Despite minor variations, *T_core_* trajectories largely overlapped across three different *T_db_*-RH combinations within each *T_w_* level (Figs. 2a–h), confirming that identical *T_w_* elicits consistent core temperature responses. These data affirm *T_w_* as a robust, physiologically grounded metric of heat stress assessment and support its fundamental role in defining human heat survivability limits.

### Core temperature at marginal *T_w_* (32-33°C)

Under (quasi-)compensable conditions at *T_w_*=32°C and 33°C, *T_core_* exhibited a biphasic trajectory in both sexes (Fig. 3). During the initial four hours, *T_core_* increased steadily at rates of 0.17-0.21°C/h (Figs. 3a–d), followed by a pronounced deceleration to rates between 0.01– 0.09°C/h over the subsequent four hours. These later rates fall below the conventional compensability threshold of 0.10°C/h, indicating the attainment of a thermal steady-state wherein evaporative and convective (if any) heat losses sufficiently counterbalance internal body and environmental heat loads. This biphasic pattern was evident in both males and females, suggesting the establishment of thermal steady-state under prolonged exposure when hydration is maintained.

**Fig. 3.**
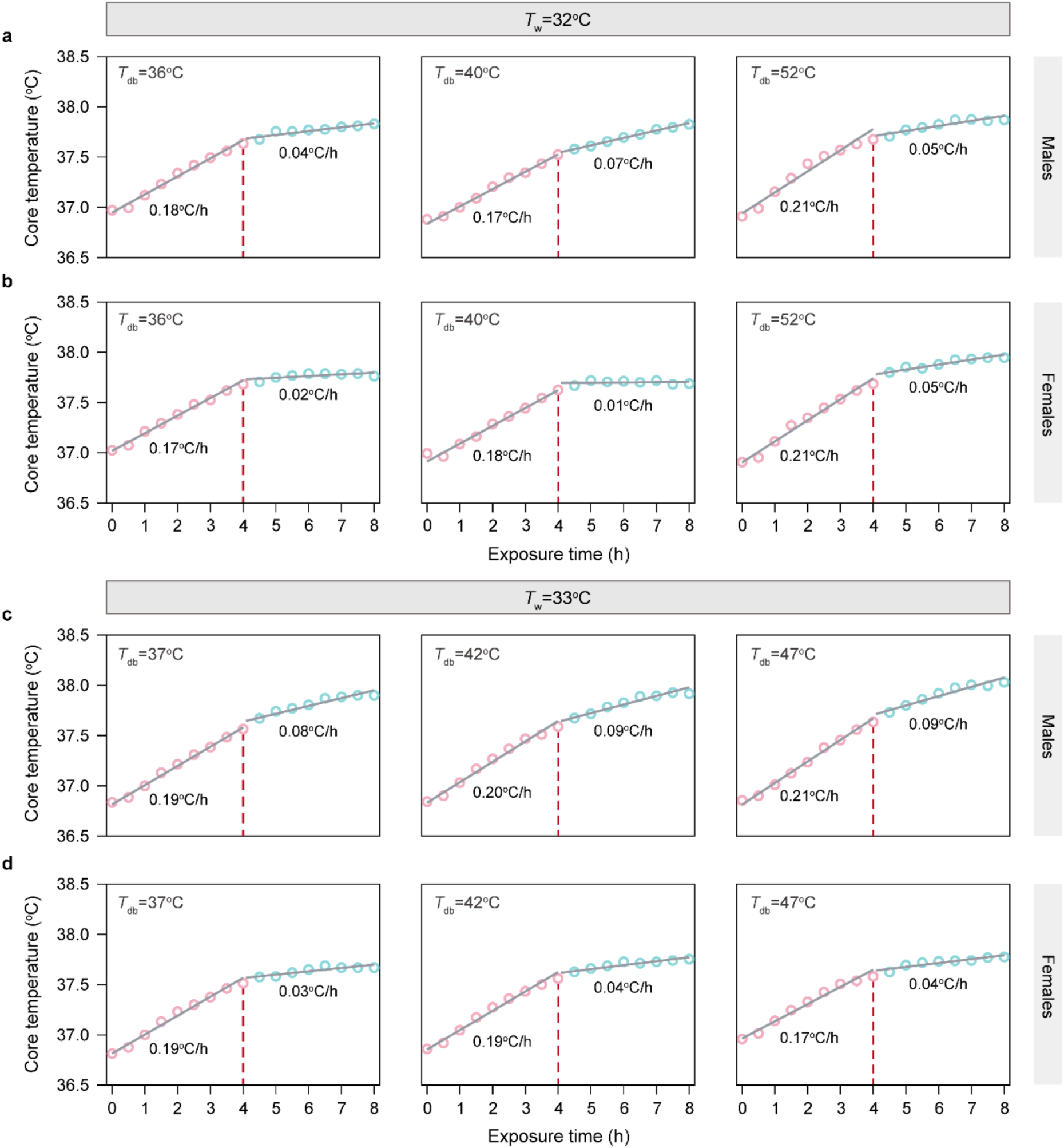
Biphasic core temperature rise trends at compensable temperatures of *T_w_*=32°C and 33°C. Core temperature (*T_core_*) trajectories in males (panels **a, c**) and females (panels **b, d**) show distinct biphasic patterns under varying dry-bulb temperatures (*T_db_*) at constant wet-bulb temperatures (*T_w_*). The initial phase reflects a rapid *T_core_* increase, followed by a plateau phase indicative of a steady-state where the rate of temperature rise falls below 0.10°C/h (dashed red lines denote transition points). Panels **a–b** show responses at *T_w_*=32°C; panels c–d depict *T_w_*=33°C. Transition times and corresponding *T_core_* rise rates are labeled for each condition. Results reveal sex-based differences in heat dissipation and steady-state onset timing, particularly at higher *T_db_* exposures.

Supporting this, time-to-steady-state analysis (Fig. 4) revealed that at *T_w_*=32°C, *T_core_* rise rates dropped below 0.10°C/h between 3.9–4.8 hours in males (Fig. 4a) and 4.0–4.9 hours in females (Fig. 4b). At *T_w_*=33°C, males reached steady-state later (5.4–5.8 hours; Fig. 4c) than females (3.8– 4.1 hours; Fig. 4d), suggesting potential sex-based differences in heat dissipation efficiency under these (quasi-)compensable conditions.

**Fig. 4.**
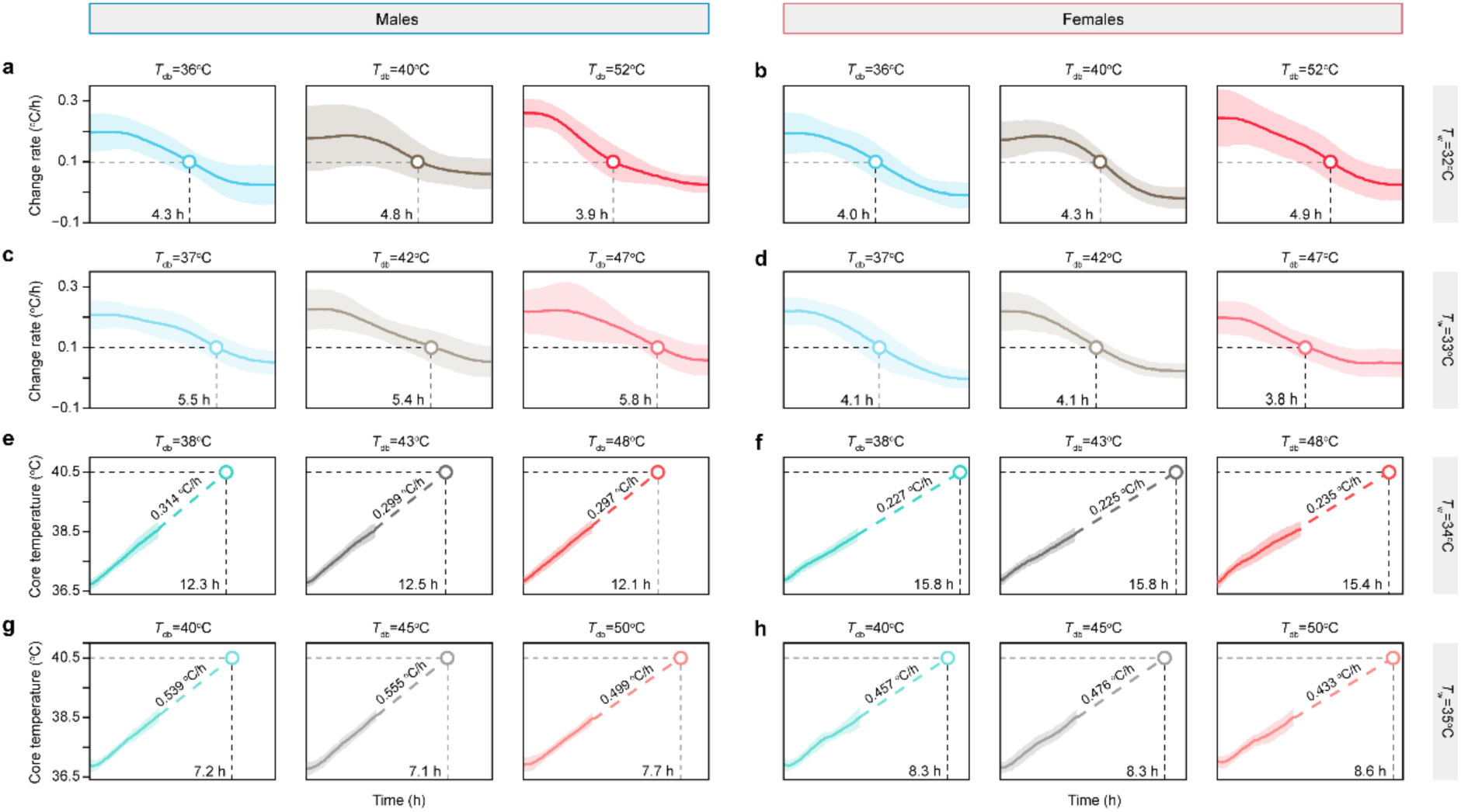
Core temperature (*T_core_*) change rates and time to thermal equilibrium or projected survival time under different wet-bulb temperature (*T_w_*) conditions in male (n=20) and female (n=16) participants. **a–d**, *T_core_* change rates (°C/h) and empirically determined time to reach thermal equilibrium under compensable heat stress conditions (*T_w_*=32°C and 33°C) in male (**a, c**) and female (**b, d**) participants, across three dry-bulb temperatures (*T_db_*) conditions per *T_w_* level. Dashed lines indicate the compensability threshold of 0.10°C/h, and circles denote the time point at which participants reached thermal equilibrium, as determined from observed *T_core_* trajectories over 8 hours of continuous exposure. **e–h**, *T_core_* rise rate and projected time to reach a core temperature threshold of 40.5°C under uncompensable conditions (*T_w_*=34°C and 35°C) in males (**e, g**) and females (**f, h**). Dashed lines represent extrapolated *T_core_* trajectory assuming a constant linear rise rate beyond the observed exposure period. Each panel represents the mean *T_core_* rise rate across three *T_db_*-RH combinations (for *T_w_*=34°C, *T_db_*=38°C & RH=75.1%, *T_db_*=43°C & RH=52.1%, and *T_db_*=48°C & RH=36.2%; for *T_w_*=35°C, *T_db_*=40°C & RH=70.3%, *T_db_*=45°C & RH=49.0%, and *T_db_*=50°C & RH=34.4%). Shaded areas represent ± one standard deviation (SD).

### Core temperature rise under uncompensable heat

In uncompensable heat (*T_w_*=34°C and 35°C), *T_core_* rose continuously without evidence of plateauing, exhibiting sex- and environment-specific variations (Figs. 2e-h). At *T_w_*=34°C, males exhibited *T_core_* rise rates of 0.314±0.063°C/h, 0.299±0.051°C/h, and 0.297±0.034°C/h at *T_db_* = 38°C, 43°C, and 48°C, respectively (Fig. 4e). Female demonstrated consistently lower rates of 0.227±0.036°C/h, 0.225±0.031°C/h, and 0.235±0.024°C/h (Fig. 4f). At *T_w_*=35°C, *T_core_* rise rates increased sharply, with males reaching 0.539±0.076°C/h, 0.555±0.059°C/h, and 0.499±0.072°C/h at *T_db_* = 40°C, 45°C, and 50°C, respectively (Fig. 4g). Female rates again remained lower: 0.457±0.087°C/h, 0.476±0.094°C/h, and 0.433±0.069°C/h (Fig. 4h).

Extrapolated times to reach the clinical heatstroke threshold (*T_core_*=40.5°C) reflected these sex disparities. At *T_w_*=34°C, males were projected to reach heatstroke in 12.6±2.2 hours, 12.9±2.1 hours, and 12.5±1.5 hours across the three increasing *T_db_* levels (Fig. 4e), whereas females required significantly longer durations: 16.2±2.1 hours, 16.2±2.2 hours, and 15.7±3.2 hours (Fig. 4f). At *T_w_*=35°C, male survival times declined to 7.2±0.9 hours, 7.1±0.7 hours, and 7.7±0.8 hours at *T_db_*=40°C, 45°C, and 50°C, respectively (Fig. 4g); females again demonstrated longer survival times of 8.5±1.8 hours, 8.3±1.7 hours, and 8.6±1.3 hours (Fig. 4h). These sex-based differences were statistically significant at all *T_db_* levels (Supplementary Fig. S2): 38°C (*p*<0.001), 40°C (*p*=0.002), 43°C (*p*<0.001), 45°C (*p*=0.004), 48°C (*p*<0.001), and 50°C (*p*=0.029), indicating greater resistance to core temperature elevation in females exposed to uncompensable heat.

To reach a core temperature of 40.5°C within six hours, a rise rate of 0.617°C/h would be necessary; this threshold was not exceeded under any tested condition. At *T_w_*=35°C, observed *T_core_* rise rates were 10.1–19.1% lower in males and 22.9–29.8% lower in females (Supplementary Fig. S1), indicating a greater physiological buffer against heat strain than predicted by thermodynamic theory. Nonetheless, the sustained upward trajectory of *T_core_* reflect severe thermal strain and highlights the high risk of life-threatening heat injury during prolonged exposure to extreme, uncompensable heat.

### Mapping human heat survivability

In males, estimated survival time decreased sharply with increasing wet-bulb temperature (*T_w_*): 7.1–7.7 hours at *T_w_*=35°C, 12.5–12.9 hours at 34°C; and >33–34 hours at 33°C and 32°C. Females consistently exhibited longer survival times: 8.3-8.6 hours at *T_w_*=35°C, 15.7–16.2 hours at 34°C, and >35 hours at 33°C and 32°C. These durations represent extrapolated times to reach *T_core_*=40.5°C, derived from observed *T_core_* trajectories up to approximately 39.0°C.

Sex-specific psychrometric charts (Figs. 5a-b) illustrate these survival thresholds across combinations of dry-bulb temperature and relative humidity. A critical transition emerged at *T_w_*=33°C and 34°C, beyond which survival time declined precipitously. Above *T_w_*≥34°C, conditions became uncompensable despite full hydration and the availability of behavioural thermoregulatory mechanisms such as sweat wiping and postural adjustments (Supplementary Video S1). Across all thresholds, females demonstrated consistently greater heat tolerance than males, underscoring the importance of sex as a factor in heat survivability under extreme humid conditions.

**Fig. 5.**
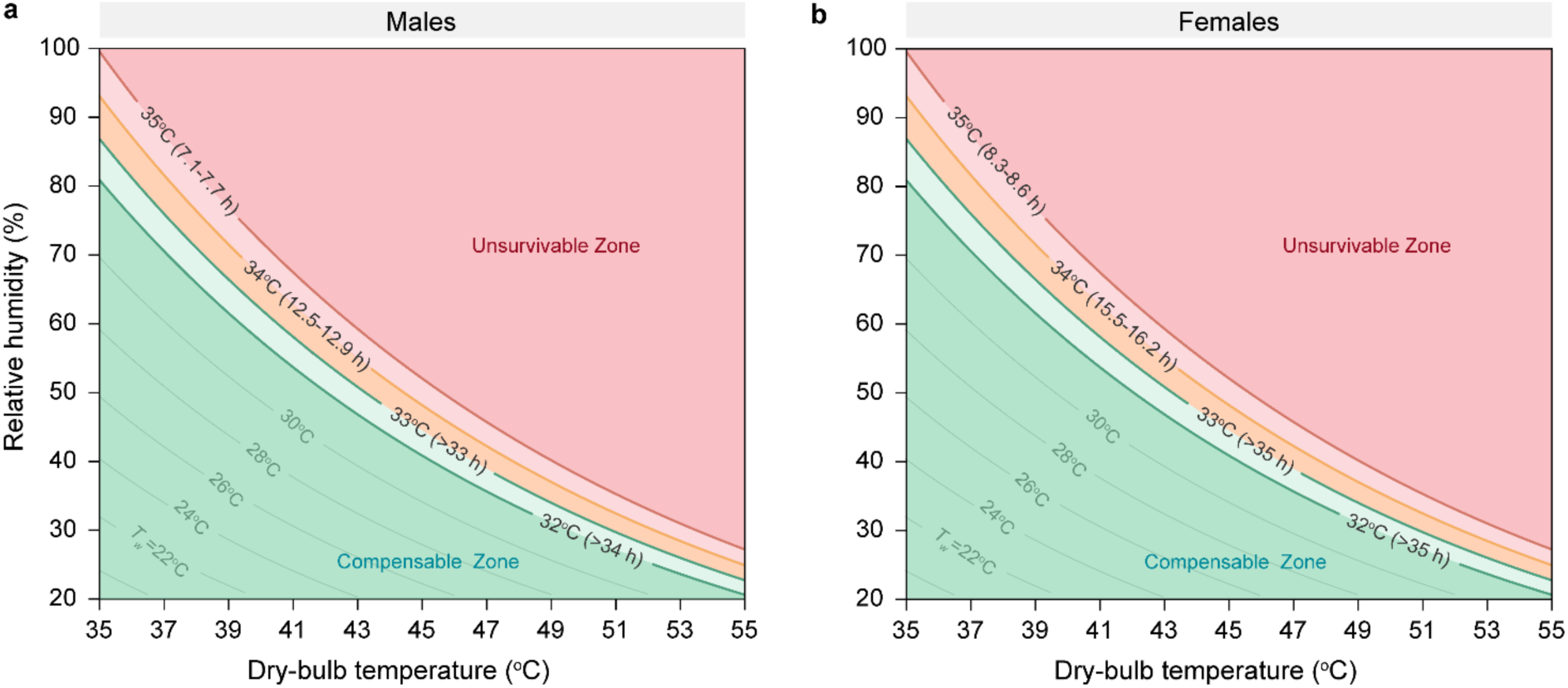
Heat stress survivability mapping for males and females. **a**, males (n=20). **b**, females (n=16). Survival time estimates were mapped onto sex-specific psychrometric charts (Figures 4a **and 4b**), delineating compensable (green zone) and uncompensable (red zone) heat stress zones across dry-bulb temperature and relative humidity. The critical transition (yellow zone) heat stress zone occurs at *T_w_*=33–34°C, beyond which survival times decrease sharply. At *T_w_* ≥34°C, both sexes enter the uncompensable zone, indicating a rapid decline in survivability.

## DISCUSSION

This study advances our understanding of human heat tolerance under extreme humid heat in three significant ways. First, it provides direct empirical evidence that identical wet-bulb temperature (*T_w_*) conditions elicit consistent core temperature (*T_core_*) responses across a wide range of temperature-humidity combinations. Despite variations in dry-bulb temperature and relative humidity (RH), Tcore trajectories remained nearly indistinguishable at each *T_w_* level. This uniformity affirms *T_w_* as a robust and physiologically grounded index of heat stress, reinforcing its relevance for climate-health surveillance and public warning systems.

Second, our data refine the widely cited *T_w_*=35°C survival limit. Previously considered an absolute upper limit for human viability, our prolonged exposure data suggest it functions more as an inflection point, where physiological risk escalates with time. Core temperatures rose steadily under all *T_w_*=35°C conditions; yet most participants, particularly females, did not reach clinical heatstroke thresholds within six hours. On average, the time required to reach a critical core temperature of 40.5°C was 7.1–7.7 hours in males and 8.3–8.6 hours in females. Notably, survival time decreased sharply by 38.4–48.8% as *T_w_* increased from 34°C to 35°C in both sexes, reinforcing the physiological significance of the *T_w_*=35°C benchmark. While conservative, this threshold remains meaningful, especially for prolonged exposures. The consistent observation of slower *T_core_* rise in females underscores the need to incorporate sex-based physiological differences in future models of heat vulnerability^32,33^.

Third, we present the first empirical maps of human heat survivability across *T_w_* values from 32°C to 35°C. By extrapolating time-to-overheat data and plotting survival durations on psychrometric charts, we delineate the transition from (quasi-)compensable to uncompensable heat stress. These sex-specific maps provide a time-resolved, physiologically grounded framework for estimating human survivability. In doing so, they move beyond abstract model assumptions and support targeted public health interventions in regions facing rising wet-bulb temperatures.

The *T_w_*=35°C threshold, originally proposed by Sherwood and Huber^14^, has shaped much of the heat risk modeling literature. However, critical environmental limits (CELs) reported in recent studies vary widely (*T_w_*=24.1–34.6°C), even under comparable metabolic loads^19–22,34–41^. These estimates often derived from short-duration, stepwise exposure protocols (with ramped increases in *T_db_* or RH) lasting ≤3 hours (Supplementary Table S1), with physiological inflection points identified within 5–10 minutes. Such designs may reflect delayed thermoeffector activation rather than true heat imbalance^24,25^, potentially misrepresenting critical physiological thresholds. Moreover, inter- and intra-individual variability is often not accounted for (Supplementary Fig. S3).

Our approach addresses these limitations directly. By maintaining stable environmental conditions and prolonged exposure durations (up to 8 hours), we demonstrate that *T_w_*=35°C induces uncompensable heat stress, characterized by a continuous rise in *T_core_* and no evidence of thermal equilibrium. Yet the rate of increase was gradual, and most participants, particularly females, did not reach *T_core_* ≥40.5°C within six hours. These time-resolved data challenge the notion of *T_w_*=35°C as an instantaneous survival limit, reframing it instead as a threshold beyond which physiological risk accumulates with duration. This temporal nuance is often missing from existing survivability limits and urban heat action plans, which tend to treat *T_w_* thresholds as binary endpoints.

Sex-based differences also emerged consistently. Females exhibited slower *T_core_* rise and longer times to high-risk thresholds under identical *T_w_* conditions. These findings likely reflect differences in body composition, metabolic rate, and evaporative efficiency^26,27,43,44^, and align with previous reports of greater heat resilience in females under passive humid heat stress^45,46^. Notably, most prior studies have either overlooked sex effects or pooled unpaired male and female data, contributing to the variability seen in reported CELs. Our findings, in line with a limited number of sex-stratified studies^34,36^, highlight the need for biological realistic modeling of human heat vulnerability^47^.

In comparing our data with biophysical and thermoregulatory models, notable discrepancies emerged. State-of-the-art frameworks—including those by Vanos et al.^19^ and Xu et al.^20^ —often assume fixed metabolic rates and constant sweating efficiency, and define heatstroke using static *T_core_* thresholds (39.5–43.0°C) without incorporating exposure duration or circadian variation^48^.

Validation studies are typically based on short-duration trials and rarely report uncertainty estimates. Few studies^21,35^, including our own, provide standard deviations or confidence intervals for *T_w_* and *T_core_*, which are essential for model calibration and evaluation.

Our findings suggest that such models may overstate the immediacy of risk at *T_w_*=35°C. For instance, Vanos et al.^19^ estimate survival durations of less than six hours at this level, whereas our participants exceeded this time—on average—by over an hour in males and more than two hours in females. These discrepancies highlight the need for models to incorporate dynamic, sex-specific core temperature trajectories and inter-individual variability. Future modeling efforts should move beyond threshold-based approaches to integrate time-dependent thermal dose metrics and delayed thermoregulatory feedbacks.

The public health implications of these refinements are significant. As climate change increases the frequency and intensity of extreme humid heat events^49–51^, decision-makers and public health authorities require evidence-based tools to assess risk. Our findings suggest that current *T_w_*-based alert systems could be improved by accounting for exposure duration and biological variability. In high-risk regions such as South Asia, the Arabian Gulf, and Northern China—where daily peak *T_w_* already approaches or exceeds 33–34°C^10–12^—understanding how long individuals can safely tolerate such conditions is critical. Vulnerable groups without access to cooling or hydration may still be at risk below sub-threshold *T_w_* levels, especially when additional stressors such as exertion, medication, or chronic disease are present. Our empirical maps establish physiologically grounded limits of human heat survivability during prolonged humid heat exposure, providing a replicable and high-resolution framework for defining survival limits that cannot be captured by short-duration inflection-point analysis or existing biophysical and thermoregulatory models.

Our study has direct implications for public health interventions, particularly in developing or tropical regions where access to cooling is limited. Current early-warning systems often rely on fixed temperature or wet-bulb thresholds (e.g., *T_w_*=35°C) as binary indicators of survivability. However, our data demonstrate that survivability is time-dependent and varies by sex, suggesting that duration of exposure must be integrated into heat alert protocols. For instance, a city experiencing *T_w_*=34°C for 12 continuous hours may pose similar physiological risks as a shorter but more intense *T_w_*=35°C event. Our sex-specific survival maps can support more granular, tailored alert systems—flagging risk earlier for vulnerable groups and guiding local authorities in planning adaptive measures such as staggered work hours, hydration checkpoints, or mandated breaks for outdoor laborers. Likewise, housing design standards, particularly in dense urban settings, may benefit from incorporating ventilation strategies or passive cooling thresholds tied to wet-bulb metrics. By embedding empirical survivability limits into public health infrastructure, climate adaptation can shift from reactive to preventive, especially in regions projected to face more frequent and severe humid heatwaves.

This study focused on healthy young adults during prolonged passive indoor exposure, thereby establishing a critical physiological baseline. Nonetheless, the results may not be generalizable to more vulnerable populations. Future work should expand to include older adults, children, individuals with chronic conditions, and populations in low-development countries to better reflect global variability in heat tolerance^52,53^. Additionally, we did not investigate thermoregulatory responses during physical exertion or solar exposure, both of which are known to accelerate core temperature rise^54–56^. The exclusion of solar radiation was intentional, based on two considerations. First, at or near survival thresholds, individuals are unlikely to remain in direct sunlight due to innate behavioral responses such as seeking shade. Second, solar exposure is highly variable, ranging from ∼10 W/m² under cloud cover to >1,000 W/m² in direct sunlight^57^, complicating standardized exposure. Including this variability would obscure the physiological limits we sought to define.

In summary, we define the physiological limits of human heat survivability under prolonged humid heat exposure and present the first comprehensive empirical maps of core temperature dynamics across *T_w_* values from 32°C to 35°C. Our results confirm that *T_w_* is a scalable and physiologically robust index of heat stress: identical *T_w_* conditions elicited consistent core temperature responses across different temperature–humidity combinations. Females consistently demonstrated greater heat tolerance than males under all uncompensable conditions. Although *T_w_*=35°C is widely regarded as a universal survival limit, we show that it functions more accurately as a conservative threshold beyond which heat resilience declines with exposure time. These findings provide foundational evidence to recalibrate human heat tolerance thresholds and inform more equitable and data-driven climate adaptation strategies.

## METHODS

### Ethical approval

This study was approved by the Institutional Review Board of Xi’an University of Science and Technology (Approval No. XUST-IRB224005). All participants provided written informed consent prior to enrollment, in accordance with the Declaration of Helsinki.

### Participants and inclusion criteria

Forty-five healthy young adults (25 males and 20 females) were initially recruited. Nine individuals (5 males and 4 females) were excluded during pre-screening due to medical or physiological conditions deemed unsuitable for prolonged heat exposure, including cardiovascular disease, impaired glucose tolerance, or signs of systemic inflammation. The final study cohort comprised 36 participants (20 males and 16 females). To minimize hormonal variability, all female participants were tested during the early follicular phase of the menstrual cycle, when circulating progesterone levels are low^58^. Baseline characteristics are summarized in Supplementary Table S2. Young adults were selected based on recent epidemiological evidence indicating that individuals under 35 years of age account for 75% of heat-related deaths and 87% of heat-related years of life-lost, highlighting their disproportionate vulnerability to extreme heat^14^.

### Experimental conditions

To map human heat survivability under extreme humid conditions, we designed exposure protocols to assess whether core temperature (*T_core_*) responses remained consistent across a range of dry-bulb temperature (*T_db_*)and relative humidity (RH) combinations at identical wet-bulb temperatures (*T_w_*). Twelve distinct *T_db_*–RH pairings were selected to span four *T_w_* levels: 32°C, 33°C, 34°C, and 35°C. For each *T_w_* level, three thermodynamically distinct ambient conditions were used to ensure wide environmental coverage (Fig. 1b). At *T_w_*=32°C, participants were exposed to *T_db_*=36°C with RH=74.5%, *T_db_*=40°C with RH=55.0%, and *T_db_*=52°C with RH=22.5%. At *T_w_*=33°C, the corresponding conditions were *T_db_*=37°C with RH=74.8%, *T_db_*=42°C with RH=51.5%, and *T_db_*=47°C with RH=35.5%. For *T_w_*=34°C, we used *T_db_*=38°C with RH=75.1%, *T_db_*=43°C with RH=52.1%, and *T_db_*=48°C with RH=36.2%. Finally, *T_w_*=35°C trials included *T_db_*=40°C with RH=70.3%, *T_db_*=45°C with RH=49.0%, and *T_db_*=50°C with RH=34.4%. The temporal profile of *T_w_* during each trial was calculated using Roland Stull’s equation^59^, based on real-time dry-bulb temperature (*T_db_*) and RH measurements from three calibrated data loggers positioned at 0.1 m, 0.6 m, and 1.1 m above the floor, in accordance with ASHRAE Standard 55^60^. Fig. 1c presents the mean Tw values recorded across all trials for each of the four target *T_w_* levels, confirming the consistency of environmental control. Air velocity was maintained at 0.15±0.05 m/s, and indoor carbon dioxide (CO_2_) levels were kept below 900 ppm to ensure optimal air quality^61^.

*T_w_*=35°C exposures were included to directly test the theoretical upper survivability threshold proposed by Sherwood and Huber^16^, which suggests that survival is unlikely beyond six hours at this extreme *T_w_*. Incorporating multiple *T_db_* values at each *T_w_* erved two key objectives: first, to evaluate whether physiological strain remains consistent across different ambient conditions when *T_w_* is held constant; and second, to test the assumption that heat tolerance (or critical environmental limits) declines with rising ambient temperature, even at a fixed *T_w_*. To ensure internal validity, radiant heat exchange was minimized by eliminating direct radiant heat sources within the chamber, thereby ensuring that heat stress was induced primarily through convective and evaporative mechanisms.

### Experimental protocol

Participants completed twelve randomized, counterbalanced heat exposure trials, each separated by a one-week washout period to ensure full physiological recovery^62^. Trials were conducted between March 2023 and May 2025, excluding the months of June through September to avoid seasonal heat acclimatization. During these excluded months, outdoor temperatures frequently exceeded 30–32°C, whereas the remaining trial periods, ambient conditions remained cooler (<27°C, 40–70% RH)^63^, minimizing the likelihood of prior heat adaptation. Each trial commenced at 9:00 a.m. Upon arrival, participants provided a urine sample to assess hydration status. A urine specific gravity (USG) threshold of <1.020 was required to begin the trial^64^; individuals exceeding this threshold consumed an isotonic electrolyte solution until euhydration was achieved. Nude body weight was recorded before the trial and at 30-minute intervals throughout to monitor fluid balance.

Participants remained seated in a climate-controlled chamber (Espec Corporation, Osaka, Japan) and engaged in light cognitive activities such as reading, computer use, or using personal devices (e.g., iPads or smartphones). Clothing insulation was standardized to reflect typical summer office attire, with thermal resistance values of 0.40 clo for males and 0.41 clo for females^60^. To maintain euhydration during light indoor office work, participants consumed a flavored isotonic electrolyte beverage (zinc: 10.8 mg/L, carbohydrates: 0.5 g/L, sodium: 283 mg/L) to match sweat losses, determined by body weight measurements every 30 minutes (target: 0.1– 0.5 L/h based on anticipated sweat rates for light activity in the extreme heat). The sodium concentration, lower than average sweat sodium losses (∼35 mmol/L), was supplemented by normal dietary intake to maintain electrolyte balance. The beverage was maintained at 37°C to match body temperature. Hydration status was confirmed by ensuring fluid intake equaled sweat losses, preventing hypohydration (i.e., body water deficit >1% body weight loss)^65^. At 12:00 p.m., a standardized 550-kCal sandwich meal was provided to ensure consistent caloric intake across trials. Restroom access was permitted as needed; facilities were located within the testing area and maintained under thermoneutral conditions (23-27°C, 40-60% RH) to avoid thermal disruption.

### Physiological and perceptual measurements

Core temperature (*T_core_*) was continuously monitored using a rectal thermistor (YSI401, Yellow Spring Instrument, Yellow Springs, OH, USA; accurate: ±0.1°C) inserted 10 cm beyond the anal sphincter. Mean skin temperature (*T_sk_*) was calculated using four calibrated iButton sensors (DS1992L, Maxim Integrated, Sunnyvale, CA, USA; calibrated to accuracy: ±0.2°C) affixed at standard anatomical sites, with values averaged using the Ramanathan formula^66^. Heart rate was monitored continuously via chest strap (Polar Vantage XL, Polar Electro Oy, Kempele, Finland), and blood pressure was assessed every 30 minutes using an automated monitor (YE660CR, Yuwell, Yancheng, China). Mean arterial pressure (MAP) was subsequently derived to evaluate cardiovascular strain.

Subjective perceptions were recorded every 30 minutes and included thermal discomfort (7-point scale, from −3: very uncomfortable to +3: very comfortable^60^, thirst sensation (9-point scale, from 1: not thirsty at all to 9: very, very thirsty)^67^, and psychological stress (10-point anxiety scale, from 0: no stress/anxiety to 10: very severe stress/anxiety)^68^. Hydration status was verified via urine specific gravity (USG)^64^, measured before and after each trial using a digital refractometer (PAL-10S, ATAGO Co. Ltd., Tokyo, Japan).

Metabolic rates were assessed for 10 minutes before and after exposure using a portable indirect calorimetry system (COSMED K5, COSMED Srl, Rome, Italy), with the final 5 minutes of steady-state data used for analysis. To minimize participant burden, metabolic testing was conducted only under three representative dry-bulb temperature–relative humidity (*T_db_*-RH) conditions corresponding to *T_w_*=33°C: *T_db_*=37°C with RH=74.8%, *T_db_*=42°C with RH=51.5%, and *T_db_*=47°C with RH=35.5%. Full metabolic profiling was not performed under all twelve environmental conditions due to discomfort and breathing difficulty reported while wearing the measurement mask at higher heat loads. Each participant completed the designated three tests, yielding 60 valid trials among males and 48 among females (Supplementary Video S2). Thermophysiological and perceptual response data are summarized in Supplementary Table S3. Metabolic equivalent (MET) values were calculated using the conventional definition of 1 MET = 1 kcal·kg⁻¹·h⁻¹ (∼4.184 kJ/kg/h), rather than the standard oxygen - uptake–based definition of 1 MET = 3.5 ml O₂·kg⁻¹·min⁻¹. This choice was made because the oxygen- based conversion systematically overestimated metabolic rate compared to caloric values derived from the COSMED system—a discrepancy consistent with previous findings that the 3.5 ml·kg-1·min-1 standard overestimates resting energy expenditure by 20–35 % across diverse populations^69,70^.

### Trial termination criteria

All exposure trials were continuously monitored by a registered nurse certified in Shaanxi Province to ensure participant safety. Trials were terminated immediately if any of the following safety criteria were met: (1) core temperature reached 39.0°C; (2) sustained cardiovascular strain, defined as heart rate exceeding 90% of the age-predicted maximum (HRmax = 208 − 0.7 × age) for more than five consecutive minutes^71^, or a drop in systolic blood pressure below 90 mmHg^72^; (3) onset of severe subjective symptoms, including dizziness, limb numbness, nausea, or pronounced discomfort; or (4) clinical judgment by the supervising nurse indicated a risk to participant safety, regardless of quantitative criteria.

### Environmental compensability assessment and *T_core_* responses

To evaluate environmental compensability across the twelve conditions, we analyzed the rate of core temperature (*T_core_*) increase during each exposure. Conditions were classified as (quasi-) compensable if the *T_core_* rise rate remained below 0.10°C/h^23,25,73,74^, indicating sufficient thermoregulatory capacity to prevent sustained heat accumulation. For these conditions, individual *T_core_* trajectories were smoothed using the *smooth.spline* function in R (v4.5.0), and rates of change was calculated in 5-minute intervals. Thermal steady state was defined by either a *T_core_* rise rate ≤ 0.10°C/h after three hours of exposure or a fluctuation of less than 0.10°C during the final 30 minutes, with the steady state time point assigned as 30 minutes before trial completion.

Although *T_core_* rise rates were minimal (< 0.1°C/h) under (quasi-)compensable conditions, we projected the theoretical time to reach the clinical heatstroke threshold of 40.5°C^26,27^, as a worst-case estimate. This was calculated by extrapolating beyond eight-hour exposure, assuming a maximum *T_core_* increase rate of 0.10°C/h, and adding the time required to reach 40.5°C from each participant’s final recorded *T_core_*.

For uncompensable conditions, in which thermoregulation failed to stabilize *T_core_*, we estimated the time to reach 40.5°C^26^ by fitting individual-level linear regressions to observed *T_core_* data. Using a baseline *T_core_* of 36.8°C, we standardized the rate of rise as (40.5°C − 36.8°C) over six hours, yielding a reference slope of 0.617°C/h. This consistent extrapolation framework allowed direct comparison of thermal strain between (quasi-)compensable and uncompensable conditions.

### Statistics

To determine whether different dry-bulb temperature and relative humidity combinations producing identical wet-bulb temperatures (*T_w_*) elicited comparable core temperature (*T_core_*) responses, we performed two-way repeated-measures ANOVAs across the three environmental conditions at each *T_w_* level. Additional analyses examined the effects of dry-bulb temperature and sex on estimated survival time at each *T_w_*. Significant main effects were followed by paired-samples t-tests with Bonferroni correction. All tests were two-tailed with significance set at *p*<0.05. Assumptions of normality and sphericity were confirmed, and Greenhouse-Geisser corrections applied as necessary.

## RESOURCE AVAILABILITY

### Lead contact

Requests for further information and resources should be directed to and will be fulfilled by the lead contact, Faming Wang.

### Materials availability

This study did not generate new unique materials.

### Data and code availability

All data supporting this study are provided in the Supplementary Information and have been deposited in figshare (doi:10.6084/m9.figshare.29476616.v1).

## ACKNOWLEDGEMENTS

This research was supported by the National Excellent Young Scientist Program (Grant No. 6119924022, to F.W.) and the “Sanqin Scholars” Plan of Shaanxi Province (Grant No. 2050225003, to F.W.). We sincerely thank all participants of the survival study for their time and invaluable contributions.

## AUTHOR CONTRIBUTIONS

F.W. conceived the project, designed the experiments, acquired funding, and wrote the manuscript. F.W., Y.X. and X.Z. prepared the illustrations and figures. H.W., Y.X. and F.W. performed the experiments and analyzed the data. Y.Z. provided the metabolic rate testing equipment. L.H., X.S., X.Z., Z.L., B.H., P.M., and Z.Z. supervised the research and critically revised the manuscript.

## DECLARATION OF INTERESTS

The authors declare no competing interests.

## SUPPLEMENTAL INFORMATION

Supplemental information has been provided and uploaded to the peer review system.

## Notes

### Competing Interest Statement

The authors have declared no competing interest.

