## Supplementary material for "Mapping Human Survivability at Extreme Wet-Bulb Temperatures 32–35°C"

**This file includes:**

Supplementary Figures S1 to S39

Supplementary Tables S1 to S3

Supplementary Videos S1 to S2

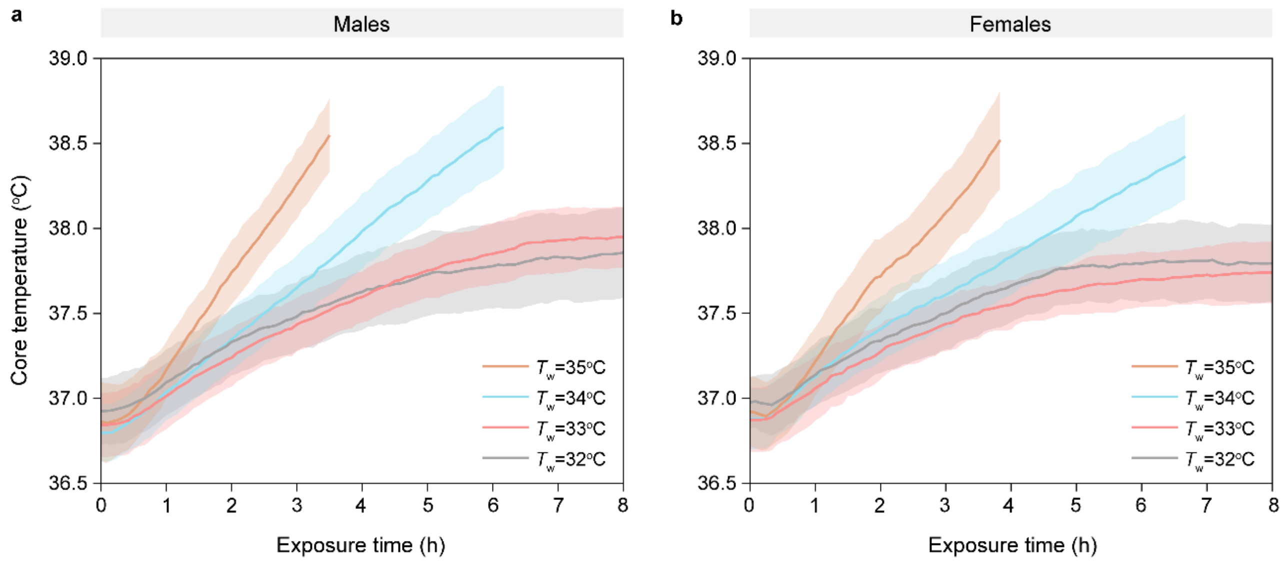


**Supplementary Fig. S1. Time-course mean core temperature responses of males and females at $T_w$=32-35°C. a**, males (n=20); **b**, females (n=16). Each line represents the mean core temperatures averaged across three dry-bulb temperature ($T_{db}$) and relative humidity (RH) conditions at each $T_w$ level, measured over 3.6-8.0 hours during controlled heat exposure trials.

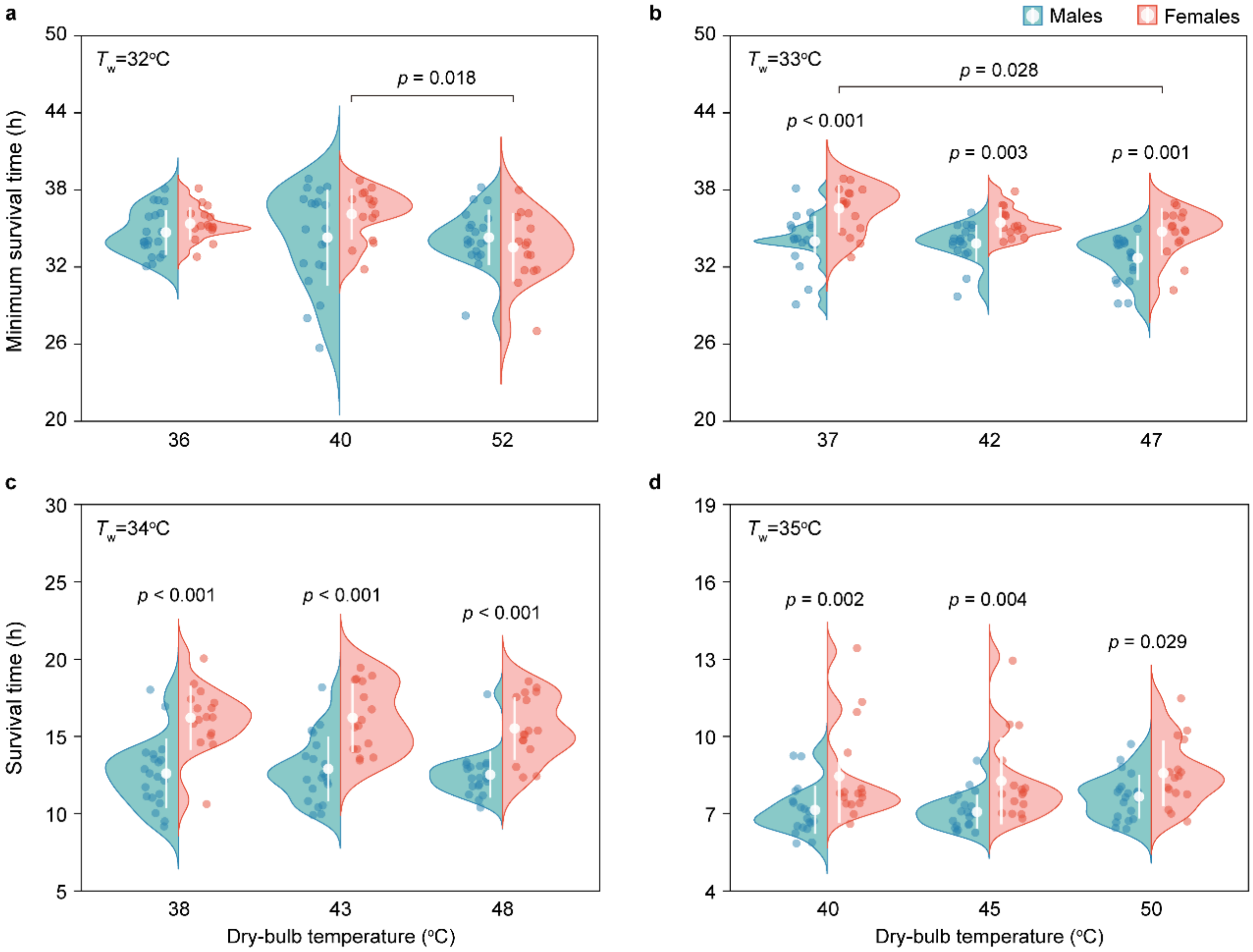


**Supplementary Fig. S2. Projected times to reach the clinical heatstroke threshold ($T_{core}$=40.5°C) at wet-bulb temperatures ($T_w$) of 32-35°C.** **a**, $T_w$=32°C; **b**, $T_w$=33°C c, $T_w$=34°C; **d**, $T_w$=35°C. Each split violin plot represents the projected time (in hours) for mean core temperature ($T_{core}$) to reach 40.5°C, calculated from heat exposure trials under three $T_{db}$-RH combinations at the specified $T_w$ levels. Data are based on 20 male and 16 female participants, with $T_{core}$ measured using rectal probes at 5-minute intervals over 3.6-8.0 hours. Projections were derived using linear extrapolation of $T_{core}$ rise rates from Supplementary Fig. S1.

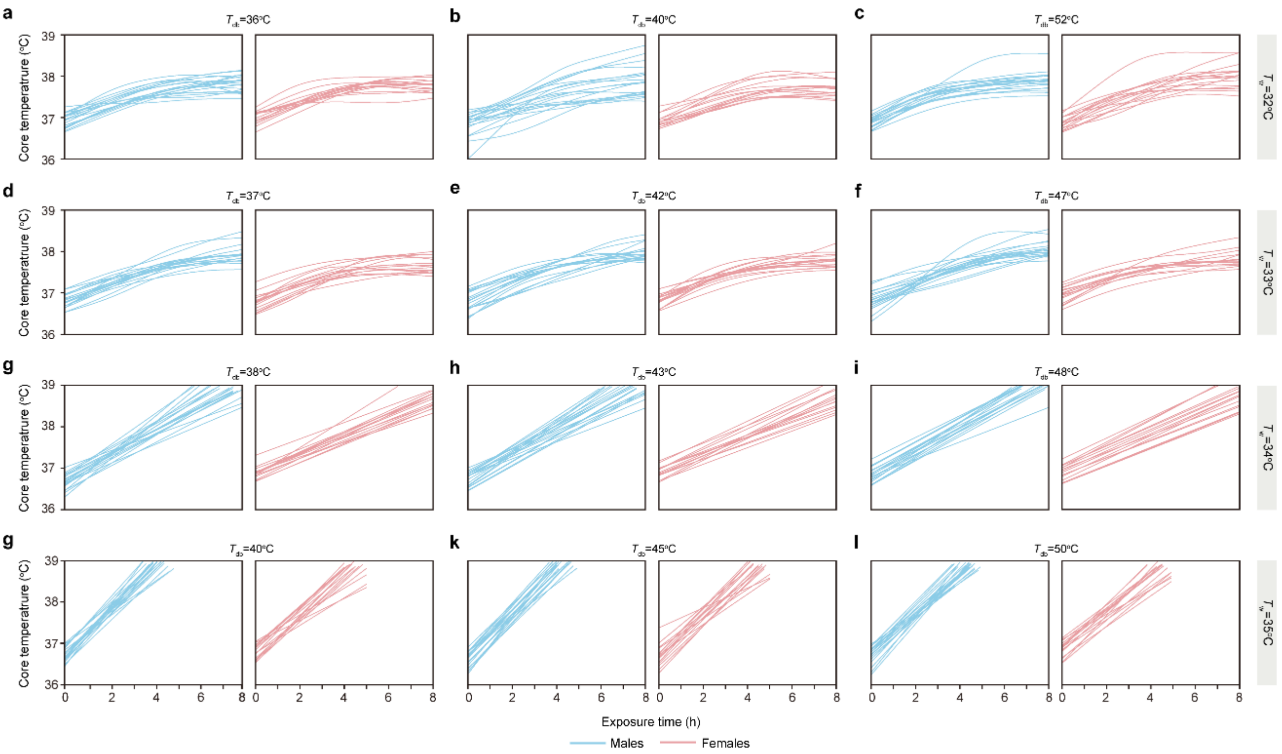


**Supplementary Fig. S3. Individual core temperature responses across wet-bulb temperatures ($T_w$) of 32-35°C at 12 varying dry-bulb temperatures ($T_{db}$). a**, $T_{db}$=36°C, **b**, $T_{db}$=40°C, **c**, $T_{db}$=52°C, **d**, $T_{db}$=37°C, **e**, $T_{db}$=42°C, **f**, $T_{db}$=47°C, **g**, $T_{db}$=38°C, **h**, $T_{db}$=43°C, **i**, $T_{db}$=48°C, **j**, $T_{db}$=40°C, k, $T_{db}$=45°C, l, $T_{db}$=50°C. Each line represents an individual participant's core temperature ($T_{core}$,°C) over time at $T_w$ levels of 32–35°C, measured during controlled heat exposure trials with 36 participants (20 males, 16 females). $T_{core}$ was continuously recorded using rectal probes at 5-minute intervals over 3.6-8.0 hours.

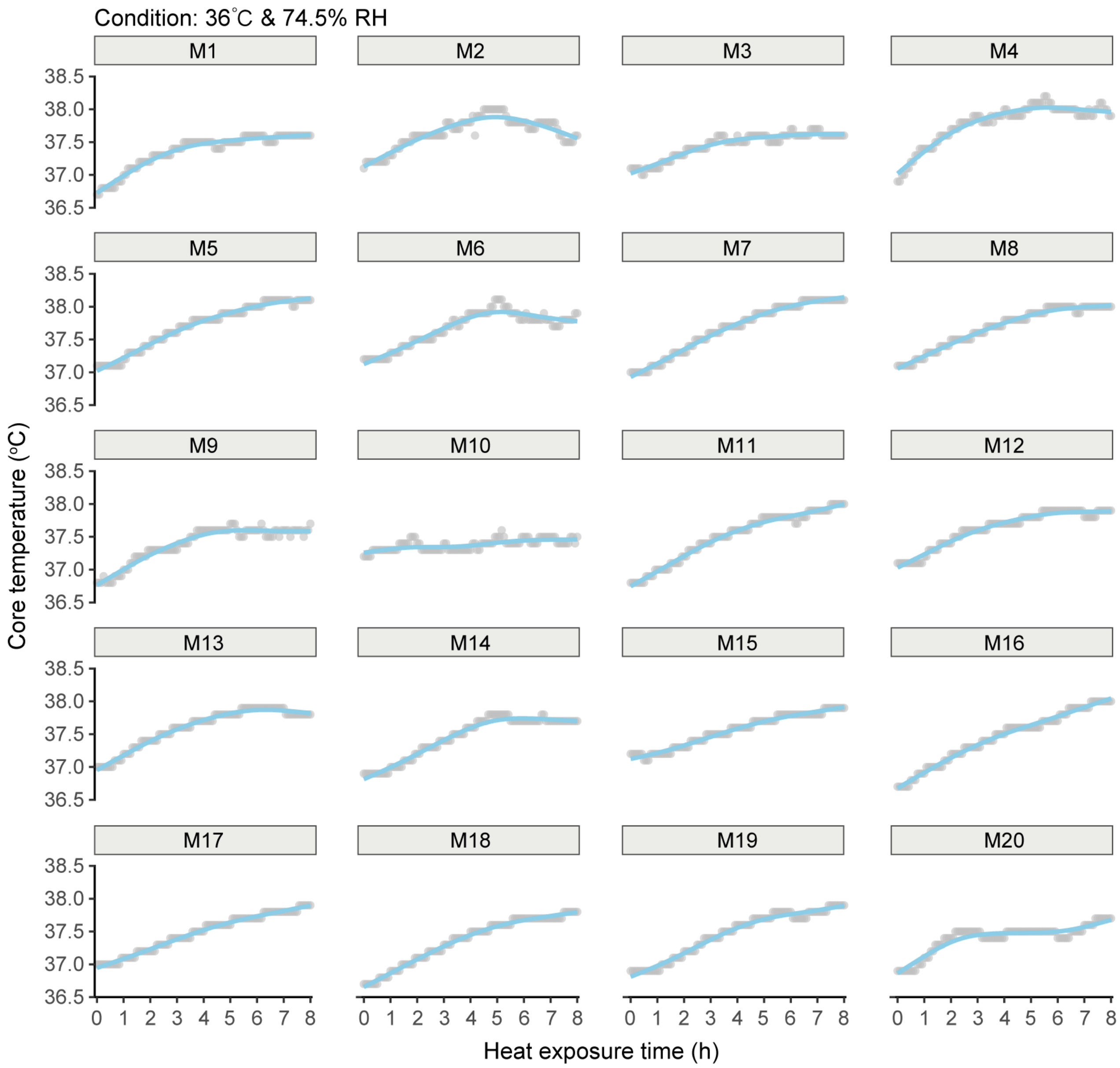


**Supplementary Fig. S4. Individual core temperature trajectories during 8-hour heat exposure in 20 young adult males at 36°C and 74.5% relative humidity.** Core temperature ($T_{core}$) was continuously monitored using a rectal thermistor. Each line represents a single participant's response under the specified environmental condition.

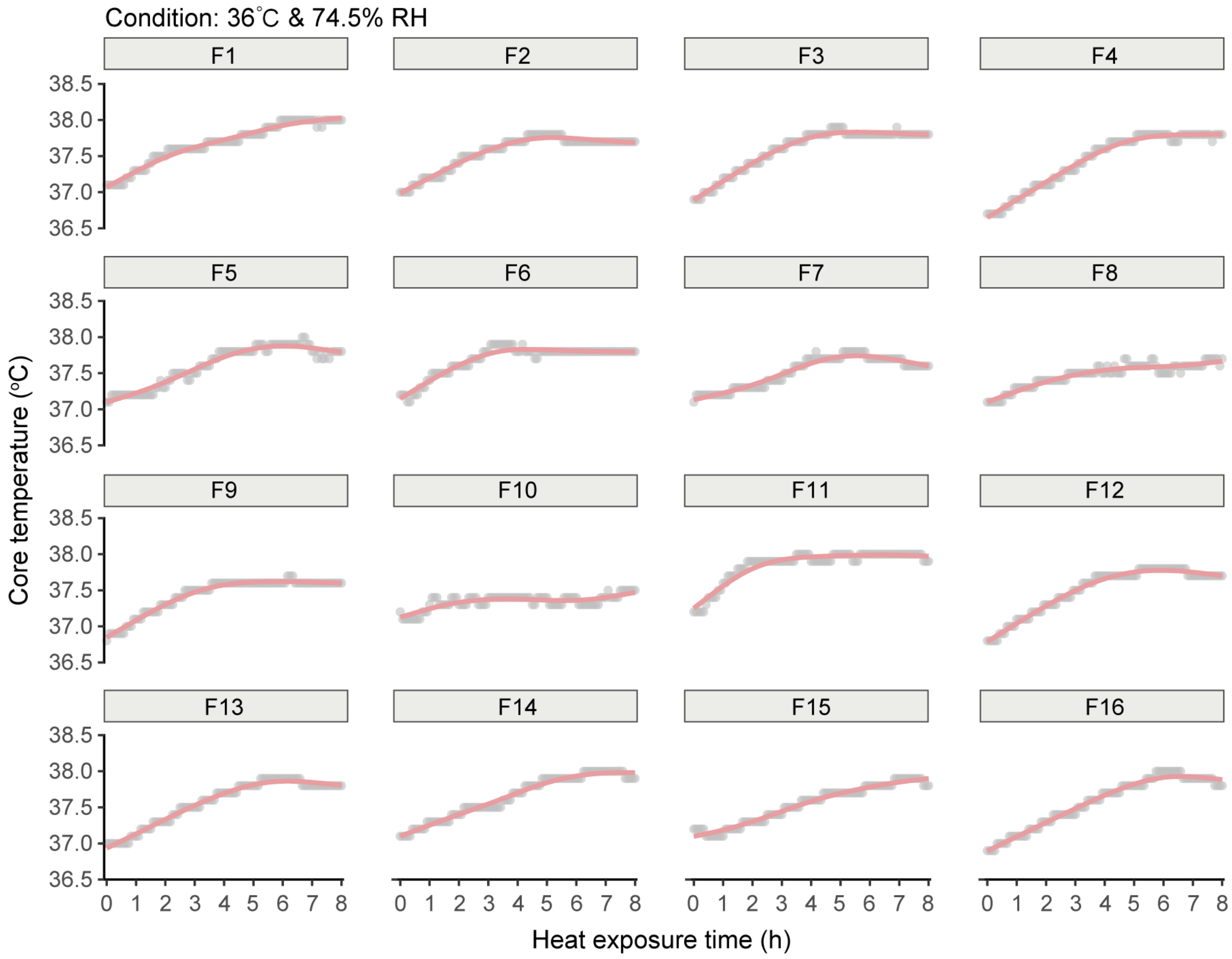


**Supplementary Fig. S5. Individual core temperature trajectories during 8-hour heat exposure in 16 young adult females at 36°C and 74.5% relative humidity.** Core temperature ($T_{core}$) was continuously monitored using a rectal thermistor. Each line represents a single participant's response under the specified environmental condition.

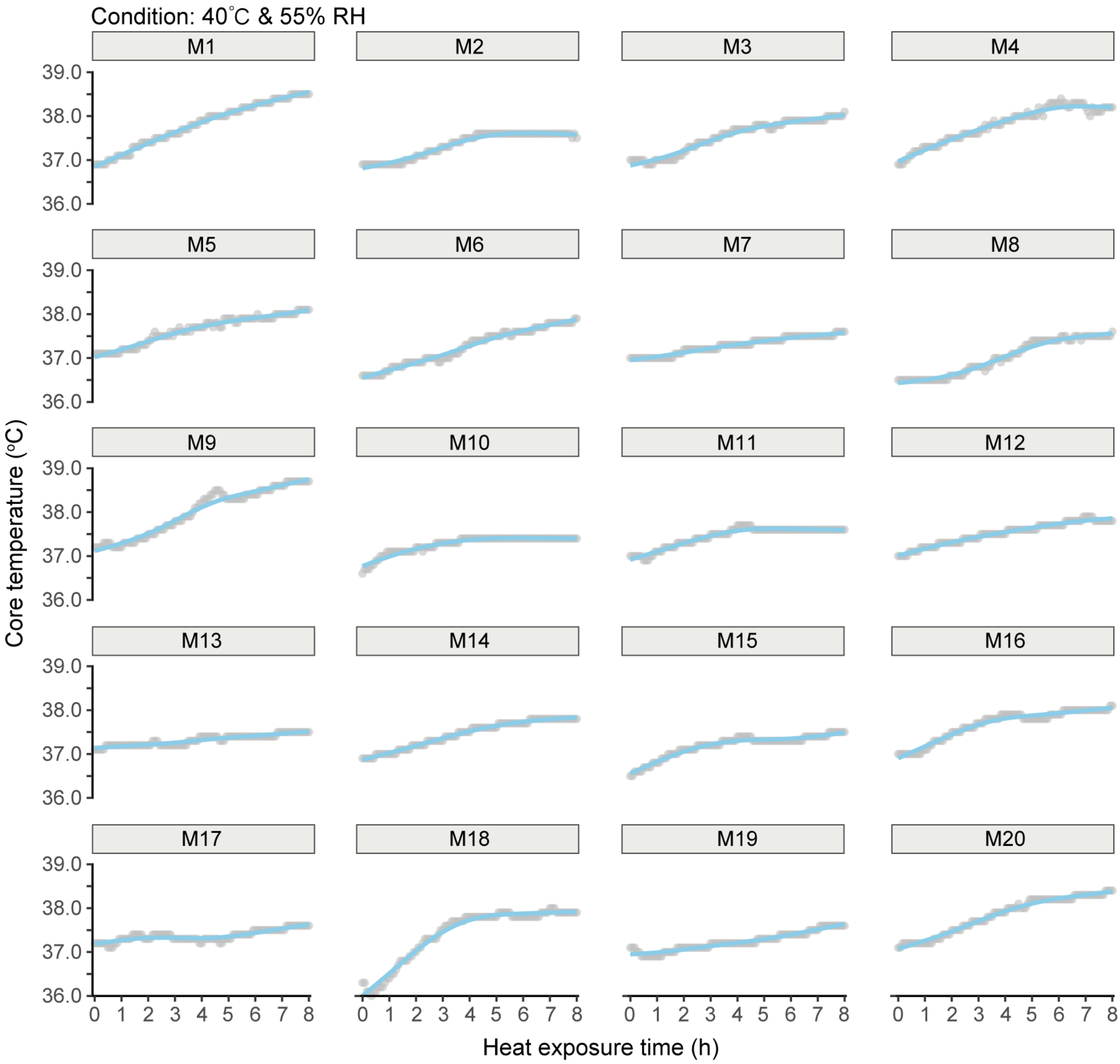


**Supplementary Fig. S6. Individual core temperature trajectories of 20 young adult males during 8-hour heat exposure at 40°C and 55% relative humidity.** Each line represents the core temperature ($T_{core}$) of a single participant, continuously measured using a rectal thermistor. Data illustrate inter-individual variability in thermoregulatory responses under uncompensable heat conditions.

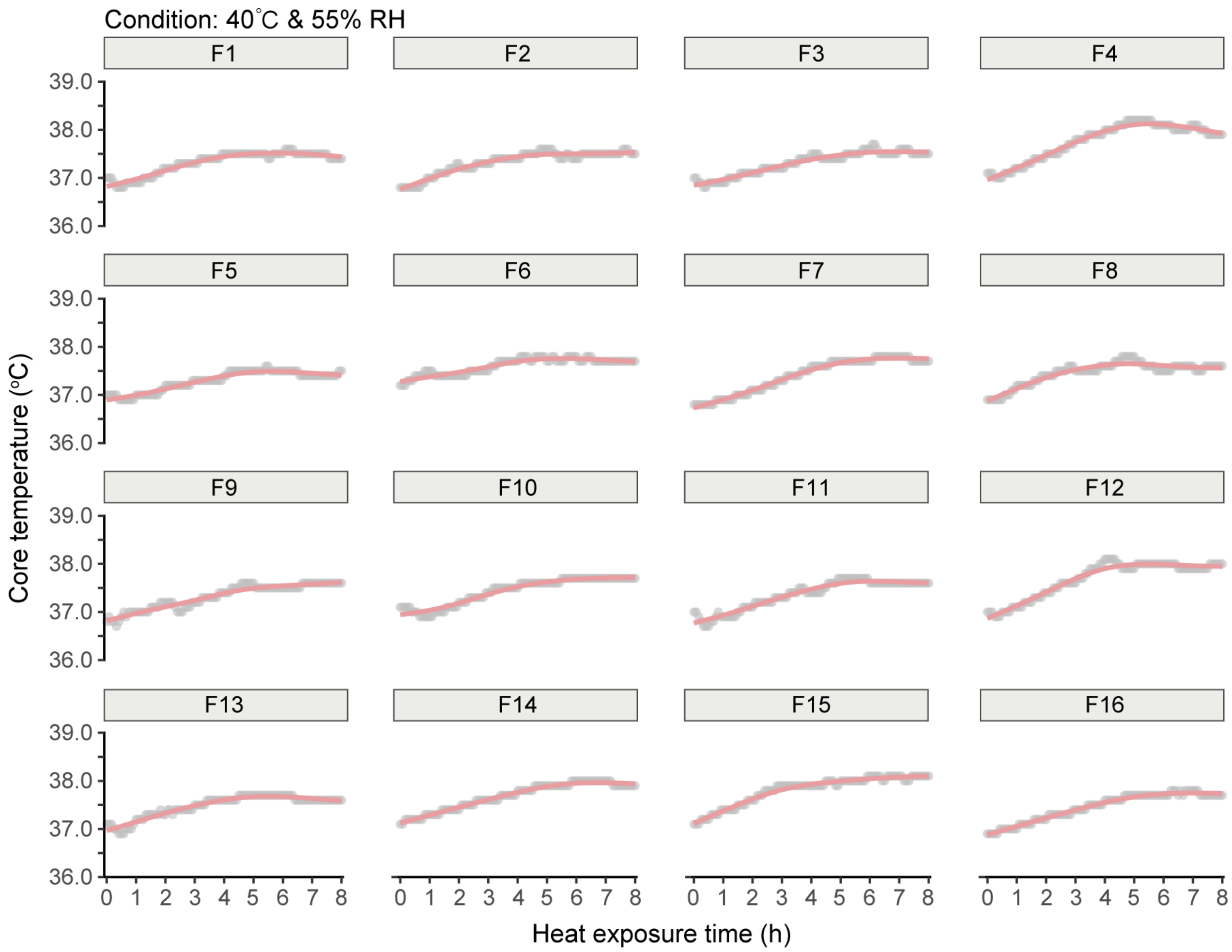


**Supplementary Fig. S7. Individual core temperature trajectories of 16 young adult females during 8-hour heat exposure at 40°C and 55% relative humidity.** Each line represents the core temperature ($T_{core}$) of a single participant, continuously measured using a rectal thermistor. Data illustrate inter-individual variability in thermoregulatory responses under uncompensable heat conditions.

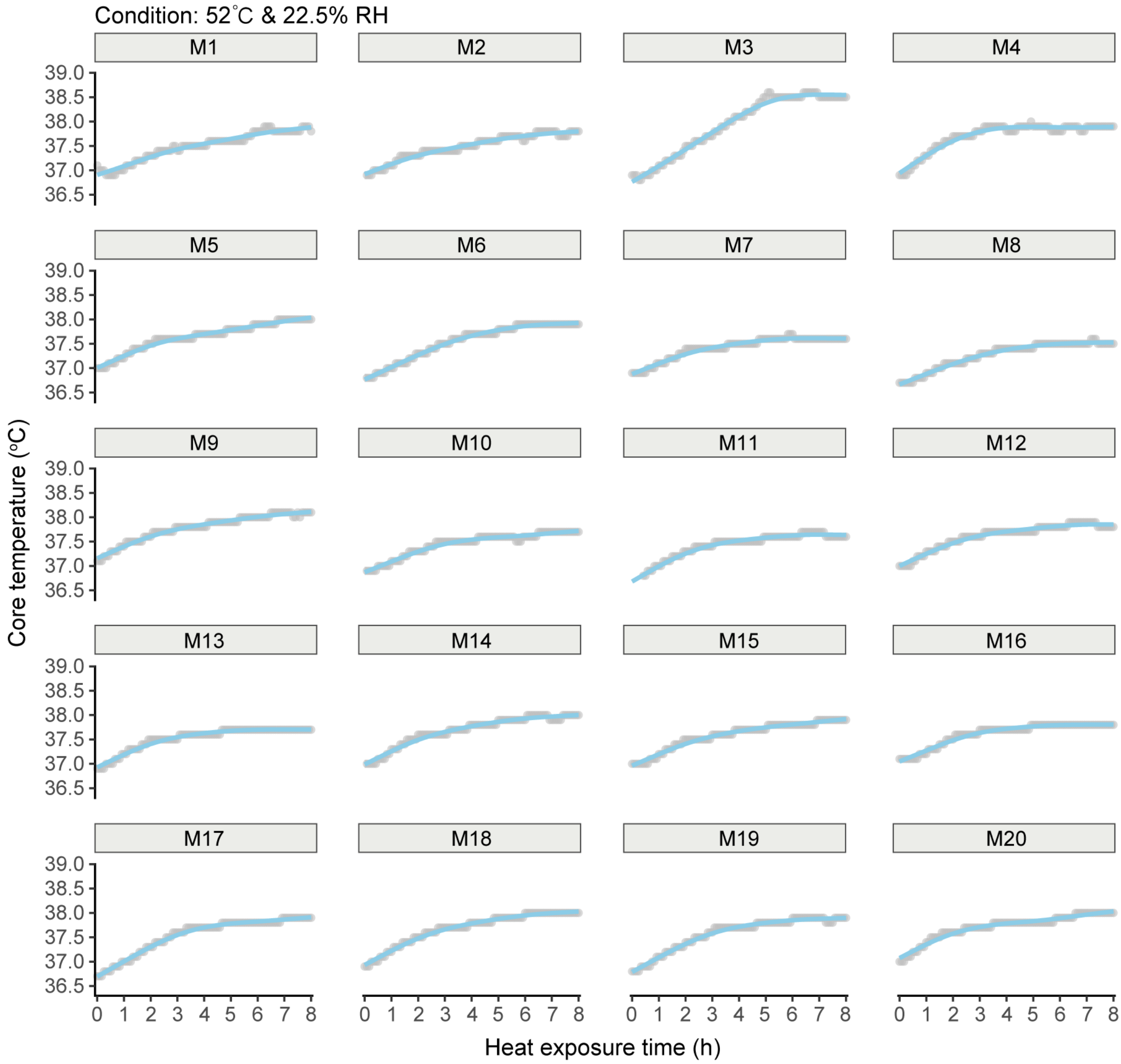


**Supplementary Fig. S8. Individual core temperature trajectories of 20 young adult males during 8-hour heat exposure at 52°C and 22.5% relative humidity.** Core temperature ($T_{core}$) was continuously monitored using a rectal thermistor. The data highlight inter-individual variability in thermoregulatory strain under extremely hot and humid conditions approaching the theoretical survivability threshold.

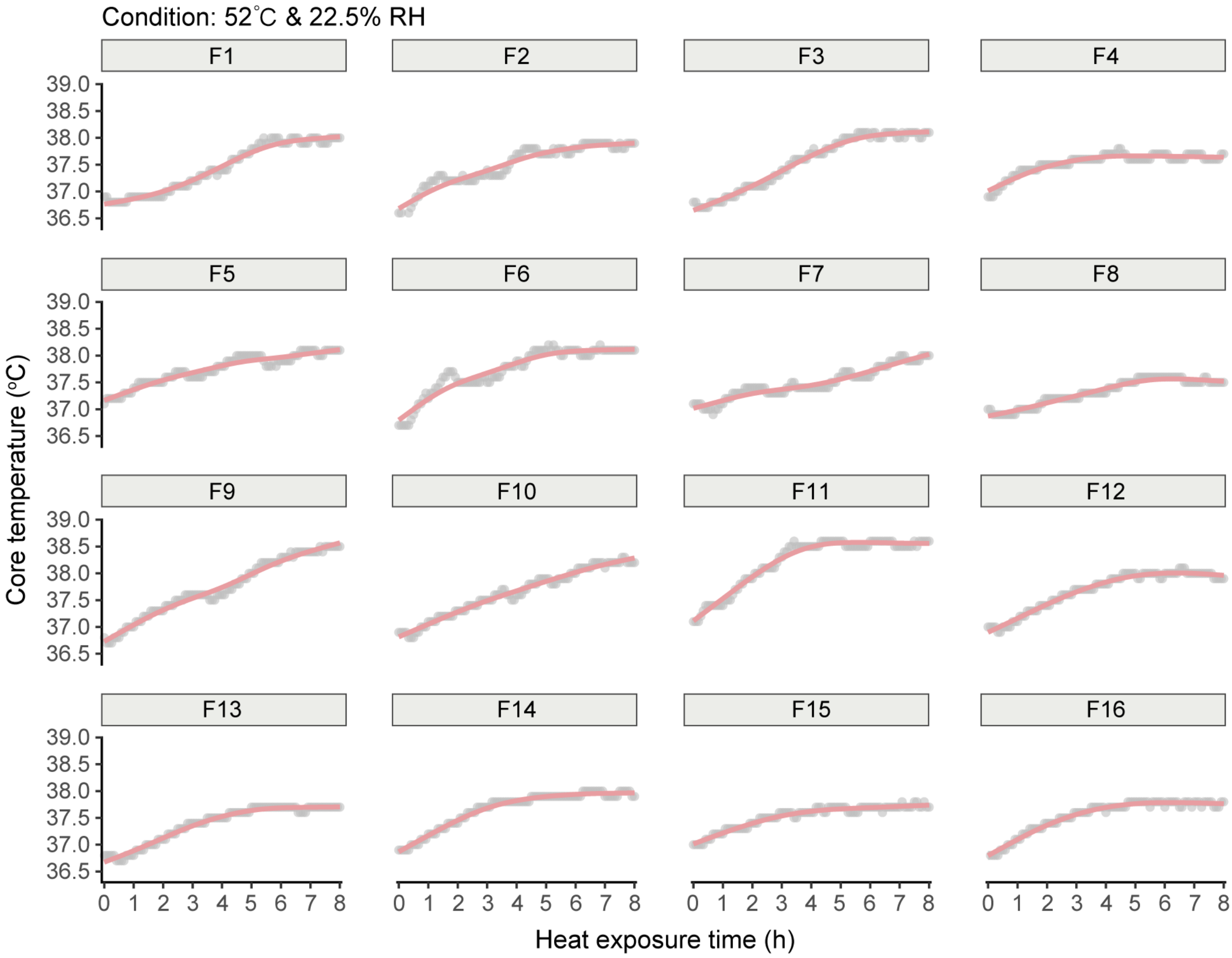


**Supplementary Fig. S9. Individual core temperature trajectories of 16 young adult females during 8-hour heat exposure at 52°C and 22.5% relative humidity.** Core temperature ($T_{core}$) was continuously monitored using a rectal thermistor. The data highlight inter-individual variability in thermoregulatory strain under extremely hot and humid conditions approaching the theoretical survivability threshold.

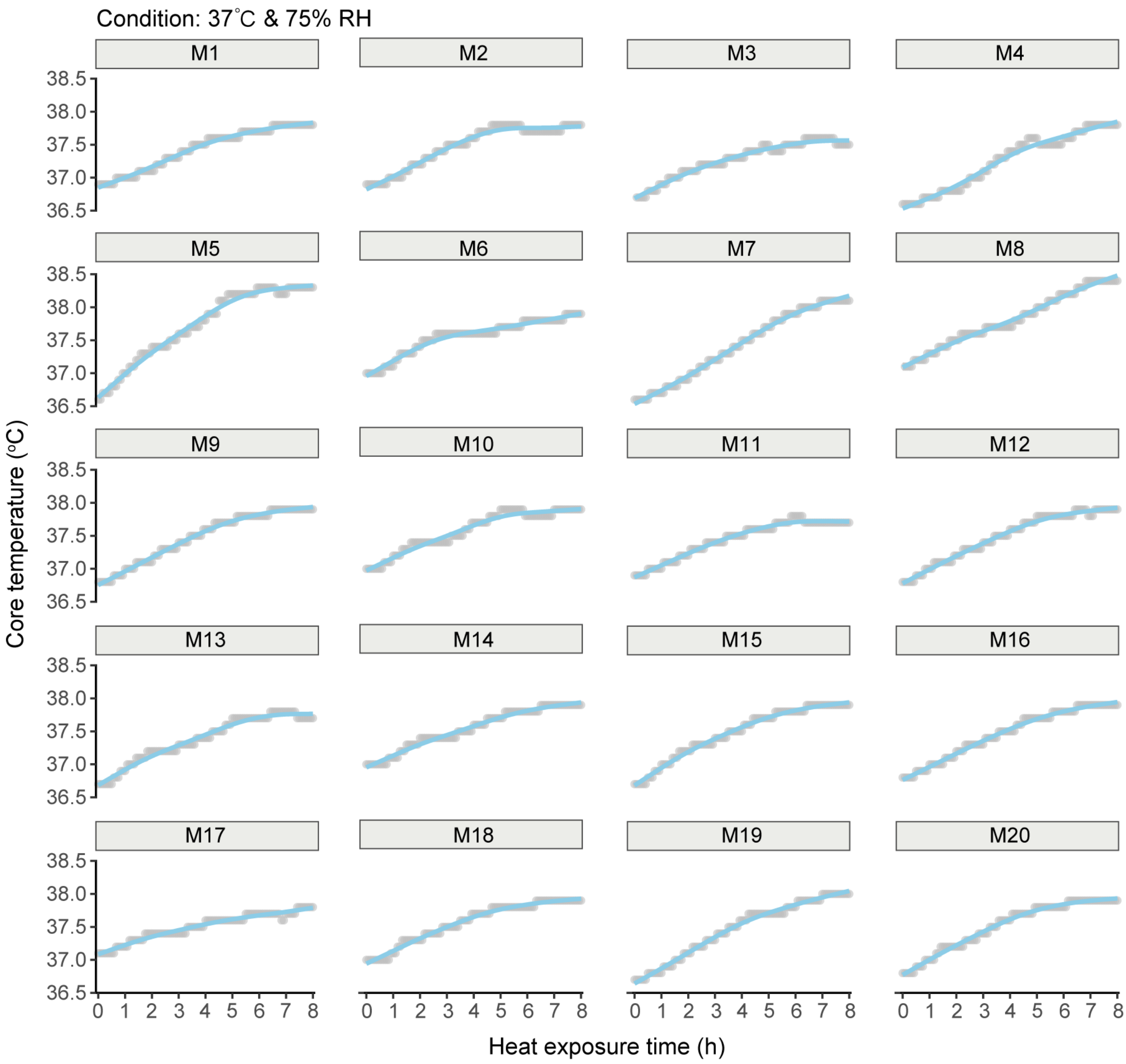


**Supplementary Fig. S10. Individual core temperature trajectories of 20 young adult males during 8-hour heat exposure at 37°C and 74.8% relative humidity.** Core temperature ($T_{core}$) was continuously monitored using a rectal thermistor. The data highlight inter-individual variability in thermoregulatory strain under extremely hot and humid conditions approaching the theoretical survivability threshold.

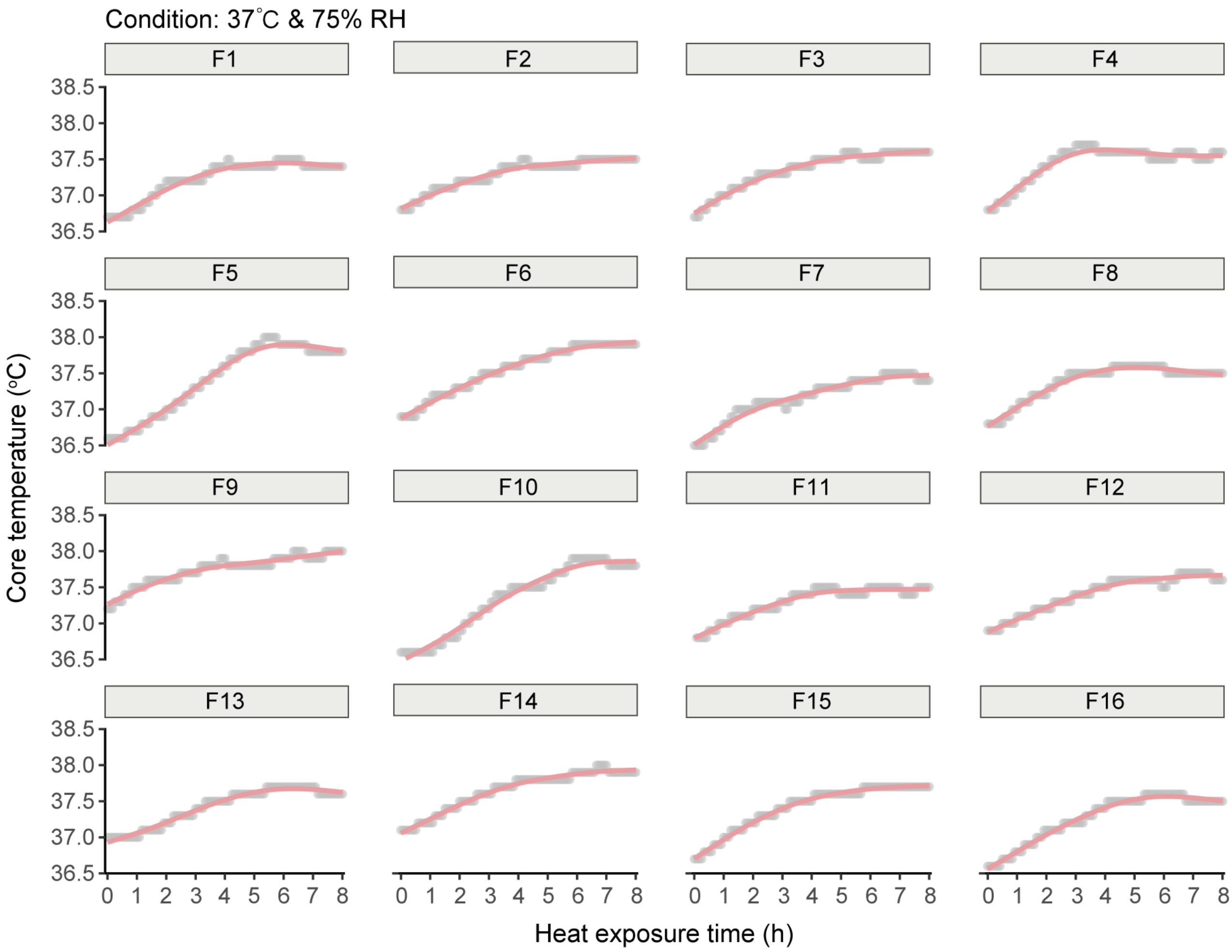


**Supplementary Fig. S11. Individual core temperature trajectories of 16 young adult females during 8-hour heat exposure at 37°C and 74.8% relative humidity.** Core temperature ($T_{core}$) was continuously monitored using a rectal thermistor. The data highlight inter-individual variability in thermoregulatory strain under extremely hot and humid conditions approaching the theoretical survivability threshold.

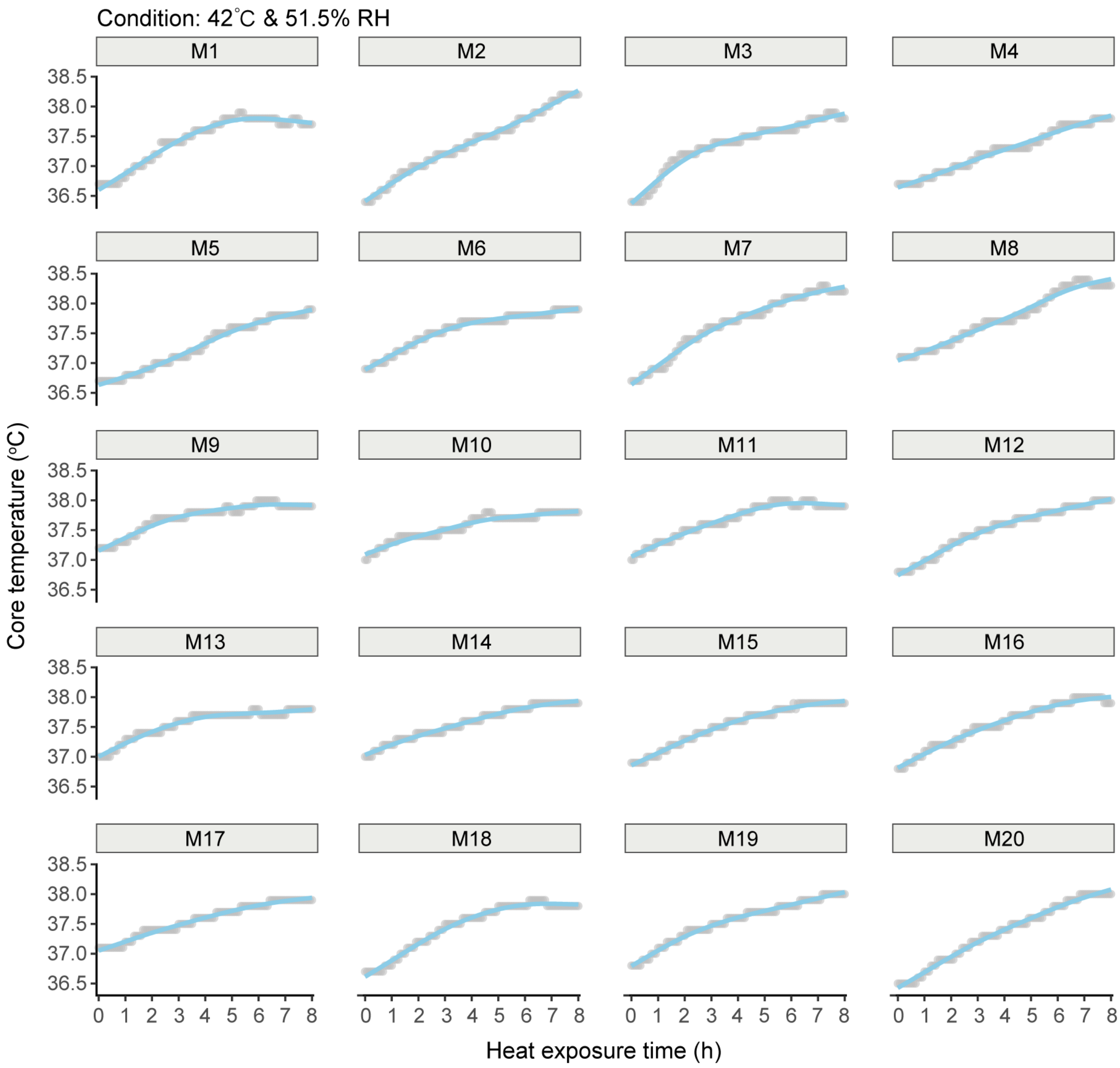


**Supplementary Fig. S12. Individual core temperature trajectories of 20 young adult males during 8-hour heat exposure at 42°C and 51.5% relative humidity.** Core temperature ($T_{core}$) was continuously monitored using a rectal thermistor. The data highlight inter-individual variability in thermoregulatory strain under extremely hot and humid conditions approaching the theoretical survivability threshold.

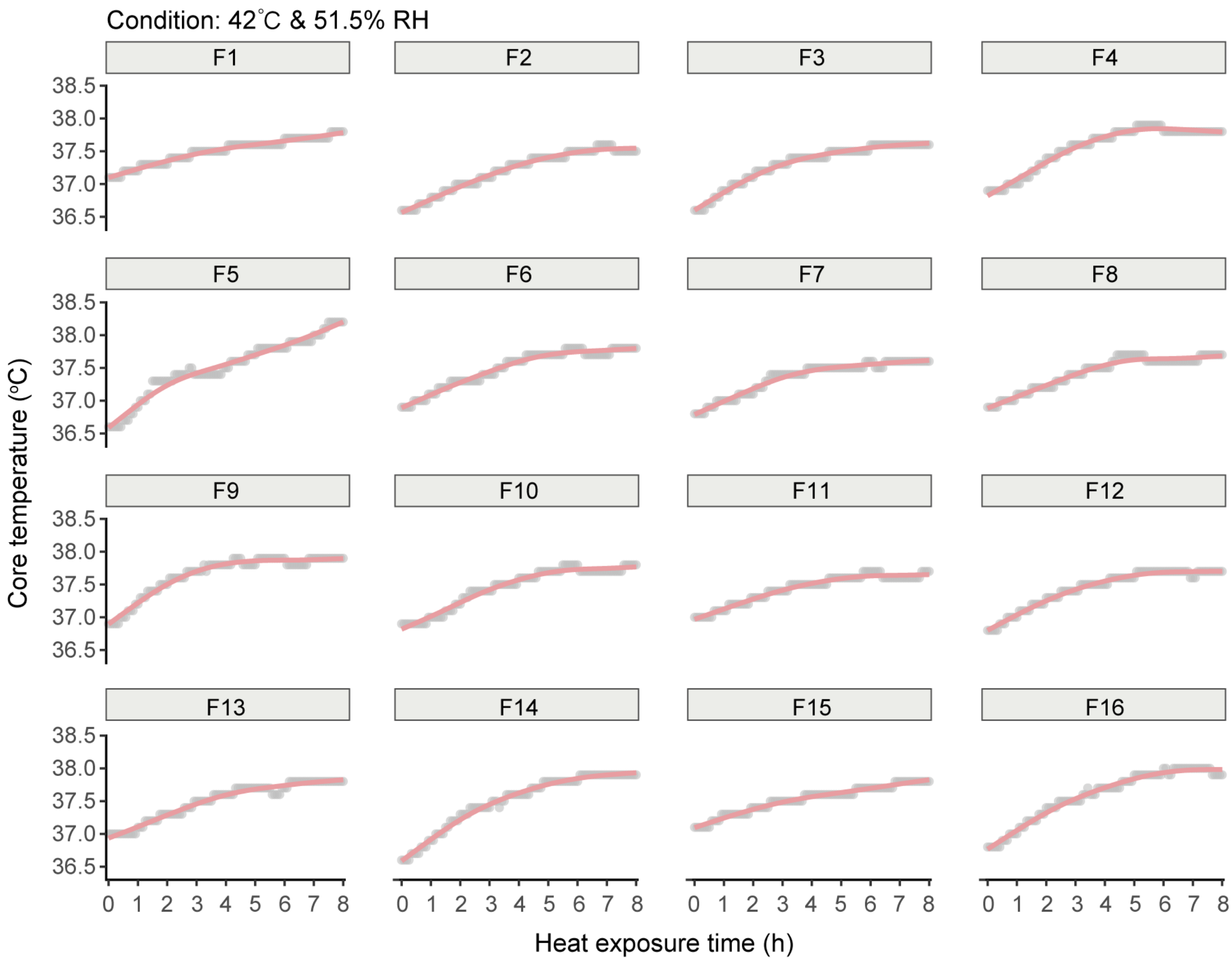


**Supplementary Fig. 1**3. Individual core temperature trajectories of 16 young adult females during 8-hour heat exposure at 42°C and 51.5% relative humidity. Core temperature (Tcore) was continuously monitored using a rectal thermistor. The data highlight inter-individual variability in thermoregulatory strain under extremely hot and humid conditions approaching the theoretical survivability threshold.

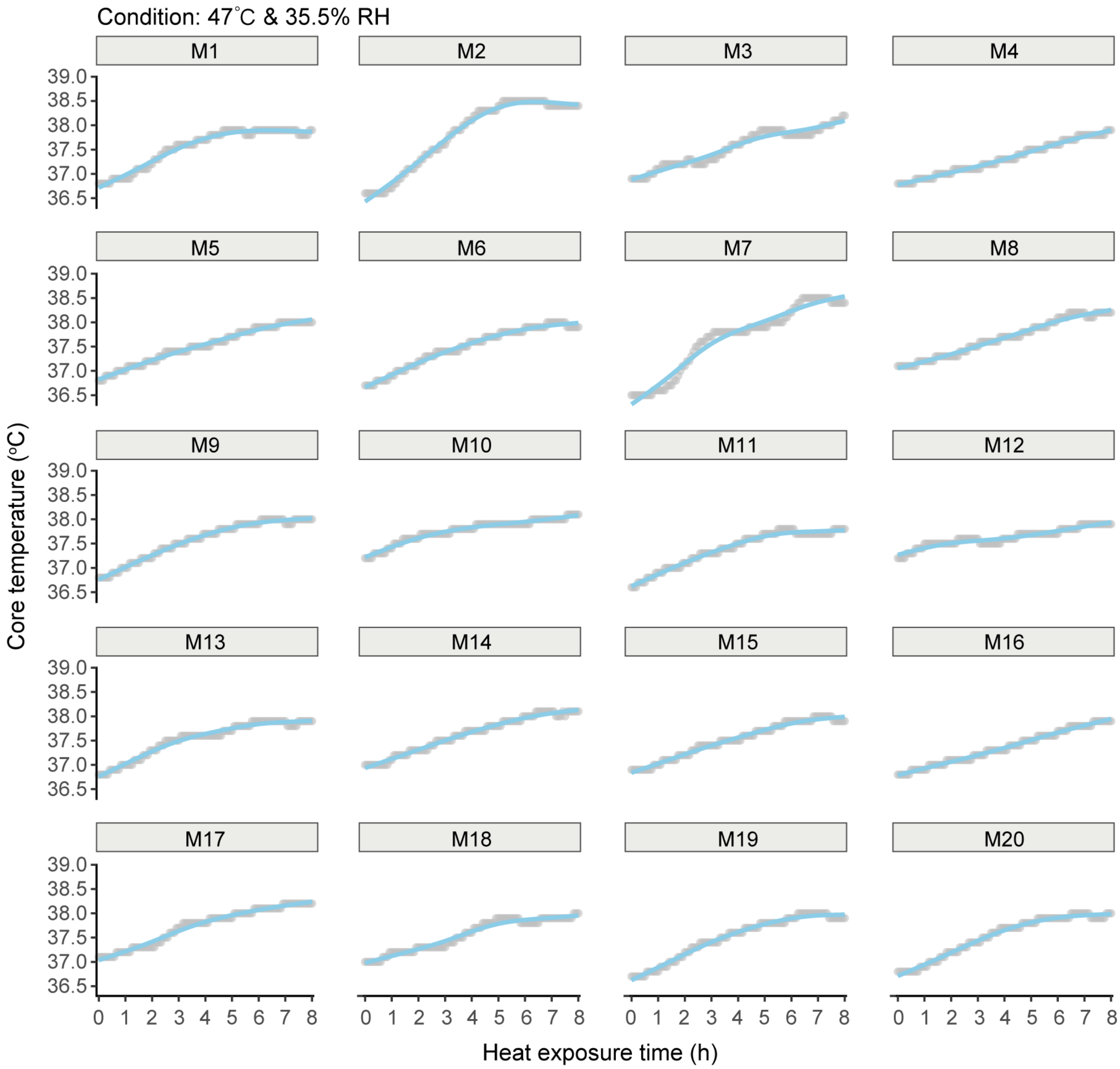


**Supplementary Fig. S14. Individual core temperature trajectories of 20 young adult males during 8-hour heat exposure at 47°C and 35.5% relative humidity.** Core temperature ($T_{core}$) was continuously monitored using a rectal thermistor. The data highlight inter-individual variability in thermoregulatory strain under extremely hot and humid conditions approaching the theoretical survivability threshold.

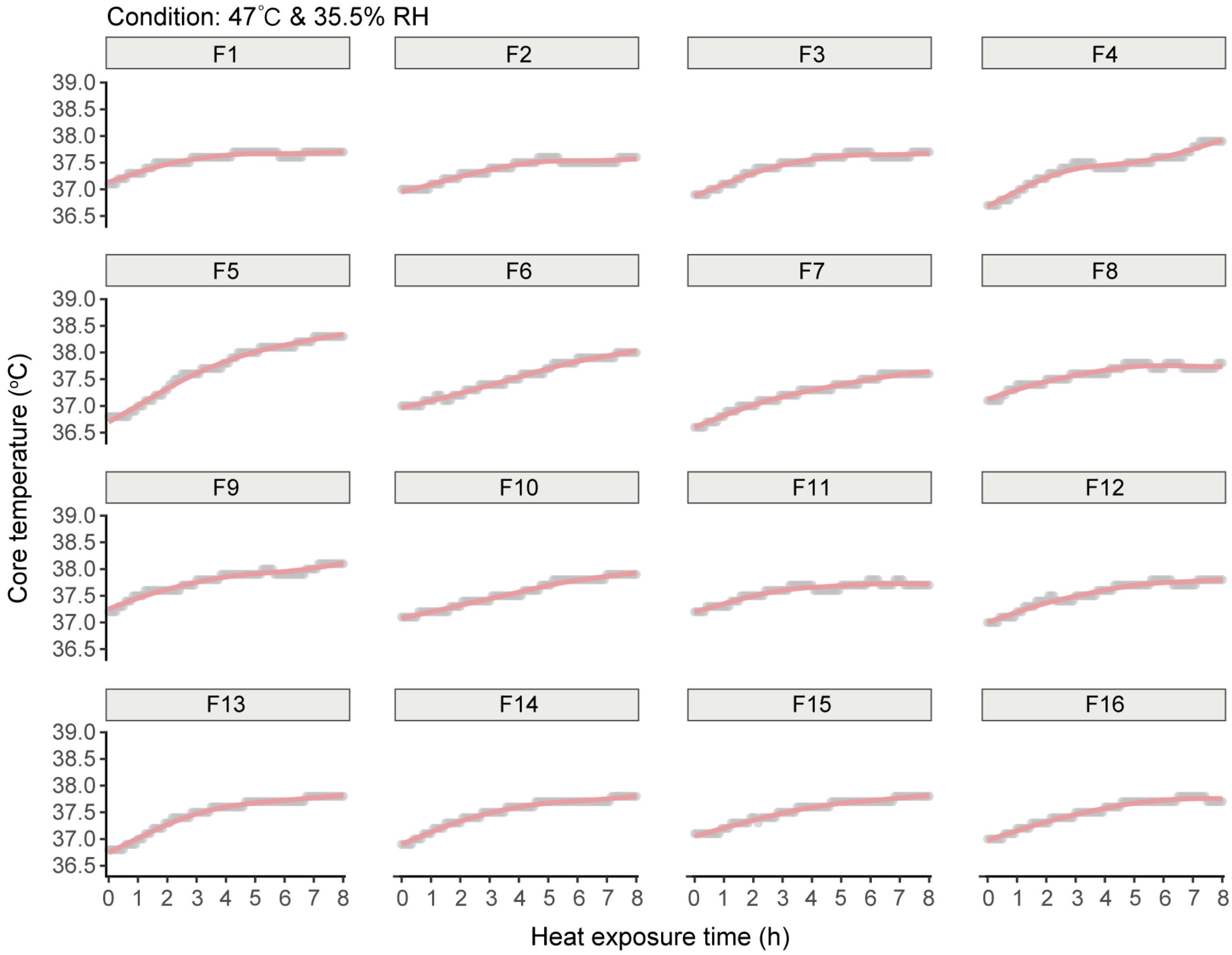


**Supplementary Fig. S15. Individual core temperature trajectories of 16 young adult females during 8-hour heat exposure at 47°C and 35.5% relative humidity.** Core temperature ($T_{core}$) was continuously monitored using a rectal thermistor. The data highlight inter-individual variability in thermoregulatory strain under extremely hot and humid conditions approaching the theoretical survivability threshold.

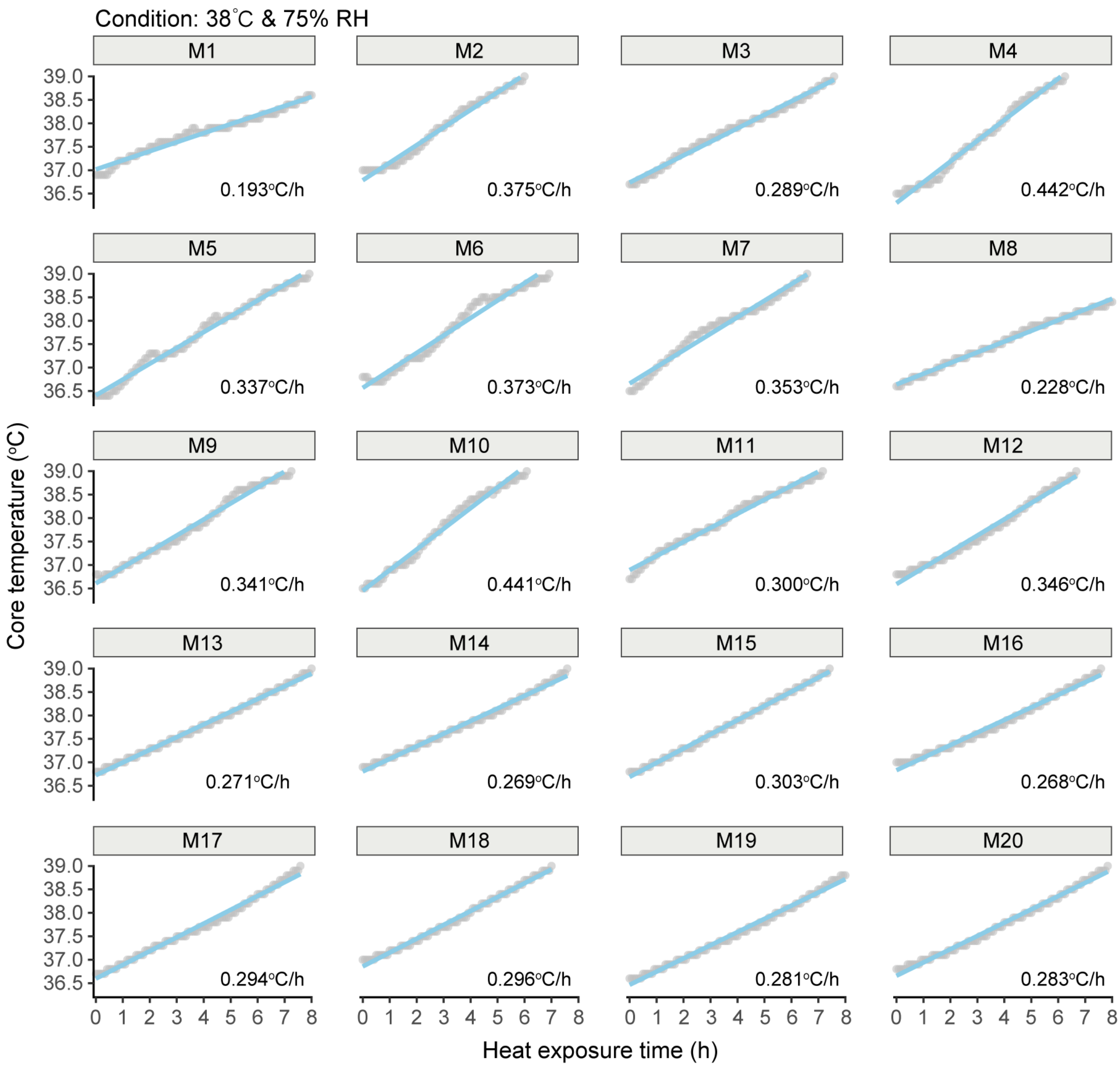


**Supplementary Fig. S16. Individual core temperature trajectories of 20 young adult males during 8-hour heat exposure at 38°C and 75.1% relative humidity.** Core temperature ($T_{core}$) was continuously monitored using a rectal thermistor. The data highlight inter-individual variability in thermoregulatory strain under extremely hot and humid conditions approaching the theoretical survivability threshold.

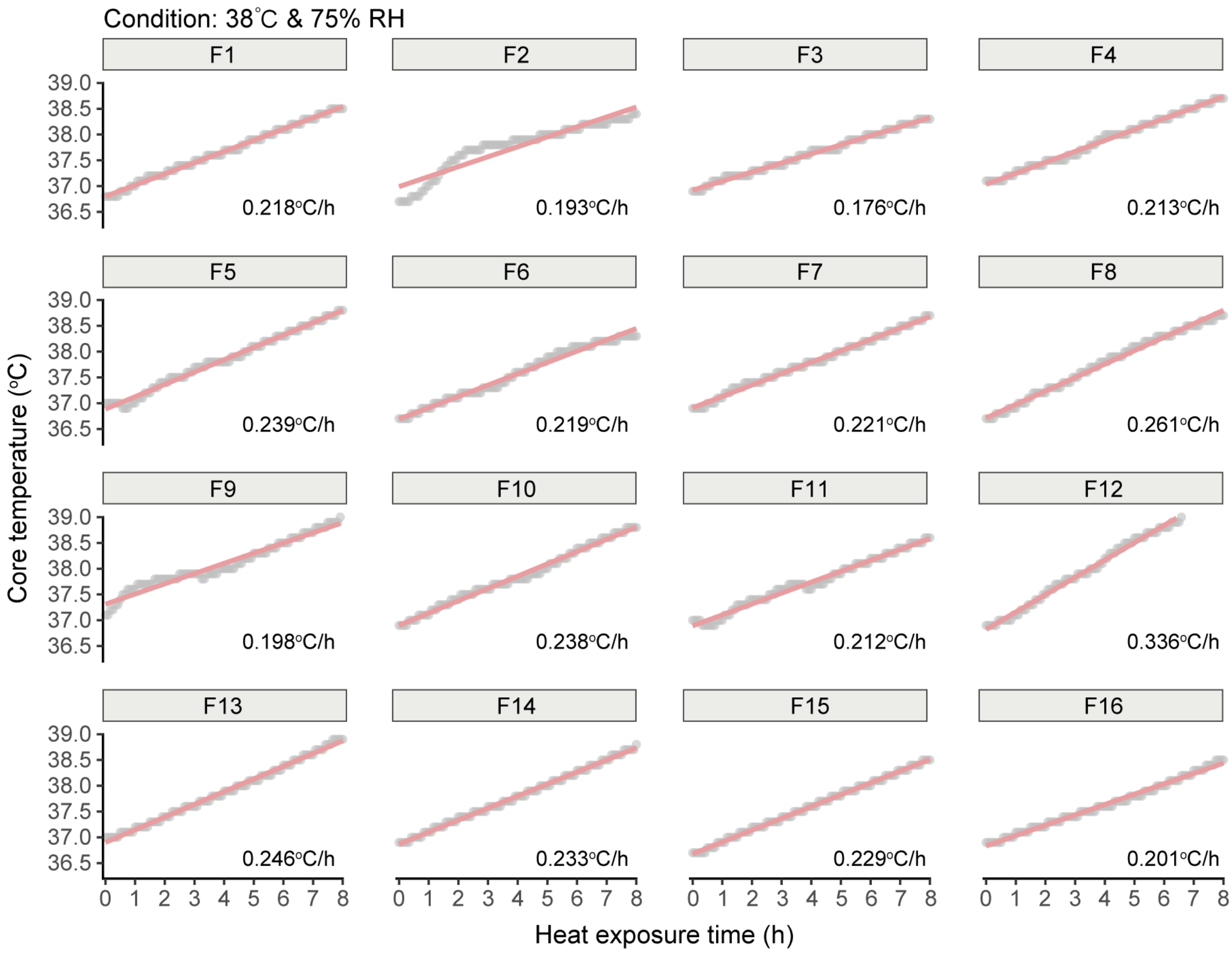


**Supplementary Fig. S17. Individual core temperature trajectories of 16 young adult females during 8-hour heat exposure at 38°C and 75.1% relative humidity.** Core temperature ($T_{core}$) was continuously monitored using a rectal thermistor. The data highlight inter-individual variability in thermoregulatory strain under extremely hot and humid conditions approaching the theoretical survivability threshold.

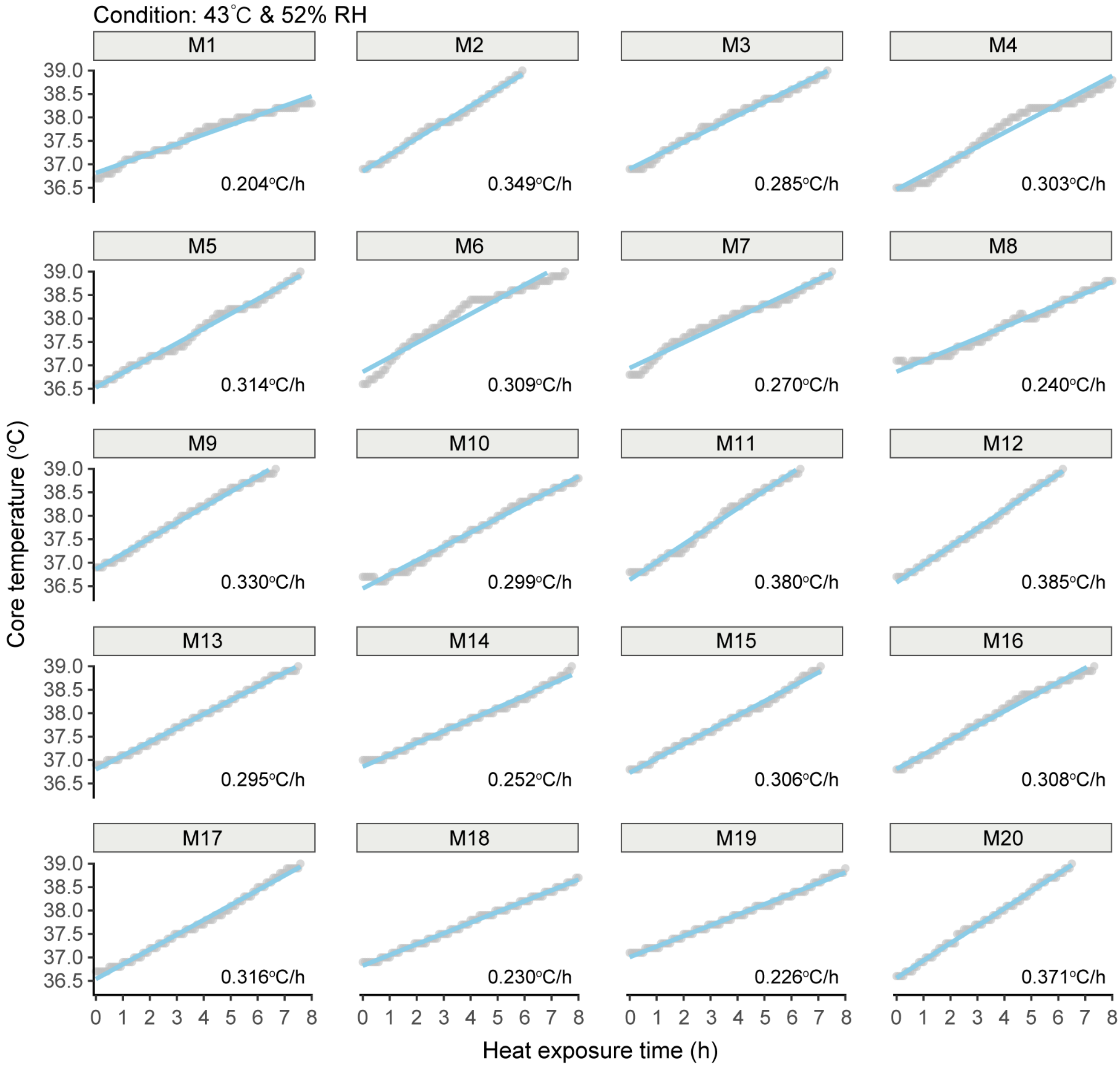


**Supplementary Fig. S18. Individual core temperature trajectories of 20 young adult males during 8-hour heat exposure at 43°C and 52.1% relative humidity.** Core temperature ($T_{core}$) was continuously monitored using a rectal thermistor. The data highlight inter-individual variability in thermoregulatory strain under extremely hot and humid conditions approaching the theoretical survivability threshold.

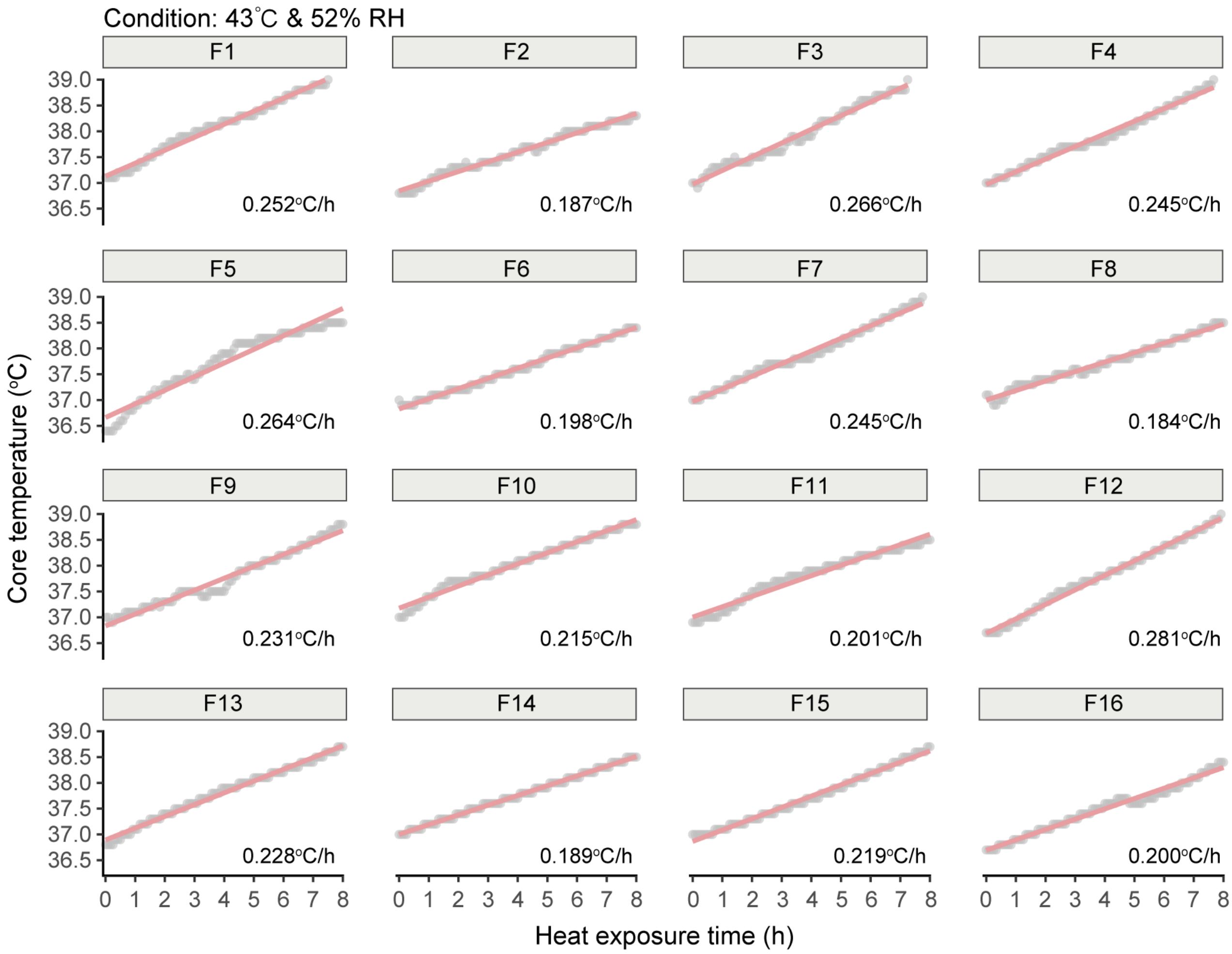


**Supplementary Fig. S19. Individual core temperature trajectories of 16 young adult females during 8-hour heat exposure at 43°C and 52.1% relative humidity.** Core temperature ($T_{core}$) was continuously monitored using a rectal thermistor. The data highlight inter-individual variability in thermoregulatory strain under extremely hot and humid conditions approaching the theoretical survivability threshold.

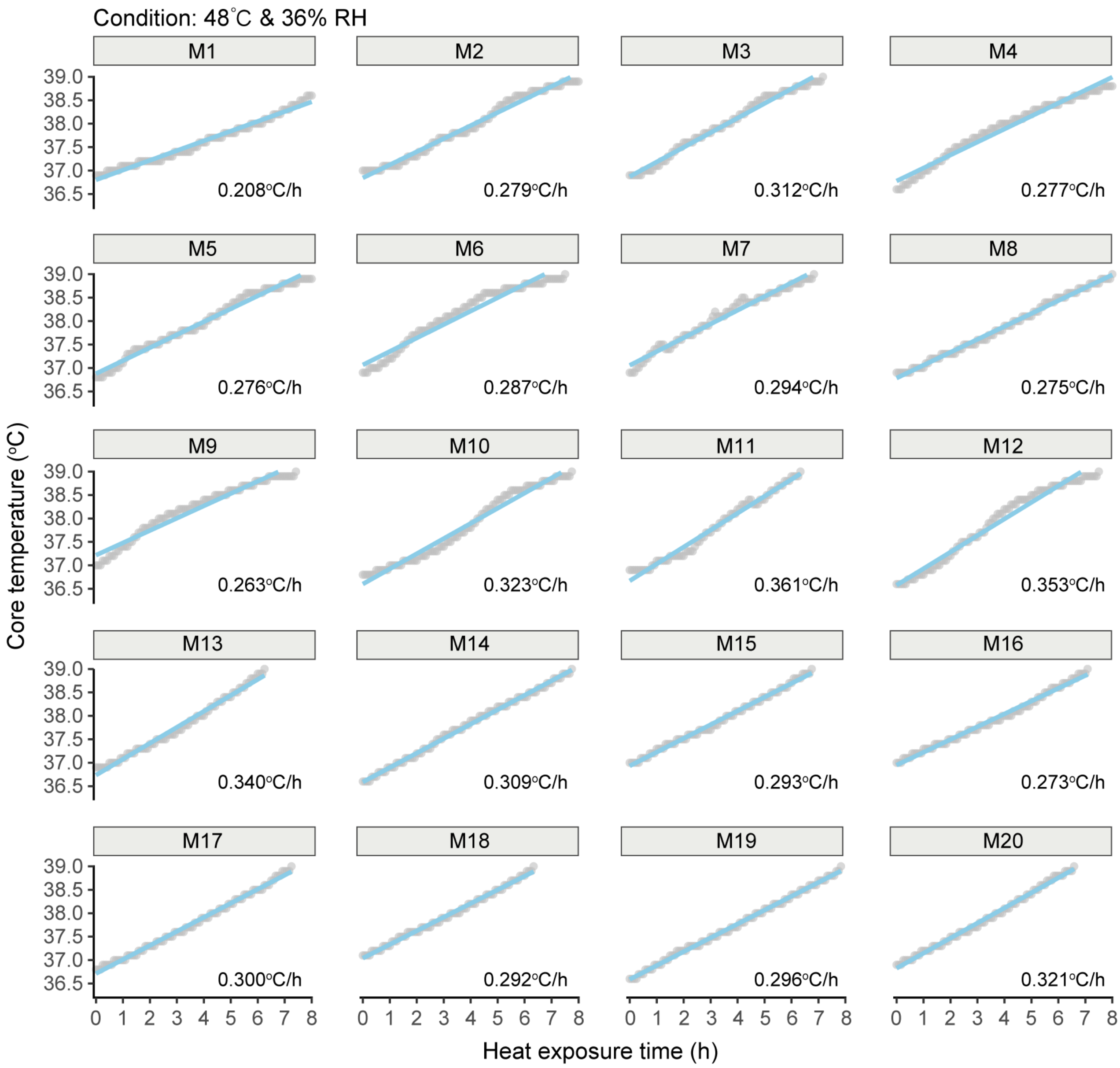


**Supplementary Fig. S20. Individual core temperature trajectories of 20 young adult males during 8-hour heat exposure at 48°C and 36.2% relative humidity.** Core temperature ($T_{core}$) was continuously monitored using a rectal thermistor. The data highlight inter-individual variability in thermoregulatory strain under extremely hot and humid conditions approaching the theoretical survivability threshold.

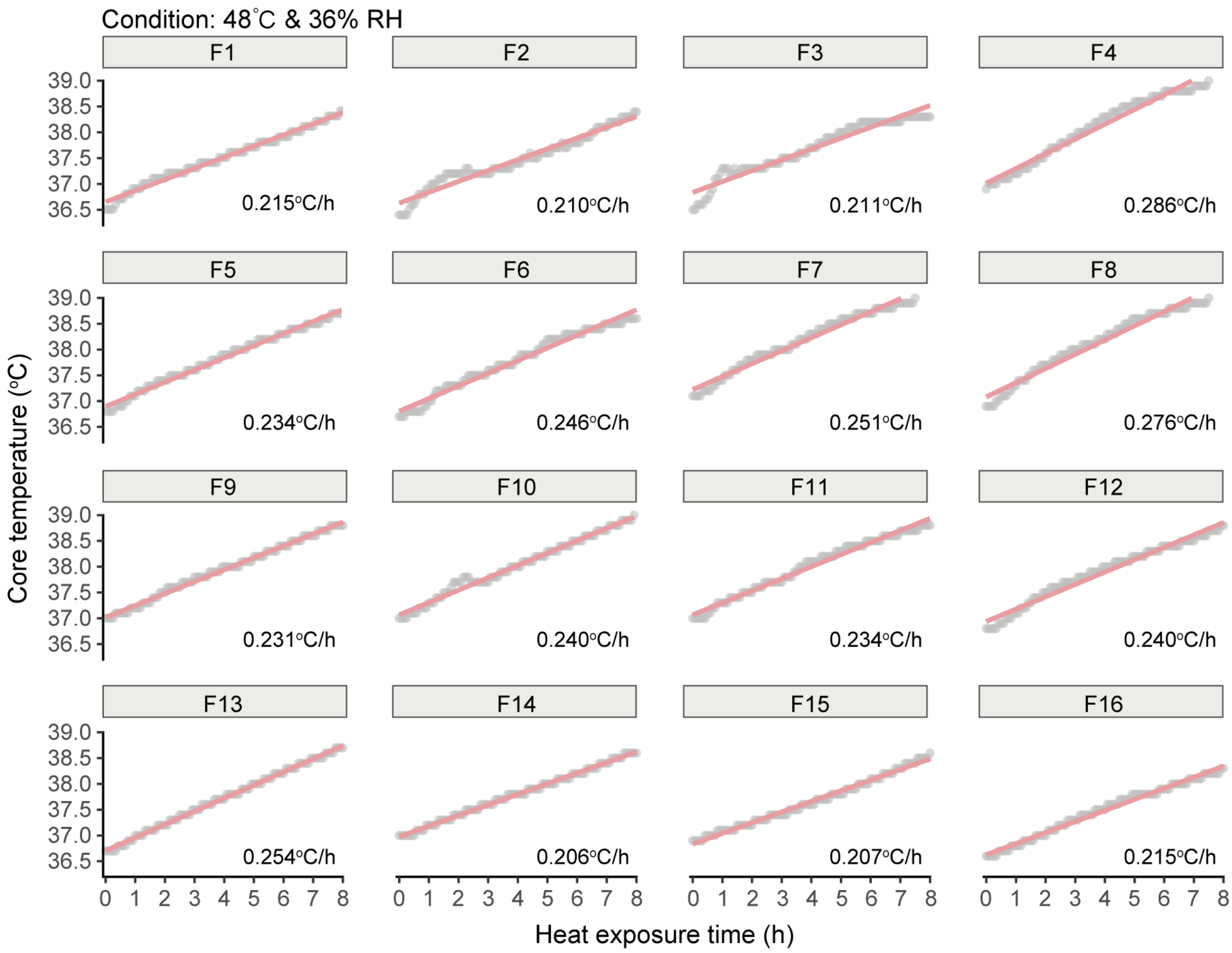


**Supplementary Fig. S21. Individual core temperature trajectories of 16 young adult females during 8-hour heat exposure at 48°C and 36.2% relative humidity.** Core temperature ($T_{core}$) was continuously monitored using a rectal thermistor. The data highlight inter-individual variability in thermoregulatory strain under extremely hot and humid conditions approaching the theoretical survivability threshold.

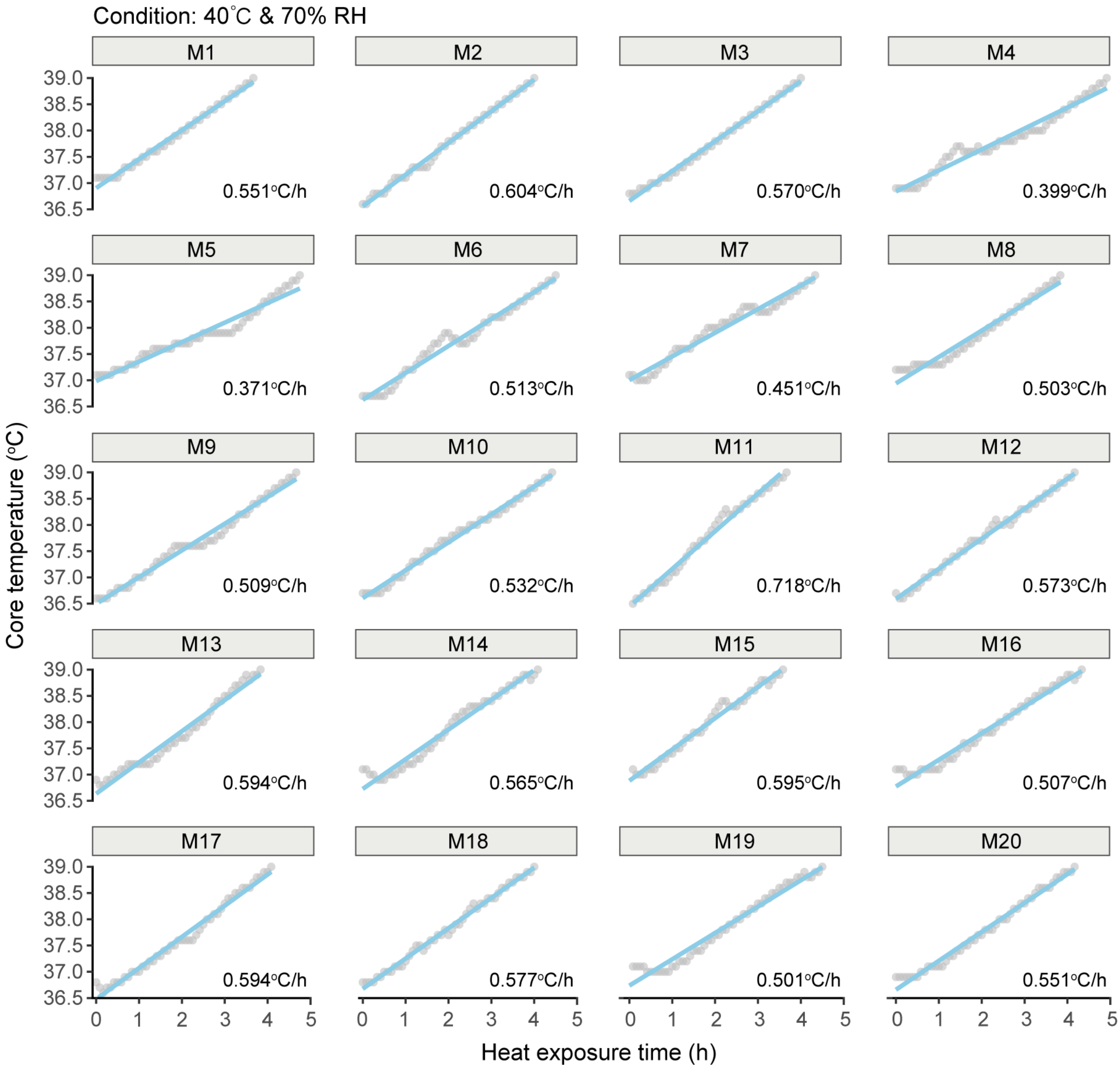


**Supplementary Fig. S22. Individual core temperature trajectories of 20 young adult males during prolonged heat exposure at 40°C and 70.3% relative humidity.** Core temperature ($T_{core}$) was continuously monitored using a rectal thermistor. The data highlight inter-individual variability in thermoregulatory strain under extremely hot and humid conditions approaching the theoretical survivability threshold.

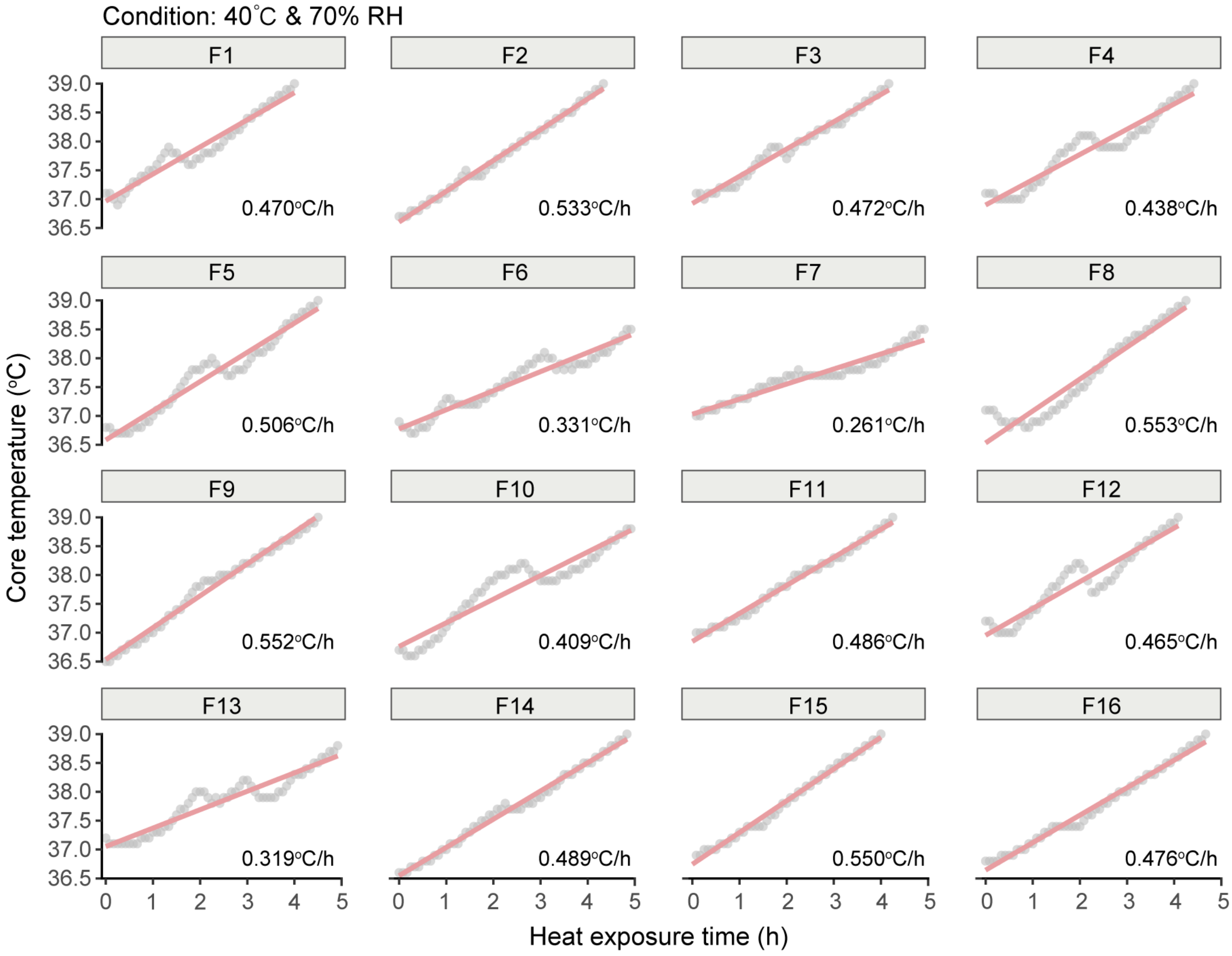


**Supplementary Fig. S23. Individual core temperature trajectories of 16 young adult females during prolonged heat exposure at 40°C and 70.3% relative humidity.** Core temperature ($T_{core}$) was continuously monitored using a rectal thermistor. The data highlight inter-individual variability in thermoregulatory strain under extremely hot and humid conditions approaching the theoretical survivability threshold.

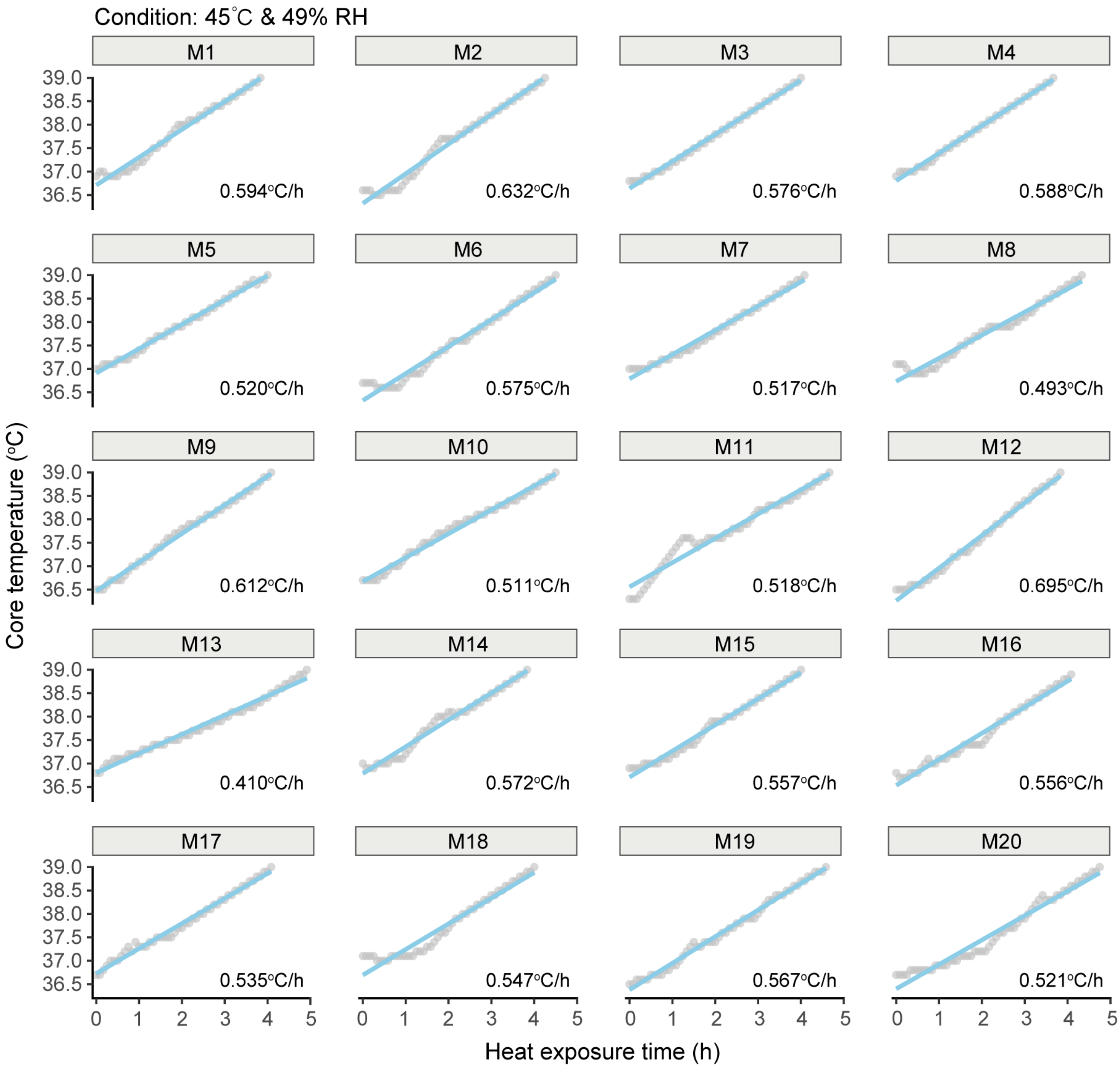


**Supplementary Fig. S24. Individual core temperature trajectories of 20 young adult males during prolonged heat exposure at 45°C and 49% relative humidity.** Core temperature ($T_{core}$) was continuously monitored using a rectal thermistor. The data highlight inter-individual variability in thermoregulatory strain under extremely hot and humid conditions approaching the theoretical survivability threshold.

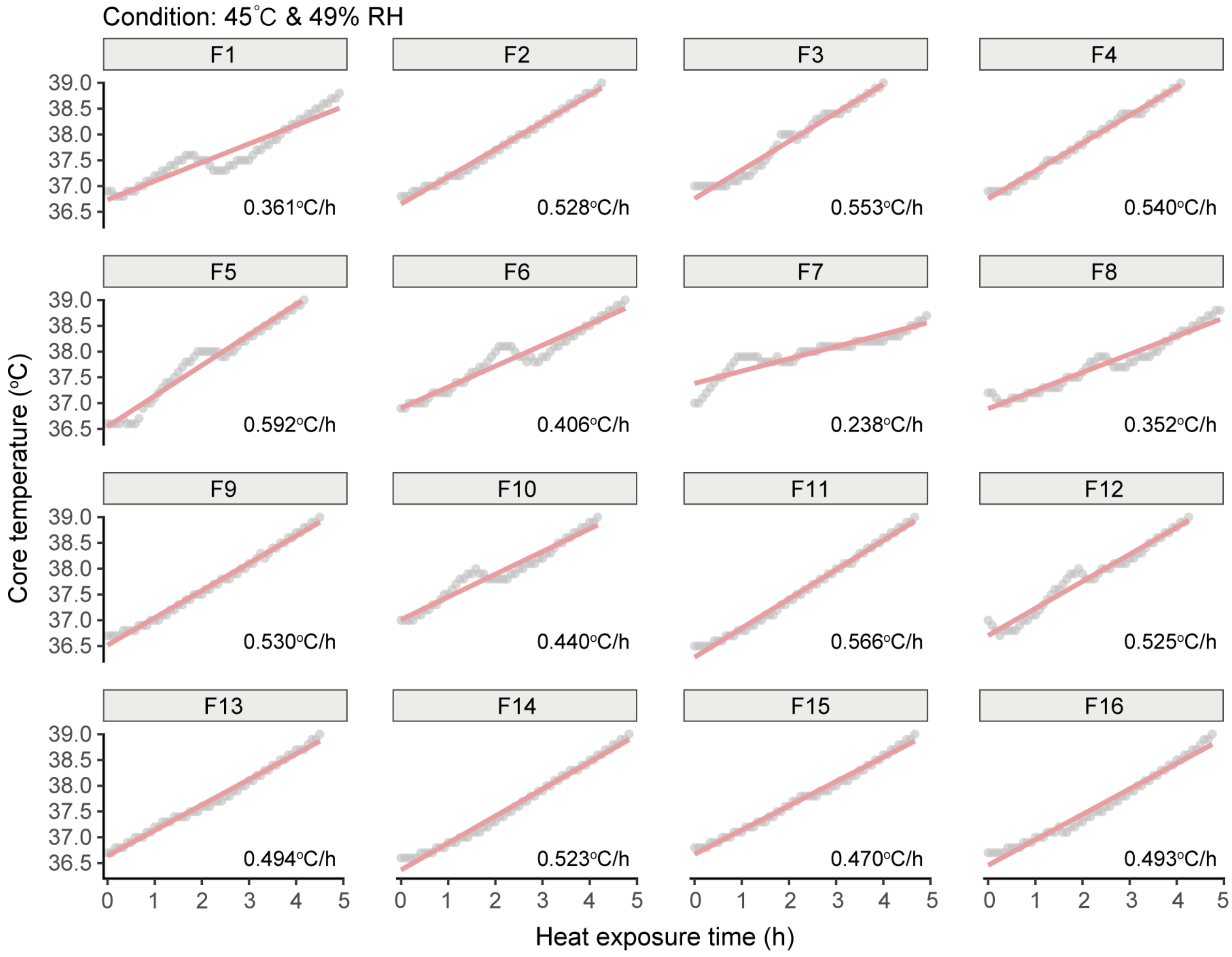


**Supplementary Fig. S25. Individual core temperature trajectories of 16 young adult females during prolonged heat exposure at 45°C and 49% relative humidity.** Core temperature ($T_{core}$) was continuously monitored using a rectal thermistor. The data highlight inter-individual variability in thermoregulatory strain under extremely hot and humid conditions approaching the theoretical survivability threshold.

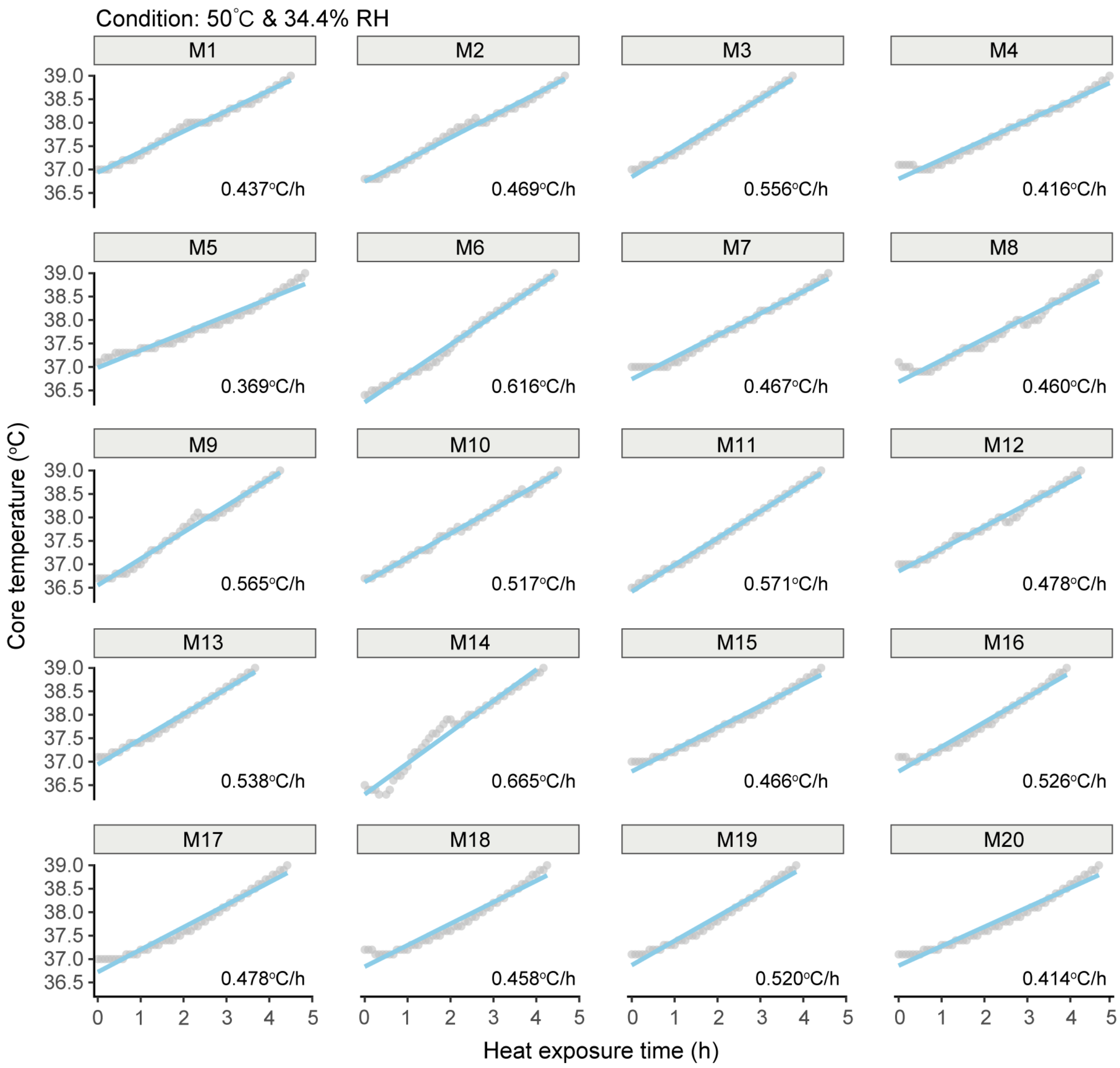


**Supplementary Fig. S26. Individual core temperature trajectories of 20 young adult males during prolonged heat exposure at 50°C and 34.4% relative humidity.** Core temperature ($T_{core}$) was continuously monitored using a rectal thermistor. The data highlight inter-individual variability in thermoregulatory strain under extremely hot and humid conditions approaching the theoretical survivability threshold.

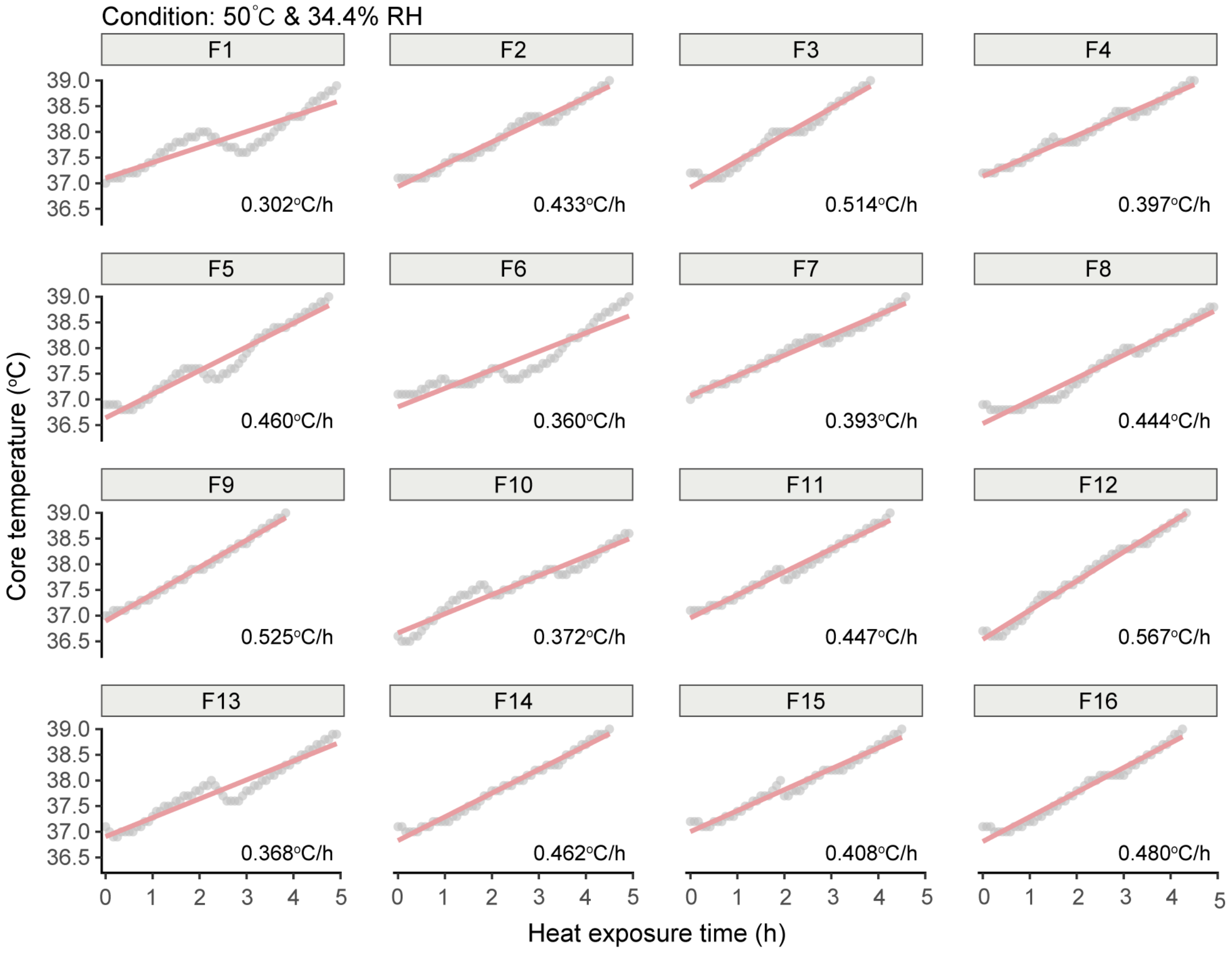


**Supplementary Fig. S27. Individual core temperature trajectories of 16 young adult females during prolonged heat exposure at 50°C and 34.4% relative humidity.** Core temperature ($T_{core}$) was continuously monitored using a rectal thermistor. The data highlight inter-individual variability in thermoregulatory strain under extremely hot and humid conditions approaching the theoretical survivability threshold.

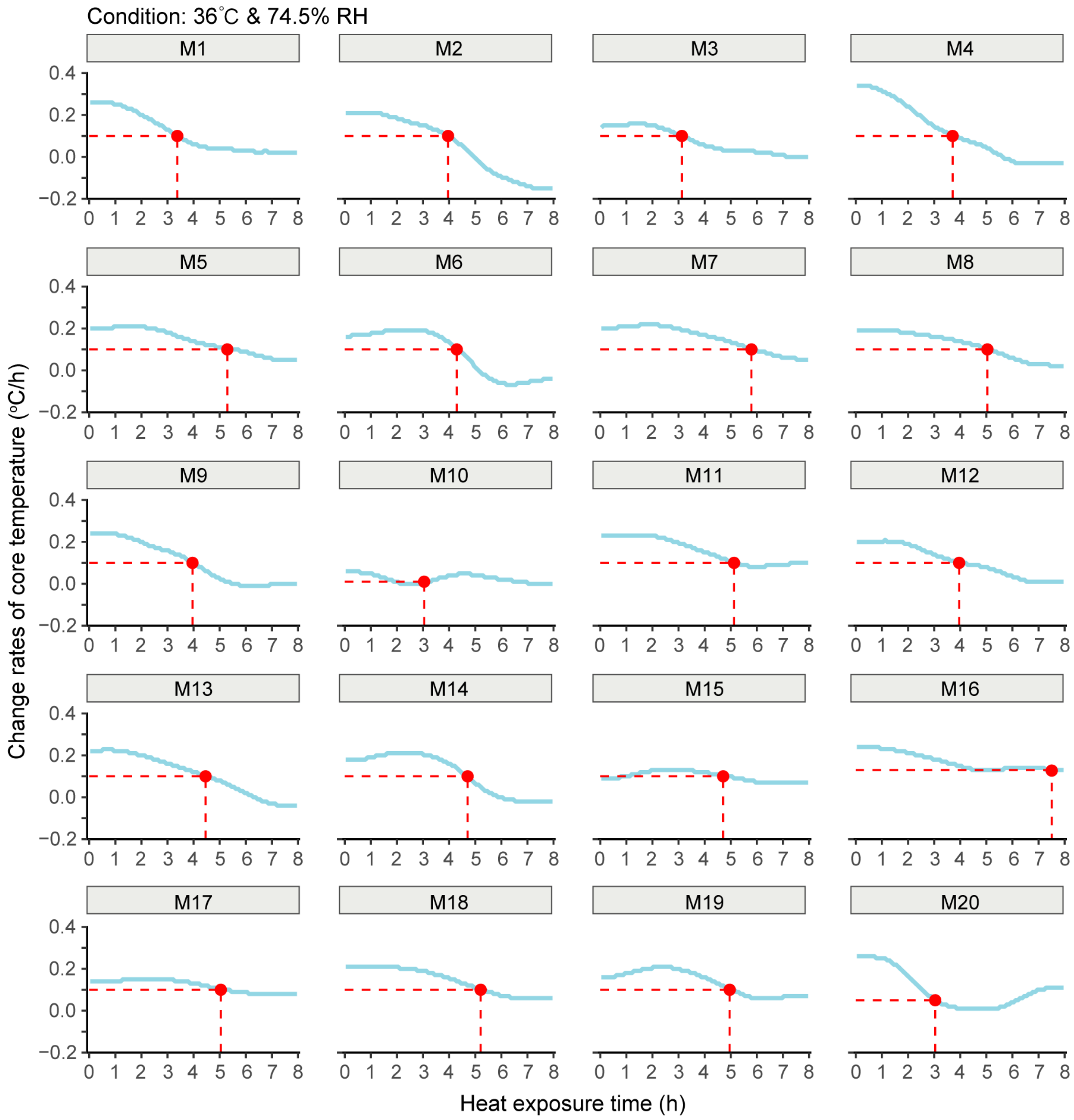


**Supplementary Fig. S28. Individual rates of core temperature change during 8-hour exposure at 36°C and 74.5% relative humidity in 20 young adult males.** Core temperature ($T_{core}$) change rates were calculated in 5-minute intervals using smoothed individual trajectories. Despite identical environmental conditions, notable inter-individual variability was observed in thermoregulatory response profiles.

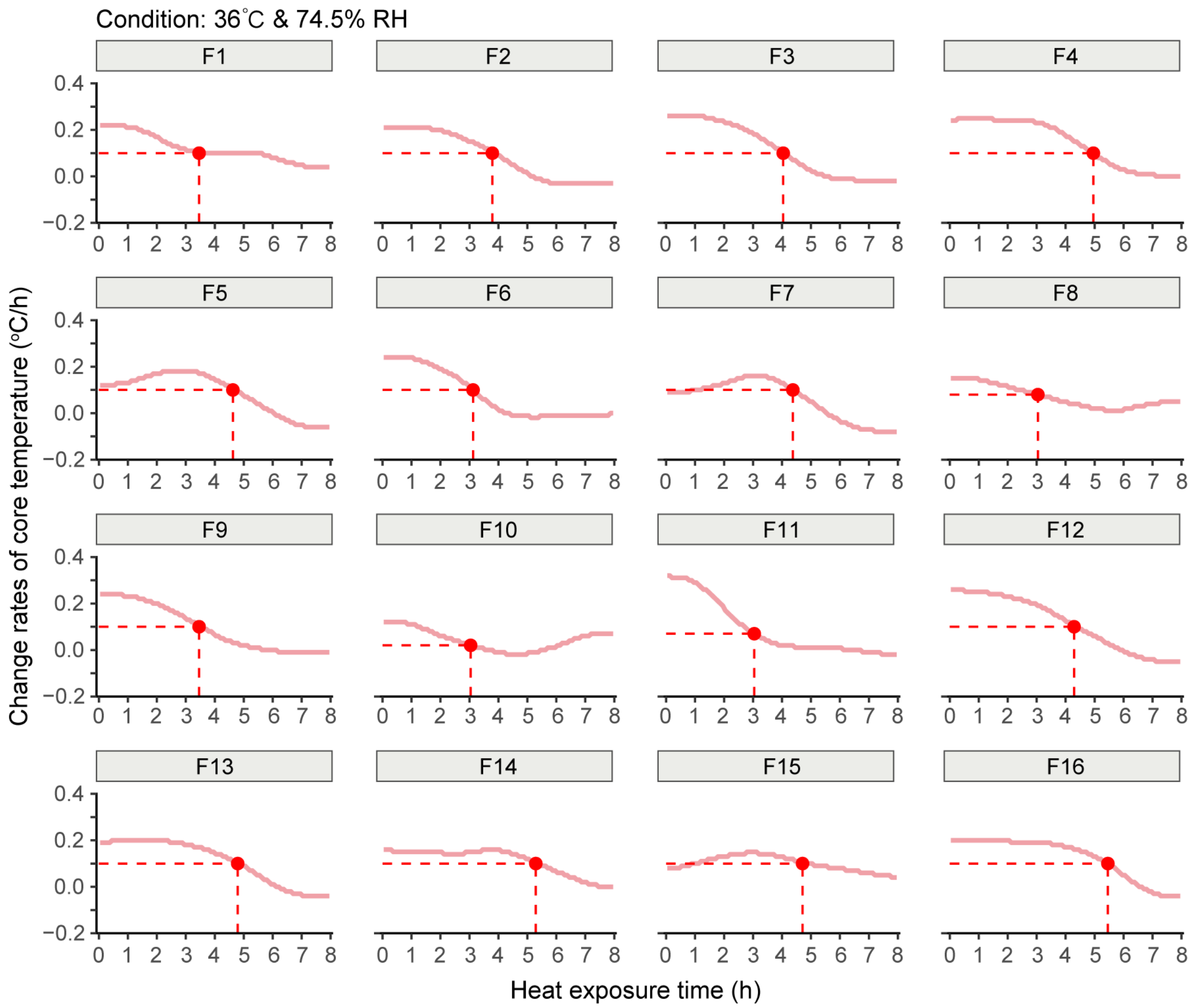


**Supplementary Fig. S29. Individual rates of core temperature change during 8-hour exposure at 36°C and 74.5% relative humidity in 16 young adult females.** Core temperature ($T_{core}$) change rates were calculated in 5-minute intervals using smoothed individual trajectories. Despite identical environmental conditions, notable inter-individual variability was observed in thermoregulatory response profiles.

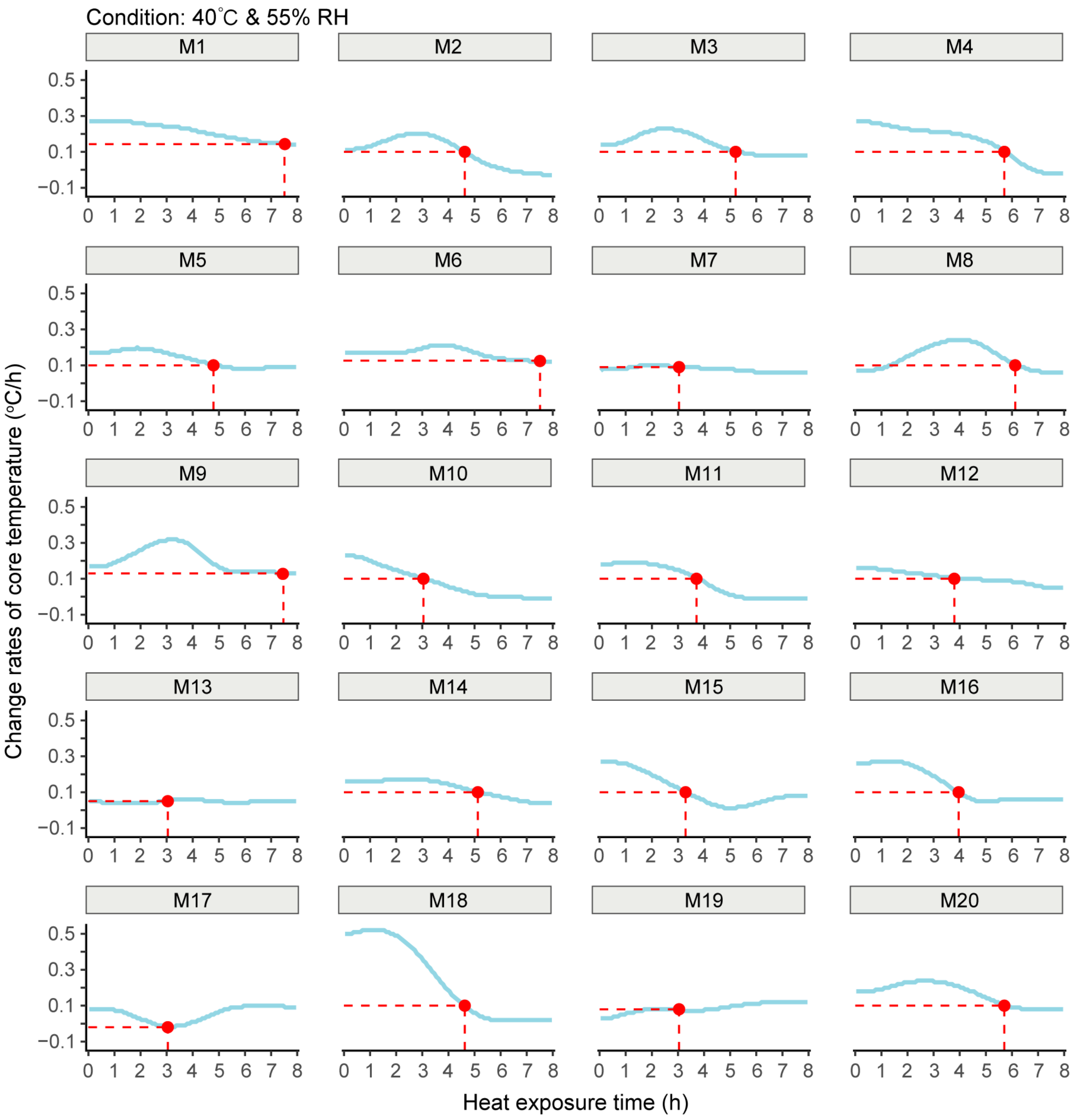


**Supplementary Fig. S30. Individual rates of core temperature change during 8-hour exposure at 40°C and 55% relative humidity in 20 young adult males.** Core temperature ($T_{core}$) change rates were calculated in 5-minute intervals using smoothed individual trajectories. Despite identical environmental conditions, notable inter-individual variability was observed in thermoregulatory response profiles.

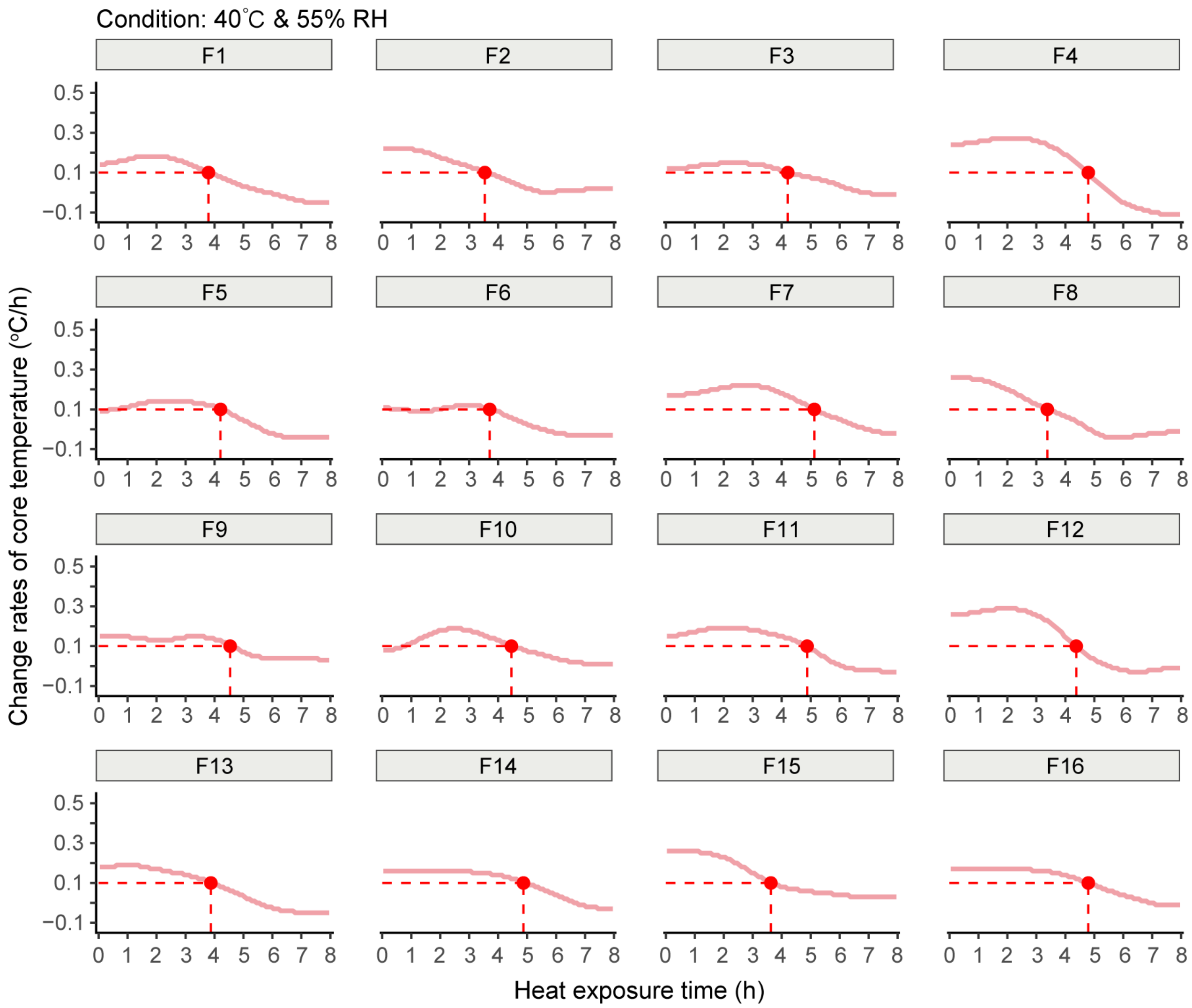


**Supplementary Fig. S31. Individual rates of core temperature change during 8-hour exposure at 40°C and 55% relative humidity in 16 young adult females.** Core temperature ($T_{core}$) change rates were calculated in 5-minute intervals using smoothed individual trajectories. Despite identical environmental conditions, notable inter-individual variability was observed in thermoregulatory response profiles.

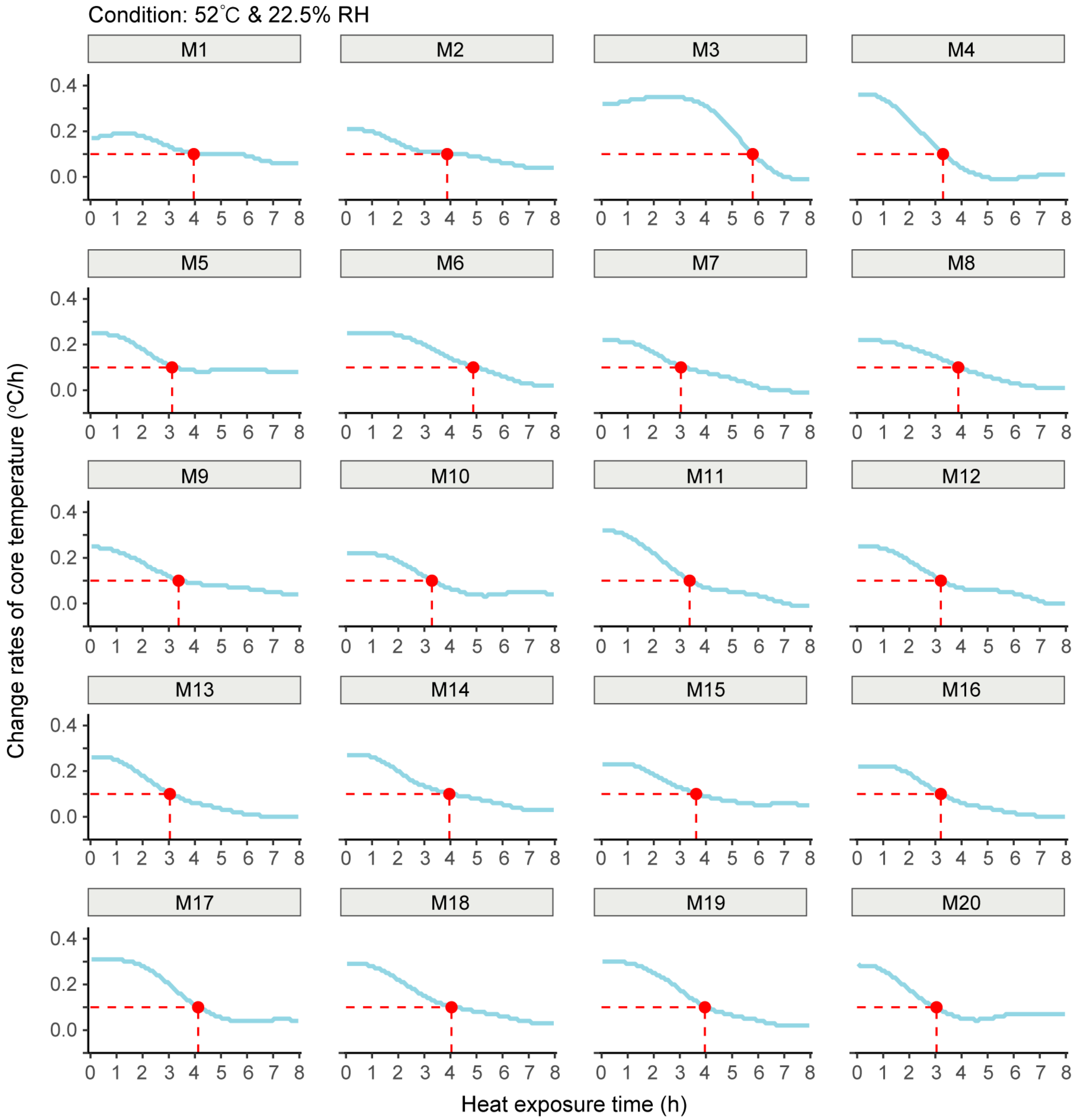


**Supplementary Fig. S32. Individual rates of core temperature change during 8-hour exposure at 52°C and 22.5% relative humidity in 20 young adult males.** Core temperature ($T_{core}$) change rates were calculated in 5-minute intervals using smoothed individual trajectories. Despite identical environmental conditions, notable inter-individual variability was observed in thermoregulatory response profiles.

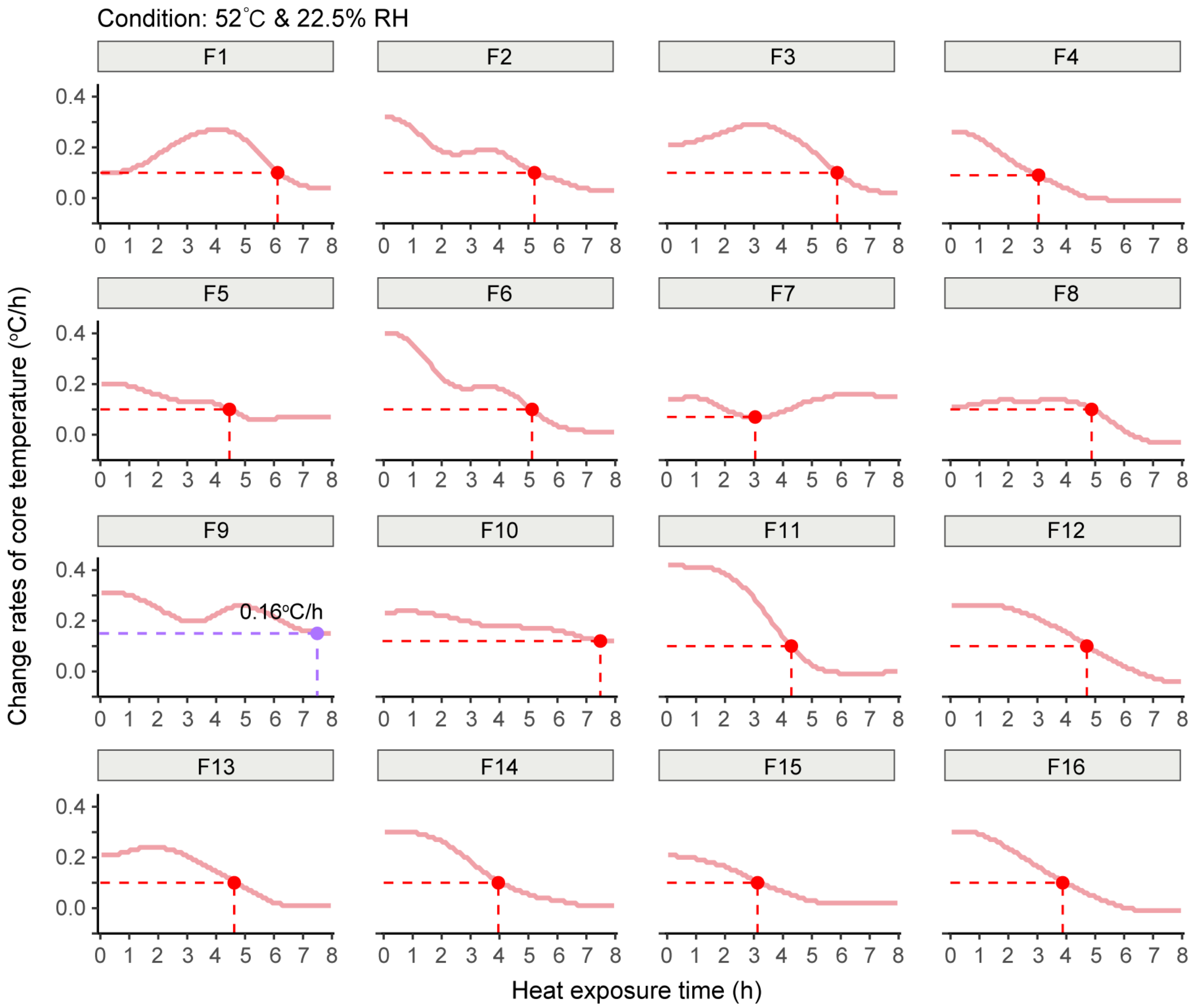


**Supplementary Fig. S33. Individual rates of core temperature change during 8-hour exposure at 52°C and 22.5% relative humidity in 16 young adult females.** Core temperature ($T_{core}$) change rates were calculated in 5-minute intervals using smoothed individual trajectories. Despite identical environmental conditions, notable inter-individual variability was observed in thermoregulatory response profiles.

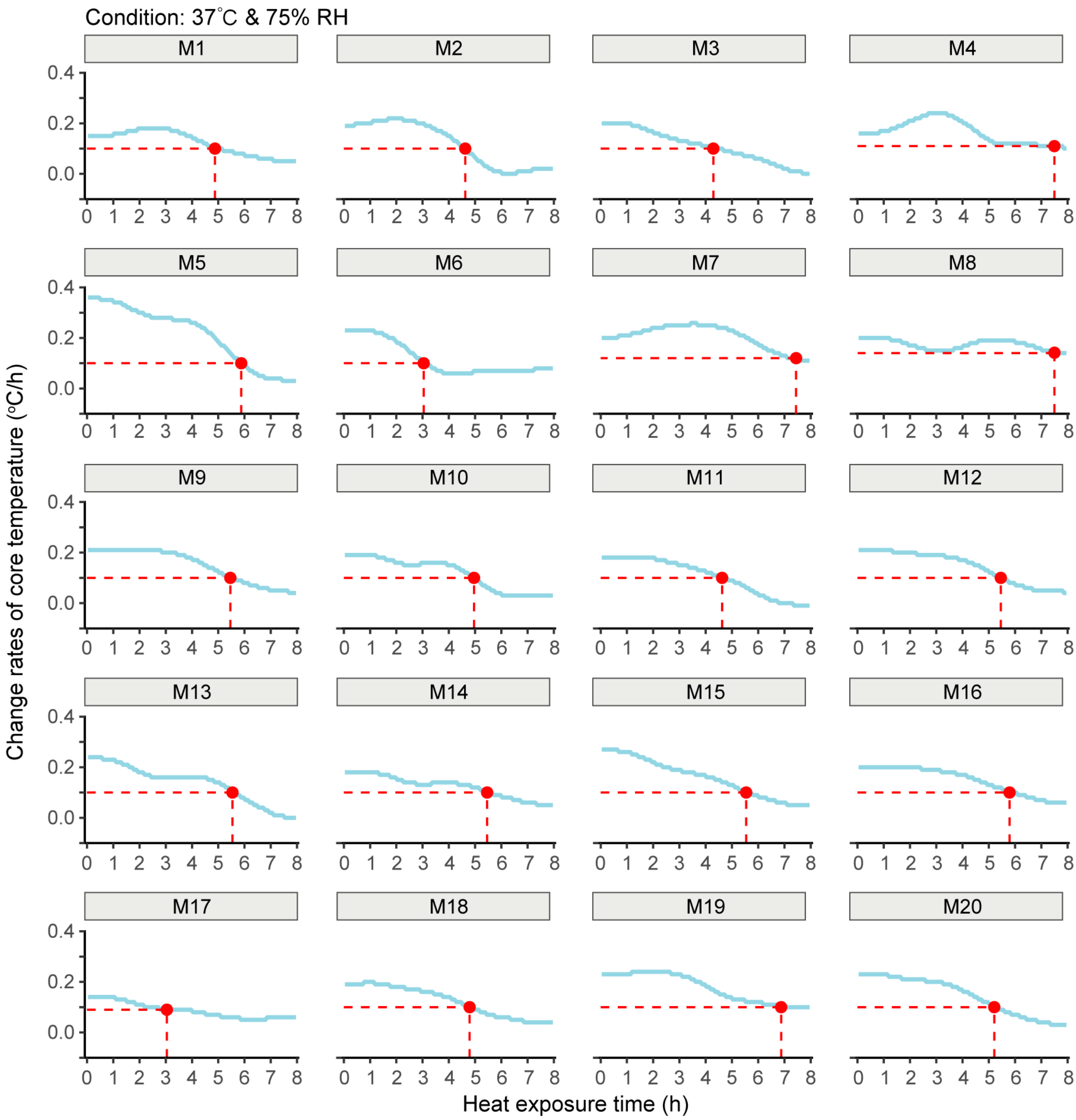


**Supplementary Fig. S34. Individual rates of core temperature change during 8-hour exposure at 37°C and 74.8% relative humidity in 20 young adult males.** Core temperature ($T_{core}$) change rates were calculated in 5-minute intervals using smoothed individual trajectories. Despite identical environmental conditions, notable inter-individual variability was observed in thermoregulatory response profiles.

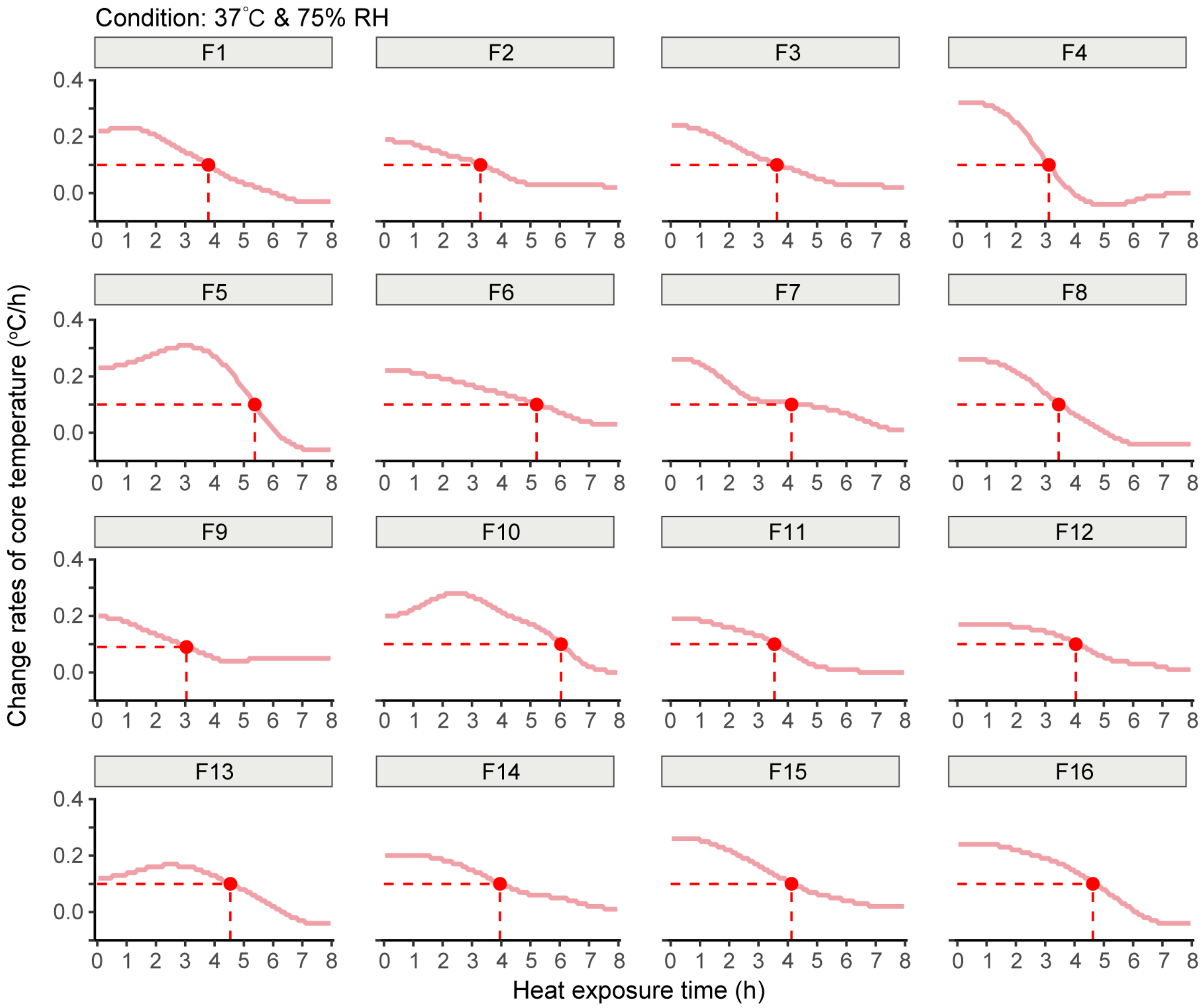


**Supplementary Fig. S35. Individual rates of core temperature change during 8-hour exposure at 37°C and 74.8% relative humidity in 16 young adult females.** Core temperature ($T_{core}$) change rates were calculated in 5-minute intervals using smoothed individual trajectories. Despite identical environmental conditions, notable inter-individual variability was observed in thermoregulatory response profiles.

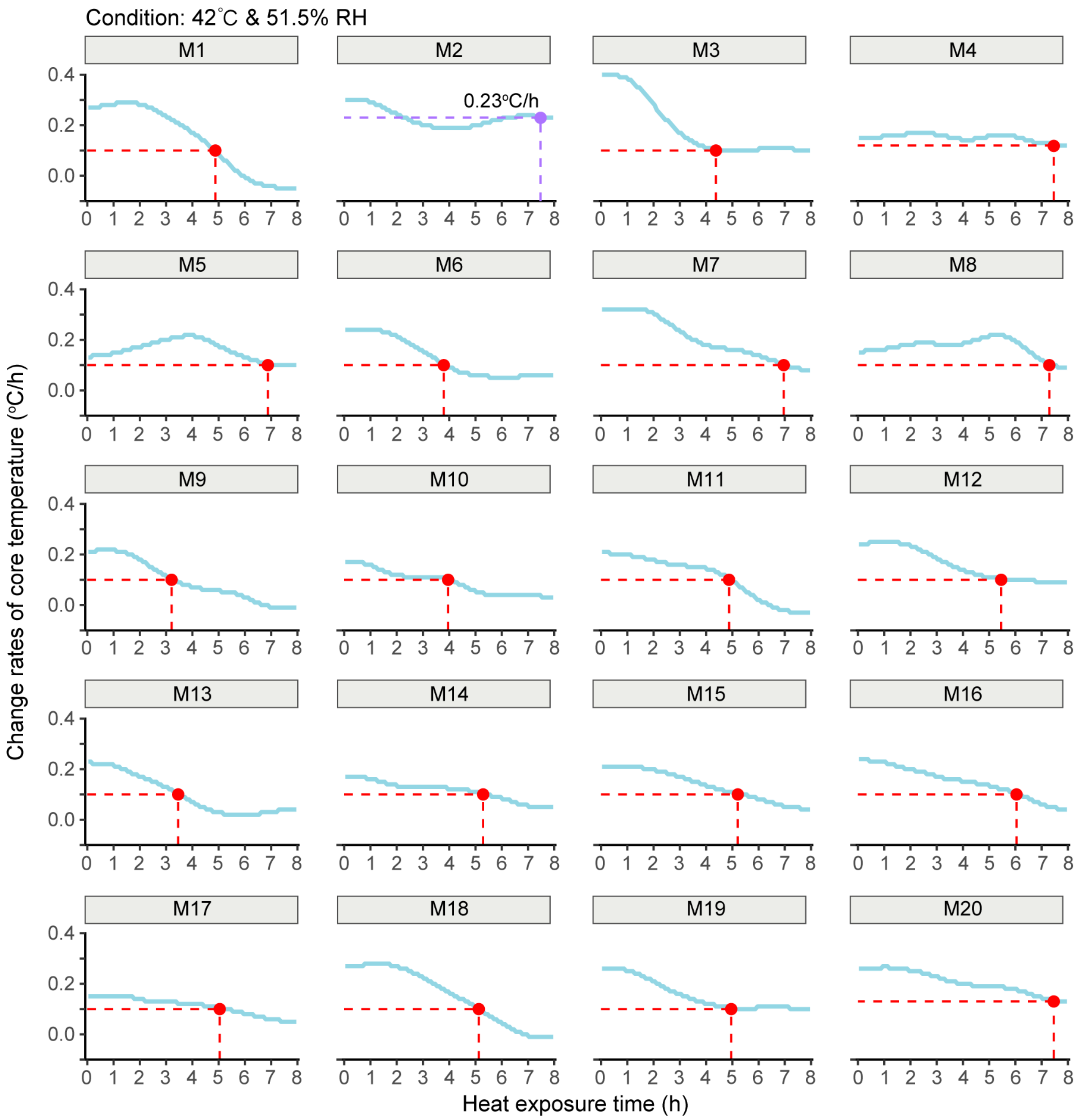


**Supplementary Fig. S36. Individual rates of core temperature change during 8-hour exposure at 42°C and 51.5% relative humidity in 20 young adult males.** Core temperature ($T_{core}$) change rates were calculated in 5-minute intervals using smoothed individual trajectories. Despite identical environmental conditions, notable inter-individual variability was observed in thermoregulatory response profiles.

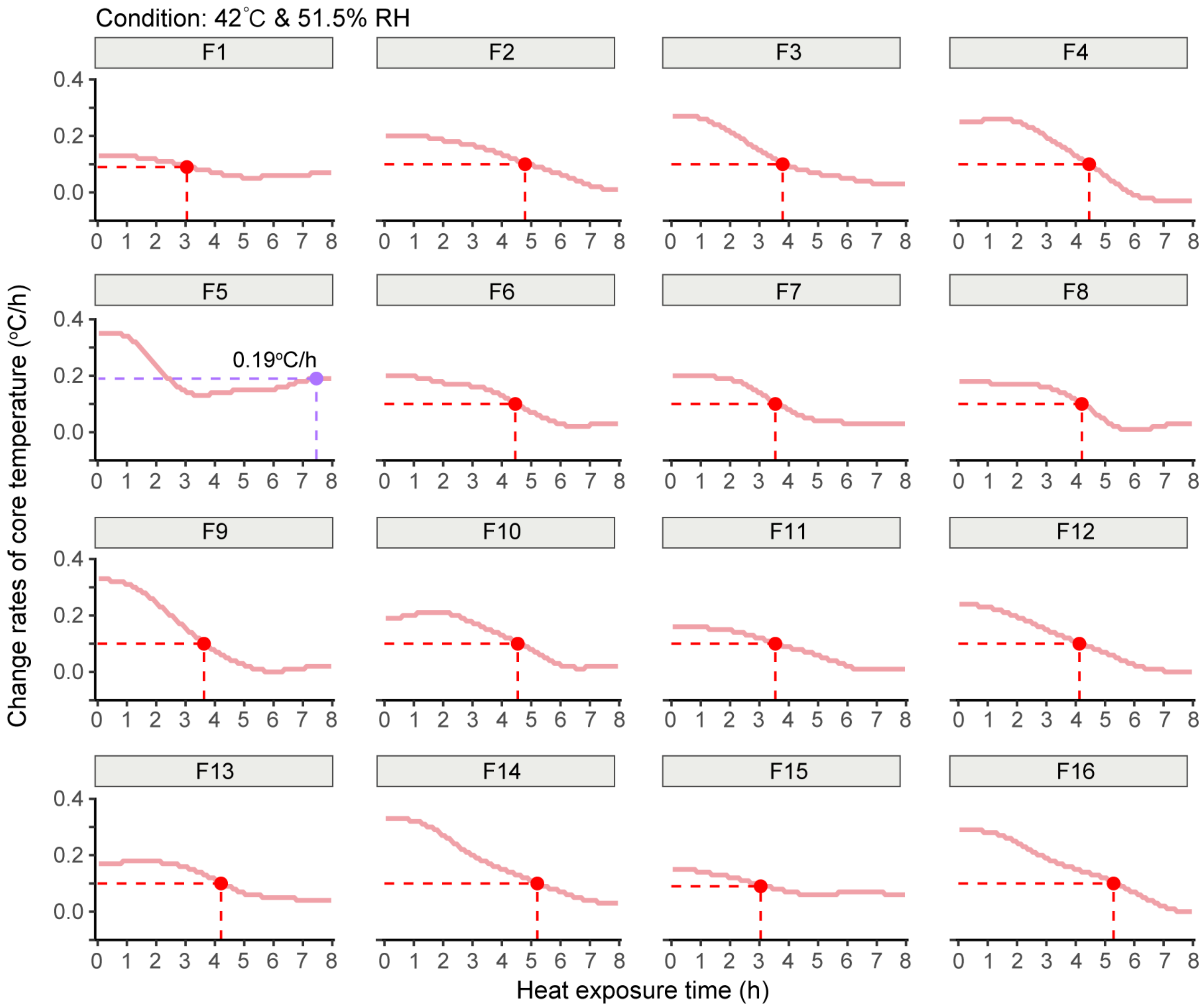


**Supplementary Fig. S37. Individual rates of core temperature change during 8-hour exposure at 42°C and 51.5% relative humidity in 16 young adult females.** Core temperature ($T_{core}$) change rates were calculated in 5-minute intervals using smoothed individual trajectories. Despite identical environmental conditions, notable inter-individual variability was observed in thermoregulatory response profiles.

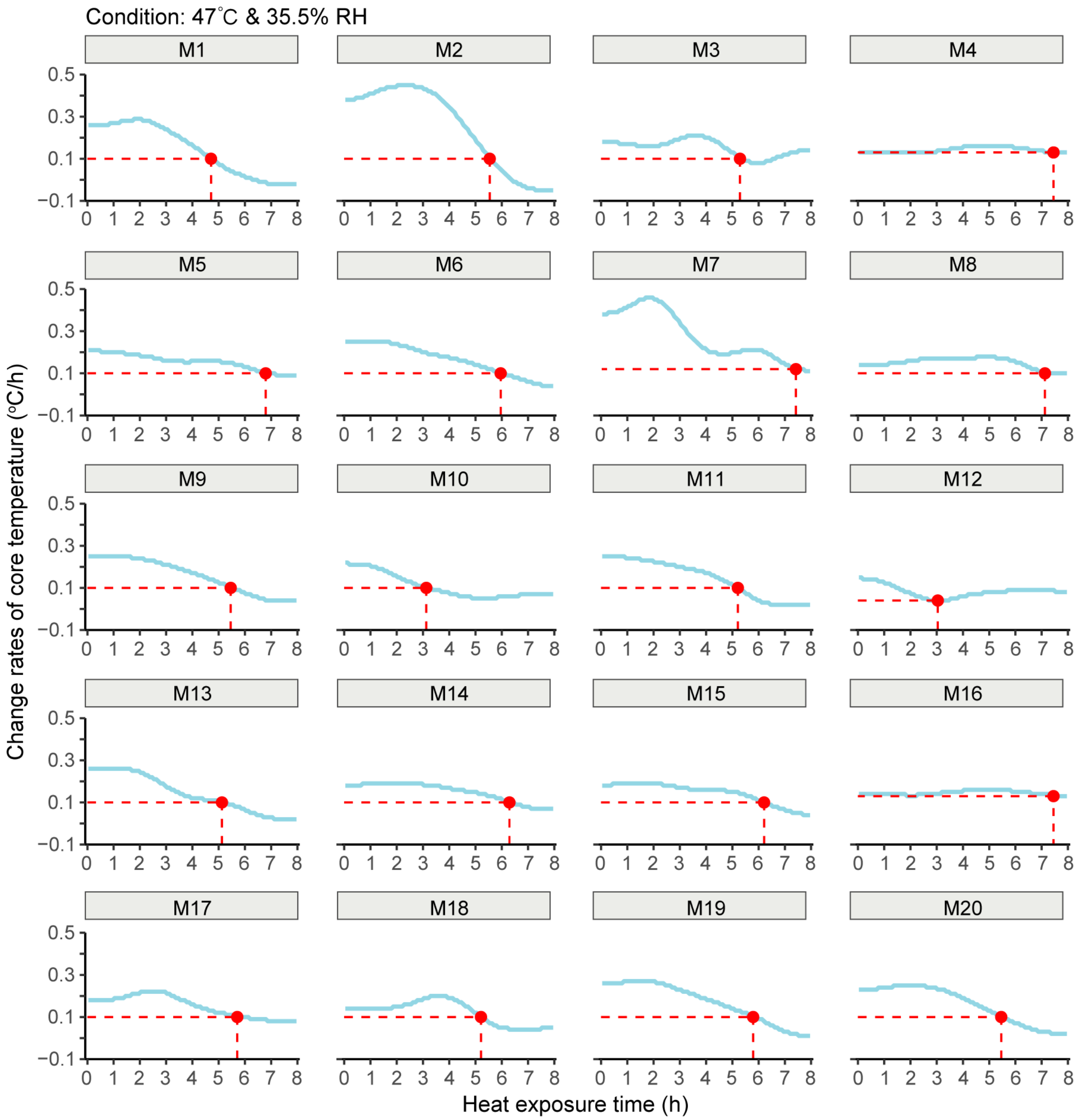


**Supplementary Fig. S38. Individual rates of core temperature change during 8-hour exposure at 47°C and 35.5% relative humidity in 20 young adult males.** Core temperature ($T_{core}$) change rates were calculated in 5-minute intervals using smoothed individual trajectories. Despite identical environmental conditions, notable inter-individual variability was observed in thermoregulatory response profiles.

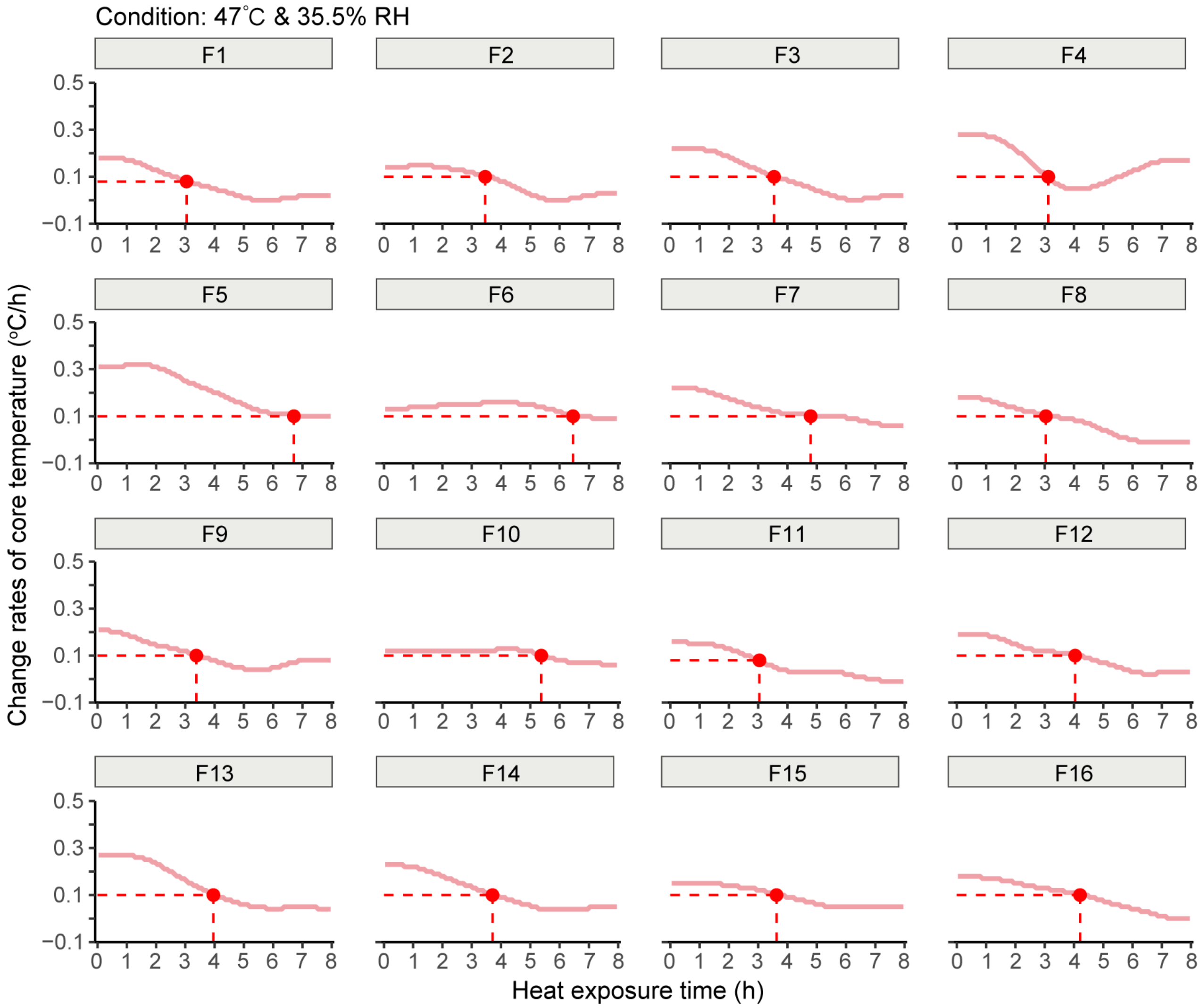


**Supplementary Fig. S39. Individual rates of core temperature change during 8-hour exposure at 47°C and 35.5% relative humidity in 16 young adult females.** Core temperature ($T_{core}$) change rates were calculated in 5-minute intervals using smoothed individual trajectories. Despite identical environmental conditions, notable inter-individual variability was observed in thermoregulatory response profiles.

**Supplementary Table S1. Overview of studies reporting critical environmental limits (CEL) and thresholds for compensable and uncompensable heat stress.** This table summarizes experimental and modeling studies that define CEL for heat exposure. CEL is identified as the psychrometric threshold defined by dry-bulb temperature ($T_{db}$), relative humidity (RH), or partial water vapor pressure ($P_a$), above which heat production exceeds heat dissipation, leading to uncompensable heat stress. The table distinguishes between studies using stepwise heat exposure protocols, biophysical modeling, and advanced human thermoregulation modeling. Information on sample size, sex inclusion, exposure duration, environmental control precision, and definitions of compensability are included to highlight methodological variability and reliability. M, males; F, females; N, total number of participants (exact sex distribution unspecified); BMI, body mass index; BSA, body surface area; $I_{cl}$, clothing insulation; $P_a$, water vapor pressure (mmHg); RH, relative humidity (%); $T_{core}$, core temperature (°C); $T_{db}$, dry-bulb temperature (°C); $T_w$, wet-bulb temperature (°C); One metabolic equivalent (MET) is defined as 58.15 watts per square meter (W/m$^2$)41. *CEL and sweat rates are reported as mean ± standard deviation (SD) or mean [95% confidence interval (CI)]; NA: data not available.

| Authors (Year) | Subject | Age (yr) | BMI (kg/m$^2$) | BSA (m$^2$) | $I_{cl}$ (clo) | Activity | Environmental condition | Protocol | Metabolic rate (METs) | Sweat rate (g/m$^2$/h) | Critical environmental limit (CEL) |
|---|---|---|---|---|---|---|---|---|---|---|---|
| Kenney & Zeman (2002)[33] | 11M | 22±1 | 26±1 | 2.08±0.05 | 0.27 | Treadmill walking at 30% $VO_2$max (2.5 h) | 34°C (constant $T_{db}$) | Ramp (30 min stabilization at 9 mmHg or 28°C; $P_a$ or $T_{db}$ increased stepwise by 1mmHg or 1 °C every 5 min) | 3.28±0.10 | 306±10 | $T_w$=29.5°C ($T_{db}$=34°C & 71.2% RH) |
| | | | | | | | 36°C (constant $T_{db}$) | | | 357±16 | $T_w$=28.8°C ($T_{db}$=36°C & 57.0% RH) |
| | | | | | | | 38°C (constant $T_{db}$) | | | 394±14 | $T_w$=28.0°C ($T_{db}$=38°C & 44.6% RH) |
| | | | | | | | 12 mmHg (constant $P_a$) | | | 318±16 | $T_w$=24.3°C ($T_{db}$=43.6°C & 17.9% RH) |
| | | | | | | | 16 mmHg (constant $P_a$) | | | 279±19 | $T_w$=26.2°C ($T_{db}$=42.1°C & 25.8% RH) |
| | | | | | | | 20 mmHg (constant $P_a$) | | | 354±24 | $T_w$=28.0°C ($T_{db}$=41.2°C & 33.9% RH) |
| | 10F | 20±1 | 24±2 | 1.62±0.04 | 0.27 | Treadmill walking at 30% $VO_2$max (2.5 h) | 34°C (constant $T_{db}$) | Ramp (30 min stabilization at 9 mmHg or 28°C; $P_a$ or $T_{db}$ increased stepwise by 1mmHg or 1 °C every 5 min) | 2.39±0.05 | 262±38 | $T_w$=31.5°C ($T_{db}$=34°C & 83.5% RH) |
| | | | | | | | 36°C (constant $T_{db}$) | | | 257±17 | $T_w$=31.2°C ($T_{db}$=36°C & 69.9% RH) |
| | | | | | | | 38°C (constant $T_{db}$) | | | 278±14 | $T_w$=30.1°C ($T_{db}$=38°C & 54.3% RH) |
| | | | | | | | 12 mmHg (constant $P_a$) | | | 250±14 | $T_w$=24.8°C ($T_{db}$=45.3°C & 16.4% RH) |
| | | | | | | | 16 mmHg (constant $P_a$) | | | 243±13 | $T_w$=26.3°C ($T_{db}$=42.4°C & 25.4% RH) |
| | | | | | | | 20 mmHg (constant $P_a$) | | | 248±20 | $T_w$=28.0°C ($T_{db}$=41.4°C & 33.6% RH) |
| Schlader et al. (2020)[34] | 15M | 23±3 | 26.4 | 2.00±0.30 | 0.47 | Seated on mesh chair, resting quietly | 32.1±0.1°C & 95±2% RH | Prolonged 8-hour exposure, passive heating | 1.11±0.03 | 145±62 | Uncompensable at $T_w$=34.4±0.5°C ($T_{db}$=35.0±0.1°C & 96±2% RH) |
| | | | | | | | 33.1±0.2°C & 94±2% RH | | 1.17±0.06 | 151±65 | |
| | | | | | | | 35.0±0.1°C & 96±2% RH | | 1.29±0.06 | 138±67 | |
| Wolf et al. (2021)[35] | 8M | 23±4 | 23±3 | 1.86±0.20 | 0.47 | Resting (seated, awake) | 38.0°C & 40% RH | Ramp (30 min stabilization at 38°C & 40% RH or 38°C & 9mmHg; *RH* or $P_a$ increased stepwise by 10% or 1mmHg every 5 min) | 0.75±0.04 | 211±73 | $T_w$=33.7°C ($T_{db}$=38.0±0.2°C & 73.8±6.5% RH) |
| | 8F | | | | | | | | 0.76±0.05 | 163±29 | $T_w$=34.6°C ($T_{db}$=37.9±0.2°C & 79.4±7.4% RH) |
| | 7M | | | | 0.47 | Cycling at 10 W | 38.0°C & 40% RH | | 2.29±0.15 | 269±46 | $T_w$=32.3°C ($T_{db}$=38.2±0.3°C & 65.0±3.2% RH) |

| Authors (Year) | Subject | Age (yr) | BMI (kg/m²) | BSA (m²) | $I_{cl}$ (clo) | Activity | Environmental condition | Protocol | Metabolic rate (METs) | Sweat rate (g/m²/h) | Critical environmental limit (CEL) |
|---|---|---|---|---|---|---|---|---|---|---|---|
| | 8F | | | | | | | | 2.48±0.24 | 181±49 | $T_w$=29.4°C ($T_{db}$=38.1±0.2°C & 50.6±11.9% RH) |
| | 9M | | | | 0.27 | 30% $VO_2$max | 38.0°C & 9 mmHg | | 3.25±0.35 | 244±37 | $T_w$=28.0°C ($T_{db}$=38.1±0.5°C & 44.1±6.3% RH) |
| | 9F | | | | | | | | 2.39±0.17 | 170±33 | $T_w$=30.0°C ($T_{db}$=38.1±0.4°C & 53.8±7.1% RH) |
| Vecellio et al. (2022)[20] | 3M & 5F | 24±4 | 23.7 | 1.84±0.20 | 0.27 | Cycling at 10 W | 36°C (constant $T_{db}$) | Ramp (30 min stabilization; $P_a$ or $T_{db}$ increased stepwise by 1 mmHg or 1 °C every 5 min) | NA | 98±65 | $T_w$=30.5±1.0°C ($T_{db}$=36°C & 66.3±5.7% RH) |
| | 5M & 3F | | | | | | 38°C (constant $T_{db}$) | | | 183±113 | $T_w$=31.3±1.2°C ($T_{db}$=38°C & 60.8±5.4% RH) |
| | 3M & 5F | | | | | | 40°C (constant $T_{db}$) | | | 160±64 | $T_w$=30.9±0.8°C ($T_{db}$=40°C & 50.2±4.6% RH) |
| | 6M & 2F | | | | | | 12 mmHg (constant $P_a$) | | | 112±36 | $T_w$=26.2±0.7°C ($T_{db}$=50.6±1.7°C & 12.7±1.5% RH) |
| | 4M & 5F | | | | | | 16 mmHg (constant $P_a$) | | | 143±66 | $T_w$=27.8±0.4°C ($T_{db}$=47.5±2.0°C & 20.1±1.6% RH) |
| | 4M & 5F | | | | | | 20 mmHg (constant $P_a$) | | | 172±98 | $T_w$=28.6±0.5°C ($T_{db}$=44±0.2°C & 28.8±2.7% RH) |
| Cottle et al. (2022)[36] | 14 adults | 23±4 | 23.7 | 1.84±0.20 | 0.27 | MinAct (simulated daily living: cycling on ergometer at 0 W, 40-50 rpm; 90-120 min) | 34°C (constant $T_{db}$) | Ramp (30 min stabilization; $P_a$ or $T_{db}$ increased stepwise by 1 mmHg or 1 °C every 5 min) | NA | 122±60 | $T_w$=30.9°C ($T_{db}$=33.9°C & 80% RH) |
| | 14 adults | | | | | | 36°C (constant $T_{db}$) | | | 136±89 | $T_w$=30.7°C ($T_{db}$=36°C & 67% RH) |
| | 8 adults | | | | | | 38°C (constant $T_{db}$) | | | 180±105 | $T_w$=31.3°C ($T_{db}$=37.9°C & 61% RH) |
| | 9 adults | | | | | | 40°C (constant $T_{db}$) | | | 147±70 | $T_w$=30.9°C ($T_{db}$=40°C & 50% RH) |
| | 5 adults | | | | | | 42°C (constant $T_{db}$) | | | 210±118 | $T_w$=29.9°C ($T_{db}$=42°C & 39% RH) |
| | 17 adults | | | | | | 12 mmHg (constant $P_a$) | | | 142±68 | $T_w$=27.5°C ($T_{db}$=46.4°C & 21% RH) |
| | 15 adults | | | | | | 16 mmHg (constant $P_a$) | | | 160±89 | $T_w$=26.1°C ($T_{db}$=49.3°C & 14% RH) |
| | 16 adults | | | | | LightAmb (simulated slow walking: treadmill at 2.2 mph, 3% grade; 90-120 min) | 34°C (constant $T_{db}$) | Ramp (30 min stabilization; $P_a$ or $T_{db}$ increased stepwise by 1 mmHg or 1 °C every 5 min) | | 181±61 | $T_w$=27.8°C ($T_{db}$=33.8°C & 62% RH) |
| | 13 adults | | | | | | 36°C (constant $T_{db}$) | | | 205±78 | $T_w$=28.4°C ($T_{db}$=35.9°C & 55% RH) |
| | 12 adults | | | | | | 38°C (constant $T_{db}$) | | | 253±100 | $T_w$=28.4°C ($T_{db}$=38.1°C & 46% RH) |
| | 8 adults | | | | | | 40°C (constant $T_{db}$) | | | 260±93 | $T_w$=27.6°C ($T_{db}$=40.1°C & 36% RH) |
| | 5 adults | | | | | | 42°C (constant $T_{db}$) | | | 246±74 | $T_w$=27.4°C ($T_{db}$=41.9°C & 30% RH) |
| | 18 adults | | | | | | 12 mmHg (constant $P_a$) | | | 193±88 | $T_w$=26.4°C ($T_{db}$=42.3°C & 26% RH) |
| | 18 adults | | | | | | 16 mmHg (constant $P_a$) | | | 221±74 | $T_w$=24.8°C ($T_{db}$=43.7°C &19% RH) |
| Wolf et al. (2022)[37] | 4M & 5F | 24±4 | 24.0 | 1.84±0.20 | 0.27 | MinAct (simulated daily living: cycling on ergometer at 0 W, 40-50 rpm; 90-120 min) | 36°C (constant $T_{db}$) | Ramp (30 min stabilization; $P_a$ or $T_{db}$ increased stepwise by 1 mmHg or 1 °C every 5 min) | 1.34±0.27 | 117±85 | $T_w$=30.6°C ($T_{db}$=36±0.2°C & 66.4±5.4% RH) |
| | 5M & 3F | | | | | | 38°C (constant $T_{db}$) | | 1.42±0.29 | 180±105 | $T_w$=31.2°C ($T_{db}$=37.9±0.3°C & 60.5±5.1% RH) |
| | 4M & 5F | | | | | | 40°C (constant $T_{db}$) | | 1.38±0.15 | 148±70 | $T_w$=30.9°C ($T_{db}$=40±0.2°C & 50.2±4.3% RH) |

| Authors (Year) | Subject | Age (yr) | BMI (kg/m²) | BSA (m²) | $I_{cl}$ (clo) | Activity | Environmental condition | Protocol | Metabolic rate (METs) | Sweat rate (g/m²/h) | Critical environmental limit (CEL) |
|---|---|---|---|---|---|---|---|---|---|---|---|
| | 6M & 3F | | | | | | 12 mmHg (constant $P_a$) | | 1.50±0.21 | 172±98 | $T_w$=26.2°C ($T_{db}$=50.6±2.0°C & 12.7±1.5% RH) |
| | 4M & 5F | | | | | | 16 mmHg (constant $P_a$) | | 1.51±0.18 | 141±62 | $T_w$=27.8°C ($T_{db}$=47.3±1.6°C & 20.3±1.5% RH) |
| | 4M & 5F | | | | | | 20 mmHg (constant $P_a$) | | 1.40±0.20 | 104±41 | $T_w$=28.5°C ($T_{db}$=43.8±2.0°C & 29.2±2.8% RH) |
| | 4M & 5F | | | | | LightAmb (simulated slow walking: treadmill at 2.2 mph, 3% grade; 90-120 min) | 36°C (constant $T_{db}$) | Ramp (30 min stabilization; $P_a$ or $T_{db}$ increased stepwise by 1 mmHg or 1 °C every 5 min) | 2.21±0.30 | 212±69 | $T_w$=28.4°C ($T_{db}$=35.9±0.1°C & 55.5±3.9% RH) |
| | 5M & 4F | | | | | | 38°C (constant $T_{db}$) | | 2.34±0.27 | 274±107 | $T_w$=28.2°C ($T_{db}$=38±0.3°C & 45.5±2.7% RH) |
| | 4M & 5F | | | | | | 40°C (constant $T_{db}$) | | 2.30±0.23 | 263±87 | $T_w$=28°C ($T_{db}$=40.1±0.4°C & 37.3±3.3% RH) |
| | 6M & 3F | | | | | | 12 mmHg (constant $P_a$) | | 2.36±0.32 | 243±102 | $T_w$=25.2°C ($T_{db}$=44.5±2.4°C & 18.6±3.3% RH) |
| | 4M & 5F | | | | | | 16 mmHg (constant $P_a$) | | 2.31±0.25 | 206±74 | $T_w$=26.5°C ($T_{db}$=43.0±1.5°C & 24.9±2.0% RH) |
| | 4M & 5F | | | | | | 20 mmHg (constant $P_a$) | | 2.24±0.18 | 187±38 | $T_w$=27.3°C ($T_{db}$=38.9±1.0°C & 38.4±1.8% RH) |
| Wolf et al. (2023)[38] | 14 adults | 23±4 | 25±4 | 1.84±0.20 | 0.27 | MinAct (simulated daily living: cycling on ergometer at 0 W, 40-50 rpm; 90-120 min) | 34°C (constant $T_{db}$) | Ramp (30 min stabilization; $P_a$ or $T_{db}$ increased stepwise by 1 mmHg or 1 °C every 5 min) | 1.47±0.16 | 122±60 | $T_w$=30.8°C ($T_{db}$=33.9°C [33.7°C, 34.2°C] & 79.7% [77.6%, 81.7%] RH) |
| | 14 adults | | | | | | 36°C (constant $T_{db}$) | | 1.34±0.23 | 136±89 | $T_w$=30.6°C ($T_{db}$=36°C [35.9°C, 36.1°C] & 66.5% [64.1%, 69%] RH |
| | 12 adults | | | | | | 38°C (constant $T_{db}$) | | 1.41±0.23 | 202±114 | $T_w$=31.2°C (38.1°C [37.8°C, 38.4°C] & 59.5% [55.5%, 63.6%] RH) |
| | 13 adults | | | | | | 40°C (constant $T_{db}$) | | 1.38±0.15 | 167±70 | $T_w$=31.1°C ($T_{db}$=40°C [39.9°C, 40.1°C] & 51.0% [48.2%, 53.8%] RH) |
| | 17 adults | | | | | | 12 mmHg (constant $P_a$) | | 1.53±0.18 | 142±68 | $T_w$=26°C ($T_{db}$=49.3°C [48.2°C, 50.4°C] & 13.8% [12.9%, 14.7%] RH) |
| | 17 adults | | | | | | 16 mmHg (constant $P_a$) | | 1.47±0.19 | 186±110 | $T_w$=27.3°C ($T_{db}$=46.4°C [45.5°C, 47.3°C] & 20.7% [19.9%, 21.5%] RH) |
| | 12 older adults | 71±6 | 26±5 | 1.82±0.23 | 0.27 | MinAct (simulated daily living: cycling on ergometer at 0 W, 40-50 rpm; 90-120 min) | 34°C (constant $T_{db}$) | Ramp (30 min stabilization; $P_a$ or $T_{db}$ increased stepwise by 1 mmHg or 1 °C every 5 min) | 1.30±0.16 | 77±45 | $T_w$=27.9°C ($T_{db}$=33.9°C [33.7°C, 34.0°C] & 62.1% [52.8%, 71.4%] RH) |
| | 13 older adults | | | | | | 36°C (constant $T_{db}$) | | 1.39±0.25 | 84±50 | $T_w$=28.3°C ($T_{db}$=36.2°C [36.0°C, 36.4°C] & 53.5% [48.2%, 58.8%] RH) |
| | 14 older adults | | | | | | 38°C (constant $T_{db}$) | | 1.42±0.30 | 89±57 | $T_w$=28.3°C ($T_{db}$=38.0°C [37.8°C, 38.3°C] & 46.1% [41.7%, 50.6%] RH) |

| Authors (Year) | Subject | Age (yr) | BMI (kg/m²) | BSA (m²) | $I_{cl}$ (clo) | Activity | Environmental condition | Protocol | Metabolic rate (METs) | Sweat rate (g/m²/h) | Critical environmental limit (CEL) |
|---|---|---|---|---|---|---|---|---|---|---|---|
| | 14 older adults | | | | | | 40°C (constant $T_{db}$) | | 1.45±0.26 | 90±50 | $T_w$=27.2°C ($T_{db}$=40.2°C [40.0°C, 40.3°C] & 34.0% [29.2%, 38.8%] RH) |
| | 10 older adults | | | | | | 12 mmHg (constant $P_a$) | | 1.34±0.17 | 92±62 | $T_w$=24.2°C ($T_{db}$=42.9°C [40.9°C, 44.8°C] & 18.8% [17.0%, 20.5%] RH) |
| | 16 older adults | | | | | | 16 mmHg (constant $P_a$) | | 1.35±0.27 | 83±70 | $T_w$=25.7°C ($T_{db}$=40.3°C [38.4°C, 42.2°C] & 28.6% [25.8%, 31.5%] RH) |
| | 12 older adults | 65-77 | 21-31 | 1.56-2.05 | 0.27 | Resting | 34°C (constant $T_{db}$) | Ramp (30 min stabilization; $P_a$ or $T_{db}$ increased stepwise by 1 mmHg or 1 °C every 5 min) | 0.83±0.16 | 45±21 | $T_w$=28.6°C ($T_{db}$=33.9°C [33.7°C, 34.1°C] & 66.5% [55.9%, 77.1%] RH) |
| | 14 older adults | | | | | | 36°C (constant $T_{db}$) | | 0.80±0.13 | 58±40 | $T_w$=29.4°C ($T_{db}$=36.2°C [35.9°C, 36.4°C] & 59.1% [54.6%, 63.5%] RH) |
| | 14 older adults | | | | | | 38°C (constant $T_{db}$) | | 0.86±0.17 | 68±58 | $T_w$=30°C ($T_{db}$=38.1°C [37.9°C, 38.3°C] & 53.4% [48.4%, 58.5%] RH) |
| | 14 older adults | | | | | | 40°C (constant $T_{db}$) | | 0.86±0.14 | 66±56 | $T_w$=28.4°C ($T_{db}$=40.1°C [40.0°C, 40.1°C] & 38.8% [34.5%, 43.1%] RH) |
| | 13 older adults | | | | | | 12 mmHg (constant $P_a$) | | 0.89±0.16 | 52±31 | $T_w$=24.9°C ($T_{db}$=46.1°C [44.1°C, 48.1°C] & 15.6% [13.7%, 17.5%] RH) |
| | 15 older adults | | | | | | 16 mmHg (constant $P_a$) | | 0.90±0.16 | 52±25 | $T_w$=27°C ($T_{db}$=44.1°C [41.8°C, 46.4°C] & 24.0% [21.2%, 26.8%] RH) |
| Wang et al. (2023)[40] | 4M | 25±0.8 | 22.6±0.8 | 1.73±0.05 | 0.40 | Seated, office work | 40±0.5°C & 20±5% RH 1.0±0.1 m/s air velocity | Prolonged 14-hour exposure, passive heating | NA | NA | $T_w$=20.8-24.4°C compensable (first 9 h); uncompensable from 9–14 h due to dehydration (limited fluid intake) |
| Meade et al. (2025)[39] | 9M & 3F | 25-32 | 22.9-30.4 | 1.8-2.2 | NA | Rest (reclined semi-supine position, no activity) | 42°C & 28% RH | Ramp (42°C & 28% RH for 70 min; RH increased stepwise by 3% every 10 min until reaching 70%); $T_w$=33.9°C (42°C & 56%RH); & $T_w$=31.2°C (42°C & 44% RH) | NA | $T_w$=33.5°C: 40.5 [8.5,72.5]; $T_w$=30.7°C: 133 [101,165]* | $T_w$=33.9°C ($T_{db}$=42°C & 56% RH): 10-h to reach heatstroke $T_{core}$=40.2°C; $T_w$=31.2°C (42°C & 44% RH): >24-h to reach heat stroke $T_{core}$=40.2°C. |
| Wang et al. (2025)[24] | 20M | 22.4±0.9 | 22.3±2.1 | 1.83±0.11 | 0.40 | Seated, light office work (reading, computer, tablet, smartphone) | 36.1±0.1°C & 74.1±1.2% RH 39.9±0.3°C & 57.7±1.7% RH 44.0±0.1°C & 29.2±1.2% RH 47.1±0.1°C & 35.8±1.5% RH | Prolonged 8-hour exposure, passive heating | 1.47-1.53 | NA | Compensable at $T_w$=28.7°C, 31.4°C, 32°C, & 33°C |

| Authors (Year) | Subject | Age (yr) | BMI (kg/m²) | BSA (m²) | $I_{cl}$ (clo) | Activity | Environmental condition | Protocol | Metabolic rate (METs) | Sweat rate (g/m²/h) | Critical environmental limit (CEL) |
|---|---|---|---|---|---|---|---|---|---|---|---|
| | | | | | | | 50.1±0.2°C & 24.0±1.3% RH | | | | |
| | 16F | 20.9±2 | 21.1±2.2 | 1.60±0.12 | 0.41 | Seated, light office work (reading, computer, tablet, smartphone) | 36.1±0.1°C & 74.1±1.2% RH<br>39.9±0.3°C & 57.7±1.7% RH<br>44.0±0.1°C & 29.2±1.2% RH<br>47.1±0.1°C & 35.8±1.5% RH<br>50.1±0.2°C & 24.0±1.3% RH | Prolonged 8-hour exposure, passive heating | 1.41-1.43 | NA | Compensable at $T_w$=28.7°C, 31.4°C, 32°C, & 33°C |
| This study (2025) | 20M | 24.6±4.0 | 22.5±1.9 | 1.85±0.09 | 0.40 | Seated, light office work (reading, device use) | 36.1±0.1°C & 74.1±1.2% RH<br>39.9±0.5°C & 57.7±2% RH<br>51.9±0.4°C & 24.2±0.7% RH | Prolonged 8-hour exposure, passive heating | NA | 153±38 | $T_w$=32°C: compensable (males required >34 h to reach $T_{core}$=40.5°C) |
| | 16F | 23.0±2.2 | 20.9±1.4 | 1.64±0.08 | 0.41 | Seated, light office work (reading, device use) | 36.1±0.1°C & 74.1±1.2% RH<br>39.9±0.5°C & 57.7±2% RH<br>51.9±0.4°C & 24.2±0.7% RH | Prolonged 8-hour exposure, passive heating | NA | 145±45 | $T_w$=32°C: compensable (females required >35 h to reach $T_{core}$=40.5°C) |
| | 20M | 24.6±4.0 | 22.5±1.9 | 1.85±0.09 | 0.40 | Seated, light office work (reading, device use) | 37.3±0.1°C & 75.3±0.5% RH<br>42.3±0.1°C & 51.5±0.6% RH<br>47.1±0.2°C & 35.2±0.7% RH | Prolonged 8-hour exposure, passive heating | 1.51±0.30 | 107±59<br>14952<br>165±71 | $T_w$=33°C: compensable (males required >33 h to reach $T_{core}$=40.5°C) |
| | 16F | 23.0±2.2 | 20.9±1.4 | 1.64±0.08 | 0.41 | Seated, light office work (reading, device use) | 37.3±0.1°C & 75.3±0.5% RH<br>42.3±0.1°C & 51.5±0.6% RH<br>47.1±0.2°C & 35.2±0.7% RH | Prolonged 8-hour exposure, passive heating | 1.39±0.17 | 107±62<br>151±38<br>187±32 | $T_w$=33°C: compensable (females required >35 h to reach $T_{core}$=40.5°C) |
| | 20M | 24.6±4.0 | 22.5±1.9 | 1.85±0.09 | 0.40 | Seated, light office work (reading, device use) | 38.3±0.1°C & 75.2±0.5% RH<br>43.3±0.2°C & 52.1±0.5% RH<br>48.1±0.2°C & 35.9±0.6% RH | Prolonged 8-hour exposure, passive heating | NA | 173±41<br>203±40<br>197±74 | $T_w$=34°C: uncompensable (males required 12.5-12.9 h to reach $T_{core}$=40.5°C) |
| | 16F | 23.0±2.2 | 20.9±1.4 | 1.64±0.08 | 0.41 | Seated, light office work (reading, device use) | 38.3±0.1°C & 75.2±0.5% RH<br>43.3±0.2°C & 52.1±0.5% RH | Prolonged 8-hour exposure, passive heating | NA | 133±40<br>187±23<br>223±62 | $T_w$=34°C: uncompensable (females required 15.5-16.2 h to reach $T_{core}$=40.5°C) |

| Authors (Year) | Subject | Age (yr) | BMI (kg/m²) | BSA (m²) | $I_{cl}$ (clo) | Activity | Environmental condition | Protocol | Metabolic rate (METs) | Sweat rate (g/m²/h) | Critical environmental limit (CEL) |
|---|---|---|---|---|---|---|---|---|---|---|---|
| | | | | | | | 48.1±0.2°C & 35.9±0.6% RH | | | | |
| | 20M | 24.6±4.0 | 22.5±1.9 | 1.85±0.09 | 0.40 | Seated, light office work (reading, device use) | 50.2±0.6°C & 35.0±1.3% RH | Prolonged exposure, passive heating | NA | 238±71 | $T_w$=35°C: uncompensable (males required 7.1-7.7 h to reach $T_{core}$=40.5°C) |
| | | | | | | | 45.3±0.4°C & 50.3±1.3% RH | | | 254±74 | |
| | | | | | | | 40.6±0.3°C & 69.5±1.4% RH | | | 239±76 | |
| | 16F | 23.0±2.2 | 20.9±1.4 | 1.64±0.08 | 0.41 | Seated, light office work (reading, device use) | 50.2±0.6°C & 35.0±1.3% RH | Prolonged exposure, passive heating | NA | 282±106 | $T_w$=35°C: uncompensable (females required 8.3-8.6 h to reach $T_{core}$=40.5°C) |
| | | | | | | | 45.3±0.4°C & 50.3±1.3% RH | | | 241±91 | |
| | | | | | | | 40.6±0.3°C & 69.5±1.4% RH | | | 264±104 | |
| **Biophysical or thermoregulatory modeling** | | | | | | | | | | | |
| **Authors (Year)** | **Subject** | **Age (yr)** | **BMI (kg/m²)** | **BSA (m²)** | **$I_{cl}$ (clo)** | **Activity** | **Condition ($T_{db}$ & RH)** | **Protocol** | **Metabolic rate (METs)** | **Sweat rate (g/m²/h)** | **Critical environmental limit (CEL)** |
| Vanos et al. (2023)[18] | Young adults | NA | NA | NA | 0.36 | Resting (seated, awake) | 25-60°C | Biophysical modelling ($T_{core}$=43°C reached after 6-h exposure) | 1.5 | NA | $T_w$=26.1°C ($T_{db}$=53.2°C &10% RH) |
| | | | | | | | | | | | $T_w$=31.5°C ($T_{db}$=49.9°C & 25% RH) |
| | | | | | | | | | | | $T_w$=33.7°C ($T_{db}$=43.3°C & 50% RH) |
| | | | | | | | | | | | $T_w$=33.9°C ($T_{db}$=38°C & 75% RH) |
| | | | | | | | | | | | $T_w$=34.1°C ($T_{db}$=35.6°C & 90% RH) |
| | | | | | | | | | | | $T_w$=34.4°C ($T_{db}$=34.3°C & 100% RH) |
| | Elderly | NA | NA | NA | 0.36 | Resting (seated, awake) | 25-60°C | Biophysical modelling ($T_{core}$=43°C reached after 6-h exposure) | 1.5 | NA | $T_w$=22.2°C ($T_{db}$=46.4°C & 10% RH) |
| | | | | | | | | | | | $T_w$=28.3°C ($T_{db}$=45.4°C & 25% RH) |
| | | | | | | | | | | | $T_w$=32.7°C ($T_{db}$=42.1°C & 50% RH) |
| | | | | | | | | | | | $T_w$=33.4°C ($T_{db}$=37.4°C & 75% RH) |
| | | | | | | | | | | | $T_w$=33.7°C ($T_{db}$=35.2°C & 90% RH) |
| | | | | | | | | | | | $T_w$=34.1°C ($T_{db}$=34°C & 100% RH) |
| Xu et al. (2024)[19] | Standard person | NA | 27.0 | 2.06 | 0.27 | Minimal activity | 28-50°C & 5-95% RH | Thermoregulatory model (critical environment limits defined as conditions where $T_{core}$ reaches 38°C within 6 h) | 1.3 | 61.4 | $T_w$=33.3°C ($T_{db}$=34°C & 95% RH) |
| | | | | | | | | | | 79.1 | $T_w$=32.9°C ($T_{db}$=36°C & 80% RH) |
| | | | | | | | | | | 99.5 | $T_w$=32.1°C ($T_{db}$=38°C & 65% RH) |
| | | | | | | | | | | 110.7 | $T_w$=32.0°C ($T_{db}$=40°C & 55% RH) |
| | | | | | | | | | | 124.7 | $T_w$=31.4°C ($T_{db}$=42°C & 45% RH) |
| | | | | | | | | | | 134.9 | $T_w$=30.4°C ($T_{db}$=44°C & 35% RH) |
| | | | | | | | | | | 147.0 | $T_w$=28.7°C ($T_{db}$=46°C & 25% RH) |

**Supplementary Table S2. Baseline characteristics of participants included in the study.** Data are presented as mean ± SD (standard deviation). Participants self-reported their activity levels as high, moderate, or low based on the types and intensity of activities performed. High activity included gym workouts. Moderate activity included running, gym workouts, strength training, basketball, and badminton. Low activity included jogging, gym workouts, playing badminton, cycling, climbing, and yoga. Prior to participating in the heat exposure trials, baseline complete blood count (CBC), plasma electrolyte concentrations, and fasting plasma glucose levels were assessed from venous blood samples at a local hospital to determine participant eligibility. The CBC was analyzed using an automated hematology analyzer, while plasma electrolytes and glucose were measured using a biochemistry analyzer, in accordance with standard venipuncture and sample processing protocols. Specific eligibility markers included hemoglobin, hematocrit, total leukocyte and neutrophil counts, sodium, potassium, chloride, and glucose. Inclusion thresholds were set to ensure participants had adequate oxygen-carrying capacity, were fully hydrated, and had no signs of systemic inflammation. Hemoglobin concentrations were required to fall between 130 and 165 g/L for males and 115 and 150 g/L for females, with corresponding hematocrit values of 38–50% and 35–45%, respectively. Total leukocyte counts had to be within 4.0–10.0 × $10^9$/L, and neutrophil counts between 2.0 and 6.0 × $10^9$/L, indicating the absence of acute or underlying inflammation. Electrolyte inclusion ranges were set at 135–145 mmol/L for sodium, 3.5–5.0 mmol/L for potassium, and 98–110 mmol/L for chloride. Fasting plasma glucose concentrations were required to fall within 65–139 mg/dL. All enrolled participants met these criteria, confirming that they were healthy, euhydrated, and physiologically suitable for prolonged heat exposure without increased risk of heat-related illness or confounding pre-existing inflammation.

| **Participant characteristics** | **Females (n=16)** | **Males (n=20)** |
|---|---|---|
| | **Mean ± SD** | **Mean ± SD** |
| Age (years) | 23.0±2.2 | 25.1±3.4 |
| Height (cm) | 166±4 | 175±5 |
| Body mass (kg) | 57.5±4.9 | 70.6±6.4 |
| Body surface area ($m^2$) | 1.64±0.08 | 1.85±0.09 |
| Body mass index (kg/$m^2$) | 20.79±1.43 | 22.98±1.98 |
| Resting heart rate (beats/min) | 77±8 | 72±10 |
| Self-reported activity level | 6.25% (high), 25% (moderate), 68.75% (low) | 65% (moderate), 35% (low) |
| Exercise frequency (days/week) | 2.0±2.0 | 2.6±1.4 |
| Leukocyte counts ($10^9$/L) | 6.65±1.21 | 5.83±1.12 |
| Erythrocyte counts ($10^{12}$/L) | 4.10±0.91 | 5.26±0.29 |
| Hemoglobin (g/L) | 130.9±18.9 | 160.6±7.3 |
| Platelets ($10^9$/L) | 206.2±55.4 | 183.4±22.0 |
| Neutrophils ($10^9$/L) | 2.89±0.94 | 2.55±1.11 |
| Lymphocytes ($10^9$/L) | 3.27±0.91 | 2.90±0.60 |
| Plasma $K^+$ level (mmol/L) | 3.79±0.32 | 4.14±0.24 |
| Plasma $Na^+$ level (mmol/L) | 141.1±5.7 | 143.8±0.9 |
| Plasma $Cl^-$ level (mmol/L) | 99.5±3.7 | 102.7±0.8 |
| Glucose (mg/dL) | 82±11 | 91±9 |

**Supplementary Table S3. Skin temperature, heart rate, mean arterial pressure, urine specific gravity, sweat production, sweat rate, and perceptual responses (ratings of thermal discomfort, thirst sensation, and psychological stress) in young adult females (n=16) and males (n=20) during exposure to 12 dry-bulb temperature ($T_{db}$) and relative humidity (RH) combinations corresponding to wet-bulb temperatures ($T_w$) of 32-35°C.** MST, mean skin temperature, °C; HR, heart rate, beats per minute (bpm); MAP, mean arterial pressure, mmHg; SW, total sweat production, g; SR, sweat rate, g/h; $T_{db}$, dry-bulb temperature, °C; -, not applicable; TD, thermal discomfort; TS, thirst sensation; PS, psychological stress.

| Indicators | | $T_w$=32°C | | | $T_w$=33°C | | | $T_w$=34°C | | | $T_w$=35°C | | |
|---|---|---|---|---|---|---|---|---|---|---|---|---|---|
| | | $T_{db}$=36°C, 74.5%RH | $T_{db}$=40°C, 55%RH | $T_{db}$=52°C, 22.5%RH | $T_{db}$=37°C, 74.8%RH | $T_{db}$=42°C, 51.5%RH | $T_{db}$=47°C, 35.5%RH | $T_{db}$=38°C, 75.1%RH | $T_{db}$=43°C, 52.1%RH | $T_{db}$=48°C, 36.2%RH | $T_{db}$=40°C, 70.3%RH | $T_{db}$=45°C, 49%RH | $T_{db}$=50°C, 34.4%RH |
| **Females** | | | | | | | | | | | | | |
| MST | Pre | 33.7±0.5 | 33.3±0.2 | 33.1±0.5 | 32.9±0.3 | 32.7±0.4 | 33.5±0.4 | 32.9±0.4 | 33.4±0.4 | 33.2±0.4 | 33.1±0.5 | 33.7±0.4 | 32.8±0.6 |
| | Post | 36.9±0.2 | 37.3±0.4 | 39.3±0.3 | 37.1±0.4 | 37.7±0.6 | 38.6±0.3 | 37.3±0.4 | 38.1±0.3 | 38.8±0.4 | 37.9±0.4 | 38.4±0.4 | 38.9±0.3 |
| HR | Pre | 78±6 | 73±4 | 76±5 | 74±4 | 73±6 | 75±4 | 76±5 | 74±6 | 75±5 | 71±4 | 72±5 | 73±4 |
| | Post | 108±13 | 110±17 | 123±11 | 112±11 | 116±15 | 120±13 | 119±13 | 118±11 | 124±15 | 129±9 | 128±12 | 130±9 |
| MAP | Pre | 80.1±7.6 | 80.1±4.6 | 79.2±6.1 | 82.9±4.9 | 79.9±6.0 | 83.2±5.2 | 81.4±5.6 | 80.9±6.0 | 79.5±4.6 | 83.3±4.9 | 85.1±6.2 | 83.2±4.5 |
| | Post | 68.4±6.0 | 73.1±4.8 | 68.7±5.7 | 71.2±3.1 | 69.0±6.6 | 70.5±2.7 | 70.8±3.2 | 70.3±3.3 | 71.5±3.0 | 73.4±3.1 | 75.0±3.2 | 73.3±3.3 |
| USG | Pre | 1.008±0.004 | 1.012±0.001 | 1.014±0.005 | 1.014±0.003 | 1.009±0.004 | 1.011±0.003 | 1.011±0.003 | 1.013±0.004 | 1.013±0.005 | 1.015±0.002 | 1.014±0.002 | 1.014±0.005 |
| | Post | 1.005±0.006 | 1.007±0.00 | 1.004±0.003 | 1.012±0.004 | 1.014±0.004 | 1.015±0.005 | 1.007±0.004 | 1.009±0.007 | 1.011±0.005 | 1.008±0.003 | 1.008±0.002 | 1.004±0.003 |
| SW | - | 1327±326 | 1844±582 | 2502±642 | 1598±496 | 1886±499 | 2255±376 | 1379±526 | 1546±507 | 1881±518 | 1676±640 | 1561±296 | 1756±634 |
| SR | - | 138±74 | 223±53 | 313±80 | 202±52 | 236±62 | 282±47 | 209±68 | 256±81 | 357±90 | 428±165 | 410±87 | 455±167 |
| Perceptual responses | | | | | | | | | | | | | |
| TC | Pre | -0.20±0.63 | 0.29±0.49 | 0.18±0.40 | -0.09±0.30 | 0.00±0.45 | -0.17±0.58 | -0.08±0.29 | 0.27±0.47 | 0.18±0.40 | -0.45±0.69 | 0.10±0.57 | -0.73±0.79 |
| | Post | -1.90±1.20 | -2.86±0.69 | -2.19±0.98 | -2.27±1.10 | -2.55±0.82 | -2.58±0.67 | -2.58±0.90 | -2.18±0.75 | -3.00±0.63 | -2.45±0.52 | -2.30±0.67 | -2.45±0.69 |
| TS | Pre | 1.10±0.32 | 1.71±0.76 | 1.36±0.50 | 1.71±0.49 | 1.91±0.30 | 1.83±0.39 | 1.58±0.51 | 1.45±0.52 | 1.36±0.50 | 1.22±0.44 | 1.13±0.35 | 1.18±0.40 |
| | Post | 2.60±1.58 | 1.86±0.69 | 1.73±0.47 | 1.86±0.38 | 1.82±0.40 | 1.92±0.29 | 1.92±0.29 | 1.82±0.40 | 1.64±0.50 | 1.89±0.33 | 1.75±0.46 | 1.91±0.30 |
| PS | Pre | 0.50±0.85 | 0.86±1.21 | 0.55±0.52 | 0.13±0.35 | 0.18±0.40 | 0.25±0.45 | 0.17±0.39 | 0.27±0.47 | 0.45±0.52 | 0.09±0.30 | 0.00±0.00 | 0.09±0.30 |
| | Post | 3.30±3.40 | 5.57±1.27 | 4.55±2.38 | 4.38±1.41 | 4.27±1.42 | 4.83±1.80 | 6.00±1.76 | 4.82±1.72 | 6.18±1.17 | 6.27±1.01 | 6.10±1.29 | 6.73±1.27 |
| **Males** | | | | | | | | | | | | | |
| MST | Pre | 33.6±0.5 | 32.9±0.3 | 33.4±0.5 | 32.9±0.4 | 33.8±0.4 | 33.5±0.4 | 33.3±0.5 | 33.1±0.4 | 33.4±0.5 | 33.0±0.4 | 33.3±0.6 | 32.9±0.5 |
| | Post | 37.0±0.2 | 37.3±0.6 | 39.1±0.6 | 37.0±0.4 | 37.6±0.4 | 38.5±0.5 | 37.2±0.3 | 38.1±0.4 | 38.6±0.6 | 37.9±0.4 | 38.3±0.3 | 38.8±0.4 |

| | | | | | | | | | | | | | |
|---|---|---|---|---|---|---|---|---|---|---|---|---|---|
| HR | Pre | 76±6 | 73±6 | 76±4 | 77±6 | 74±5 | 74±4 | 75±5 | 76±6 | 76±4 | 75±6 | 74±6 | 76±4 |
| | Post | 104±14 | 110±17 | 115±15 | 112±10 | 113±9 | 117±10 | 114±10 | 120±13 | 121±15 | 120±11 | 120±12 | 122±10 |
| MAP | Pre | 80.9±4.2 | 87.3±5.8 | 86.9±4.6 | 86.4±2.5 | 85.8±4.8 | 86.3±3.5 | 85.1±3.9 | 83.9±5.6 | 83.6±3.9 | 83.4±3.8 | 83.7±5.5 | 81.8±5.2 |
| | Post | 71.5±4.5 | 80.2±5.8 | 73.8±5.8 | 74.2±1.9 | 75.9±2.3 | 76.3±2.6 | 74.9±3.1 | 73.5±4.1 | 73.2±3.4 | 77.0±2.8 | 75.9±5.0 | 77.4±4.7 |
| USG | Pre | 1.008±0.005 | 1.016±0.006 | 1.014±0.004 | 1.011±0.004 | 1.010±0.004 | 1.012±0.003 | 1.014±0.004 | 1.013±0.003 | 1.013±0.004 | 1.017±0.002 | 1.015±0.003 | 1.016±0.003 |
| | Post | 1.007±0.009 | 1.014±0.010 | 1.011±0.005 | 1.015±0.005 | 1.016±0.006 | 1.020±0.005 | 1.009±0.004 | 1.010±0.005 | 1.011±0.007 | 1.008±0.003 | 1.010±0.003 | 1.012±0.004 |
| SW | - | 1580±428 | 2222±630 | 2947±705 | 1846±544 | 2365±367 | 2288±294 | 1854±482 | 2242±512 | 2109±649 | 1767±740 | 1834±690 | 1697±674 |
| SR | - | 195±44 | 278±79 | 368±88 | 231±68 | 296±46 | 346±57 | 316±79 | 365±75 | 368±132 | 437±137 | 462±126 | 435±131 |
| | | | | | | | Perceptual responses | | | | | | |
| TD | Pre | -0.25±0.62 | -0.71±0.56 | 0.17±0.41 | 0.27±0.47 | 0.08±0.29 | -0.08±0.67 | 0.09±0.30 | 0.18±0.60 | 0.25±0.45 | 0.08±0.29 | -0.20±0.42 | -0.42±0.51 |
| | Post | -2.64±0.67 | -2.17±0.78 | -2.67±1.03 | -2.18±0.98 | -2.08±0.90 | -2.42±0.90 | -2.64±1.29 | -2.55±1.13 | -2.75±0.87 | -1.92±0.67 | -2.80±0.92 | -2.42±0.67 |
| TS | Pre | 1.17±0.39 | 2.76±1.45 | 1.33±0.52 | 1.64±0.50 | 1.58±0.51 | 1.58±0.67 | 1.09±0.54 | 1.18±0.40 | 1.42±0.51 | 1.25±0.45 | 1.30±0.48 | 1.33±0.49 |
| | Post | 2.25±1.36 | 3.17±1.18 | 1.50±0.55 | 1.91±0.30 | 1.75±0.45 | 1.83±0.39 | 1.55+0.52 | 1.73±0.47 | 1.75±0.45 | 1.83±0.39 | 1.80±0.42 | 1.75±0.45 |
| PS | Pre | 0.58±0.67 | 0.44±0.53 | 0.33±0.52 | 0.36±0.50 | 0.17±0.39 | 0.08±0.29 | 0.55±0.52 | 0.64±0.50 | 0.33±0.49 | 0.33±0.65 | 0.60±0.97 | 1.08±0.90 |
| | Post | 4.73±2.69 | 4.56±1.81 | 5.33±2.50 | 4.45±1.57 | 4.75±1.22 | 5.42±1.51 | 6.00±2.14 | 6.45±1.04 | 5.25±2.09 | 5.50±1.00 | 5.90±2.02 | 6.00±1.13 |

### Supplementary video legends

**Supplementary Video S1.** Females and male participants periodically wipe sweat from their faces, consume lunch at 12:00 p.m., drink fluids, and undergo blood pressure measurements during prolonged exposure (up to 8 hours) at $T_w$=34°C ($T_{db}$=43°C, RH=52.1%).

**Supplementary Video S2.** Experimenters perform metabolic rate measurements on a female participant during prolonged exposure at $T_w$=33°C ($T_{db}$=42°C, RH=51.5%).
